# A computational model of dopaminergic modulation of hippocampal Schaffer collateral-CA1 long-term plasticity

**DOI:** 10.1101/2021.01.27.428522

**Authors:** Joseph Schmalz, Gautam Kumar

## Abstract

Dopamine plays a critical role in modulating the long-term synaptic plasticity of the hippocampal Schaffer collateral-CA1 pyramidal neuron synapses (SC-CA1), a widely accepted cellular model of learning and memory. Limited results from hippocampal slice experiments over the last four decades have shown that the timing of the activation of dopamine D1/D5 receptors relative to a high/low-frequency stimulation (HFS/LFS) in SC-CA1 synapses regulates the modulation of HFS/LFS-induced long-term potentiation/depression (LTP/LTD) in these synapses. However, the existing literature lacks a complete picture of how various concentrations of D1/D5 agonists and the relative timing between the activation of D1/D5 receptors and LTP/LTD induction by HFS/LFS, affect the spatiotemporal modulation of SC-CA1 synaptic dynamics. In this paper, we have developed a computational model, a first of its kind, to make quantitative predictions of the temporal dose-dependent modulation of the HFS/LFS induced LTP/LTD in SC-CA1 synapses by D1/D5 agonists activating cAMP-mediating biochemical pathways. Our model combines the biochemical effects with the electrical effects at the electrophysiological level. We have estimated the model parameters from the published electrophysiological data, available from diverse hippocampal CA1 slice experiments, in a Bayesian framework. Our modeling results demonstrate the capability of our model in making quantitative predictions of the available experimental results under diverse HFS/LFS protocols. The predictions from our model show a strong nonlinear dependency of the modulated LTP/LTD by D1/D5 agonists on the relative timing between the activated D1/D5 receptors and the HFS/LFS protocol as well as the applied concentration of D1/D5 agonists. Particularly, our model predicts that D1/D5 agonists could significantly boost the LTP induced by weak HFS if the agonist is applied much before the HFS protocol. Furthermore, our model predicts that specific D1/D5 agonists can convert the LFS-induced LTD in SC-CA1 synapses to LTP if D1/D5 receptors are activated before the applied LFS protocol.

**Author summary:** Dopamine, a reward neuromodulator, plays an essential role in shaping hippocampal-dependent learning and memory of behavioral tasks. Limited experimental studies have revealed that pharmacological agents of dopaminergic receptors can significantly modulate the electrically-induced long-term potentiation/depression (LTP/LTD) of the hippocampal Schaffer collateral CA1 pyramidal (SC-CA1) synapses, a cellular model of learning and memory, in a time and dose dependent manner.

However, exploring the effect of the parameter space of various concentration levels of the applied pharmacological agent as well as the frequency-specific characteristics of the high (low) frequency stimulation (H(L)FS) protocol on the dopaminergic receptors’ mediated spatiotemporal modulation of LTP/LTD is a combinatorically challenging problem which is both expensive and time-consuming to address in experiments alone. Here, we develop a multi-timescale computational modeling framework to address this question. Our model integrates the slow biochemical dynamics and the fast-electrical dynamics of the CA1 pyramidal neuron and makes quantitative predictions of the experimentally observed modulation of H(L)FS-induced LTP/LTD in SC-CA1 synapses by dopaminergic (D1/D5) receptors agonists. Our modeling results complement the experimental findings and show specific predictions on the potential role of dopamine in strengthening weak synapses.

## Introduction

Dopamine acts as a critical neuromodulator involved in significantly modulating the hippocampal Schaffer collateral CA1 pyramidal (SC-CA1) synaptic plasticity in a dose-dependent manner and thus plays an essential role in shaping hippocampal-dependent learning and memory [1–4]. In the last four decades, diverse experimental protocols used to induce long-term potentiation (LTP) have revealed molecular signaling pathways underlying the modulation of SC-CA1 LTP by appropriate pharmacological agents activating dopaminergic D1/D5 receptors [1, 5–8]. One of the significant findings from the various experimental studies is that the timing of the activation of D1/D5 receptors by a pharmacological agent relative to the high-frequency stimulation (HFS) induced LTP protocol determines the modulation of SC-CA1 LTP by the pharmacological activation of D1/D5 receptors [2]. Another significant result showed from these experimental studies is the dose-dependent modulation of SC-CA1 LTP by appropriate pharmacological agents [4]. Despite all these advances, our current understanding of how dopamine modulates the HFS-induced synaptic integrity and plasticity of SC-CA1 synapses is still incomplete. For example, it is not clear from the existing literature how much LTP modulation will occur if one activates the D1/D5 receptors at a time in between the two extreme timings relative to the HFS. Moreover, how do these modulations vary on the temporal scale with different concentration levels of the applied pharmacological agents or a specific HFS protocol? In this paper, we attempt to provide essential insights to fill in the gaps of our current understanding on the D1/D5 receptors’ mediated modulation of SC-CA1 synaptic long-term potentiation/depression (LTP/LTD) by developing a multi timescale computational modeling framework, for the first time, that predicts the D1/D5 receptors mediated spatiotemporal modulation of high/low frequency stimulation (HFS/LFS) induced LTP/LTD at the electrophysiological level.

Exploring the effect of the parameter space of various concentration levels of the applied pharmacological agent as well as the frequency-specific characteristics of the HFS/LFS protocol on the D1/D5 receptors mediated spatiotemporal modulation of HFS/LFS-induced LTP/LTD is a combinatorically challenging problem which is both expensive and time-consuming to address in experiments alone. A computational modeling approach integrating experimental findings at various levels (such as molecular and cellular levels) and from diverse experimental protocols in a biophysiological manner could potentially address this challenge. In this regard, computational modeling approaches have been developed to explain molecular level mechanisms underlying LTP/LTD [9–16] and modulation of the neural activity by the activation of G-protein coupled receptors, such as dopamine receptors [17–20], muscarinic receptors [21, 22], and *β*-adrenergic receptors [23, 24]. Furthermore, unified multi-scale models have been developed that combine detailed biochemical molecular level dynamics with electrical dynamics in medium spiny neurons [25] and CA1 hippocampal neurons [26]. Detailed molecular-level modeling of chemical signaling pathways in these works explained the published biochemical data from several diverse experimental protocols on LTP induction in the CA1 pyramidal neurons. Moreover, it provided a comprehensive understanding of critical molecular mechanisms involved in the modulation of this LTP by dopamine. Despite these initial modeling efforts, the literature still lacks a unified modeling approach that integrates biochemical effects on electrophysiology to systematically investigate the spatiotemporal modulation of HFS/LFS-induced LTP/LTD of the SC-CA1 synapses by the activation of D1/D5 receptors under various parametric conditions.

In this paper, we have developed a computational modeling approach to integrate the spatiotemporal impact of D1/D5 agonists on the HFS/LFS-induced early and late LTP/LTD at the electrophysiological level. Our modeling hypothesis is that the chain of biochemical signaling initiated by HFS/LFS and D1/D5 receptors agonists compete for a limited available biochemical resources to induce and/or modulate late-LTP/LTD in the hippocampal SC-CA1 synapses. We have formulated our hypotheses based on the available classical experimental results on the modulation of the HFS-induced LTP by a D1/D5 agonist SKF 38393, where the authors showed that the application of SKF 38393 more than 200 minutes before the HFS protocol initially enhances the HFS-induced LTP but after 3 hours the dopaminergic enhancement decays back to the LTP level induced by only HFS [2]. In contrast, the application of SKF 38393 50 minutes after the HFS protocol does not affect the HFS-induced LTP [2]. Additionally, it has been shown experimentally that D1/D5 receptor modulation of LFS-induced LTD due to the release of SKF 38393 immediately after the LFS protocol disappears when the same amount of SKF 38393 is released 60 minutes after the LFS protocol [27]. These experiments highlight a reduced efficacy of either the D1/D5 receptors agonist or HFS/LFS protocol when it arrives as the second input. Based on these experimental results, we hypothesize that the first input may use up the shared biochemical resources rendering the second input less effective.

Using the hypotheses mentioned above, we have developed a set of phenomenological models to describe the temporal dose-dependent effect of dopamine D1/D5 receptors agonist on the maximum synaptic conductance using published electrophysiological data from hippocampal CA1 slice experiments on the % change in field excitatory postsynaptic potential (fEPSP) slope in response to D1/D5 agonists SKF 38393, 6-bromo-APB, and dopamine. Since the fEPSP slope is an extracellular EPSP recording from the stratum radiatum of the CA1 area, we have used the recent data on the simultaneous recording of fEPSP slope, and intracellular EPSP slope [28] to develop a linear correlation and used it to transform the synaptic EPSP from CA1 neuron model into fEPSP. To model the synaptically evoked EPSP, we have used an experimentally validated single compartmental biophysiological model in a Hodgkin-Huxley formalism from [29] to represent the CA1 neuron dynamics. The synaptic dynamics, as well as the dynamics of high/low-frequency stimulation (HFS/LFS), induced LTP/LTD of SC-CA1 synapse are described using published phenomenological models of SC-CA1 synaptic dynamics and LTP/LTD [30–32]. We have used an approximate Bayesian computation method with a sequential Monte Carlo scheme [33] to estimate model parameters from the available electrophysiological data in the literature.

## Results

### The relative time between D1/D5 receptors activation and HFS significantly impact the temporal modulation of HFS-induced LTP in hippocampal SC-CA1 synapses

We begin this section by summarizing the results from the limited *in vitro* hippocampal slice experiments on the importance of the time window of the activation of the D1/D5 receptors relative to the high-frequency stimulation (HFS) protocol used to induce the long-term potentiation (LTP) in the Schaffer collateral-CA1 pyramidal (SC-CA1) synapses. In a classical experiment [2], Huang and Kandel showed that the 15 minutes administration of D1/D5 agonist SKF 38393 212 minutes before the HFS protocol enhanced the LTP of the synapse immediately after the HFS protocol but decayed to approximately the fEPSP level induced by only HFS after 2 hours. In the same experiment, the authors found no noticeable changes in the LTP of a SC-CA1 synapse when they administered SKF 38393 50 minutes after the HFS protocol. In [34], Navakkode et al. investigated the HFS-mediated LTP modulation in SC-CA1 synapses by dopamine. In *in vitro* slice experiments, the authors showed that the application of 50*μM* of dopamine in three five minute pulses spaced ten minutes apart three hours prior to an HFS protocol of three trains of 100 pulses at 100 *Hz* induced a transient enhancement of HFS induced LTP that decayed to the level LTP induced by only HFS in 2 hours. These results highlight the activation of the D1/D5 receptors a long time before HFS significantly enhances the HFS-induced LTP of the SC-CA1 synapse. By contrast, the activation of D1/D5 receptors more than an hour after HFS does not significantly modulate the HFS-induced LTP. However, it is not clear from these results whether the activation of D1/D5 receptors can still modulate the HFS-induced LTP of a SC-CA1 synapse if one administers SKF 38393 sufficiently close to the timing of the HFS protocol (a few minutes before or after HFS). A few experiments [35, 36] have investigated this question in hippocampal slices under the bath application of SKF 38393. The results from these experiments suggest that the activation of D1/D5 receptors at an earlier time than 200 minutes can also modulate the HFS-induced LTP of a SC-CA1 synapse. In summary, the available experimental results suggest that the relative timing between the HFS protocol and the activation of the D1/D5 receptors plays an essential role in the dopaminergic modulation of HFS-induced LTP of a SC-CA1 synapse. However, it is still not clear from these experiments how various concentrations and relative timings of D1/D5 agonists modulate the temporal dynamics of the HFS-induced LTP of a SC-CA1 synapse.

We developed, for the first time, a hybrid model capable of predicting the temporal dynamics of the modulation of HFS-induced LTP of the SC-CA1 synapse by various concentrations and relative timings of D1/D5 agonists, SKF 38393, dopamine (DA), and 6-bromo-APB, that integrate the present experimental findings under diverse stimulation protocols. We describe our complete model of the HFS-induced LTP modulation by D1/D5 agonists in the Materials and methods section. Briefly, we have used a published, experimentally validated, single compartment biophysiological model of CA1 pyramidal neuron in the form of Hodgkin-Huxley formalism [29] and a phenomenological model of SC-CA1 synapse [30] to model the electrophysiological activity of CA1 pyramidal neuron in response to the electrical and pharmacological stimulations of the SC-CA1 synapse (see Eqs (1), (2a)–(2h), (3a)–(3r), and (4a)–(4h) in the Materials and methods section). To model the dynamics of HFS-induced LTP and LFS-induced LTD, we modified a published phenomenological model of LTP/LTD [31, 32] and inferred the model parameters using an approximate Bayesian inference approach from the available hippocampal slice experimental data [35, 37, 38] on the HFS/LFS induced LTP/LTD (see Eqs (6a)–(8) and Table 4 in the Materials and methods section). Finally, we developed a phenomenological model to integrate the dose-dependent temporal effects of D1/D5 agonists (SKF 38393, DA and 6-APB) relative to the HFS/LFS on the SC-CA1 synaptic LTP/LTD induced by HFS/LFS (see Eqs (14a)–(14d) and Eqs (15a)–(15r) in the Materials and methods section). We inferred our model parameters in an approximate Bayesian inference paradigm from the available rat hippocampal slice experimental data [2, 4, 8, 27, 34, 39] in the form of % fEPSP slope from control in response to D1/D5 agonists and HFS/LFS simultaneously (see Tables 5 and 6 in the Materials and methods section).

We used our developed model to quantify how different relative timings of DA and the D1/D5 agonist SKF 38393 modulate the HFS-induced LTP of a SC-CA1 synapse. We first used our model under a similar experimental protocol condition as described in [2] to demonstrate the capability of our model in predicting the experimental results shown in [2]. To do so, we delivered 50 *μM* SKF 38393 for 15 minutes to our SC-CA1 model more than 200 minutes before or 50 minutes after the HFS protocol. For the HFS protocol, we used 3 trains of 100 *Hz* pulses for 1 second with a 10 minute inter-train interval and stimulated our SC-CA1 synapse model (see Eqs (4a)–(4h) and Eqs (6a)–(8) in the Materials and methods section) to induce LTP. We measured the change in the SC-CA1 synaptic strength through the percentage change in the slope of evoked fEPSPs normalized to the slope of evoked fEPSPs prior to any induction of potentiation. Figures 1A and 1B show the comparison between the experimental [2] and model predicted D1/D5 enhancement in the HFS-induced LTP by SKF 38393. As shown in Figure 1A, our model makes a quantitative prediction of the time-dependent enhancement in the late LTP reported by Huang and Kandel [2] when SKF 38393 is delivered 200 minutes before the HFS administration. Moreover, our model predicts the experimental observation from Huang and Kandel [2] that 50 *μM* administration of SKF 38393 leads to no significant changes in the HFS-induced LTP when it is delivered 50 minutes after the HFS protocol (see Figure 1B).

**Fig 1.**
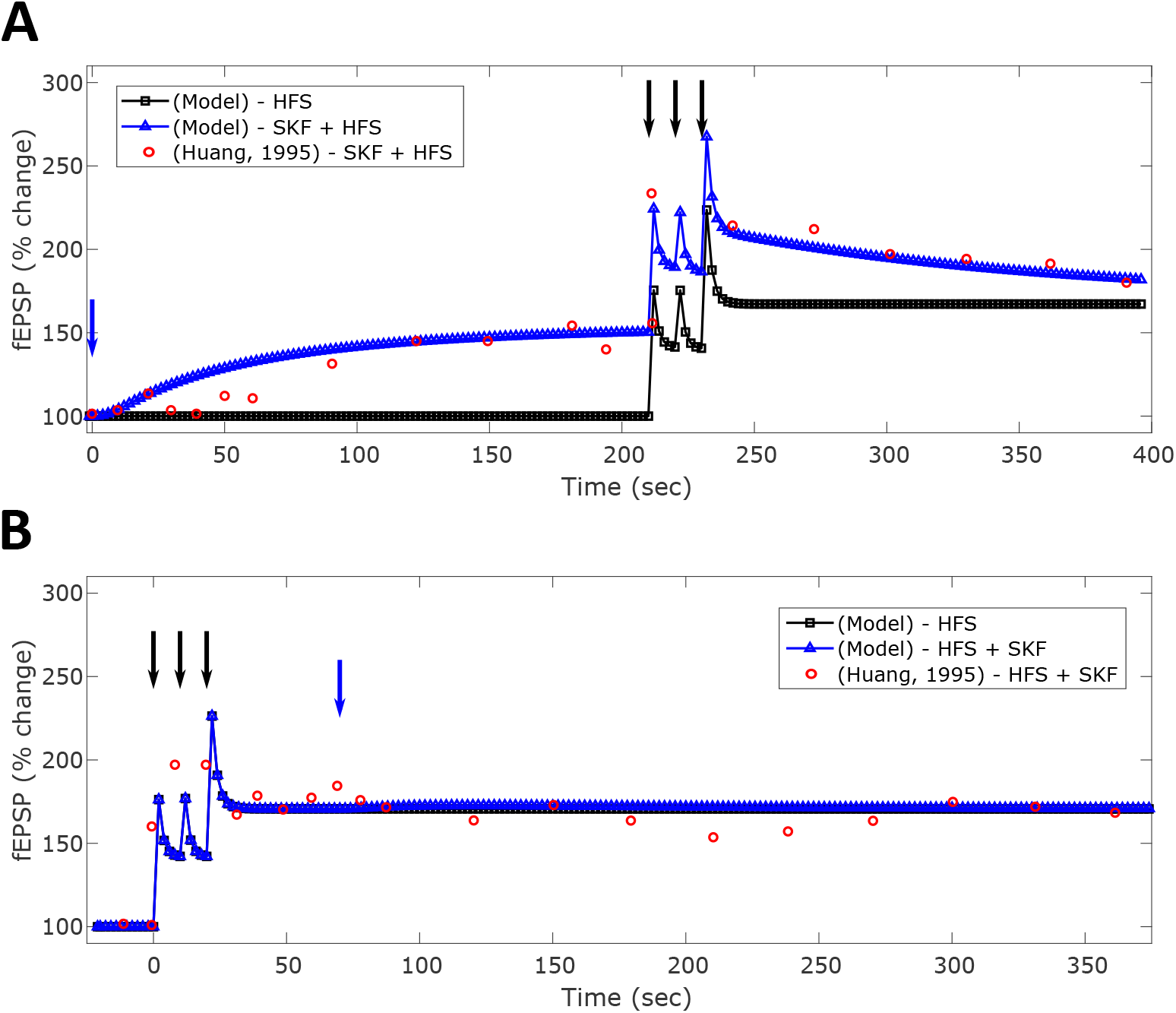
Quantitative comparison between the model predicted and experimentally observed [2] modulation of HFS-induced LTP in hippocampal SC-CA1 synapse by D1/D5 agonist SKF 38393. The induced LTP of the SC-CA1 synapse is measured in terms of the percentage (%) change in evoked fEPSP slope from the control. The black-squares represents the application of the HFS protocol of three trains of pulse at 100 Hz for 1 second with a 10 minute intertrain interval, while the blue-triangles represent the same HFS protocol in combination with 50 *μM* SKF 38393 for 15 minutes applied at various time distances from the HFS protocol (Δ*t* = *t*_*SKF*_ − *t*_*HFS*_). **(A)** shows the HFS-induced LTP without (black-squares) and with (blue-triangles) 50 *μM* SKF 38939 delivered 212 minutes before the HFS protocol. The experimentally reported SKF 38393 enhancement of LTP [2] is shown as the red-circles (Δ*t* = 212 min). **(B)** shows the SKF 38393 enhancement of the HFS-induced LTP when 15 minutes of 50 *μM* SKF 38393 is delivered 50 minutes after the HFS administration (Δ*t* = 50 min).

Next, we validated our model’s capability in predicting the experimental results from [34] on the transient enhancement in the late LTP by 50 *μM* application of dopamine (DA). Since we fitted the dopaminergic portion of our model to specific D1/D5 agonists (see Eqs. (15a)–(15h) in the Materials and methods section), we used the DA specific parameters specified in Table 5 to model the spatiotemporal effect of DA on the HFS-induced late LTP of the SC-CA1 synapse. By following the exact stimulation protocol used in the experiment [34], we injected 50 *μM* of DA in three pulses 165 minutes before the HFS protocol consisting of three trains of 100 pulses at 100 *Hz* in our model and measured the change in the fEPSP slope. As shown in Figure 2, our model is able to quantitatively predict an enhancement of approximately 27% in the HFS-induced LTP by DA, measured 60 minutes after the HFS protocol, as reported in the experiment.

**Fig 2.**
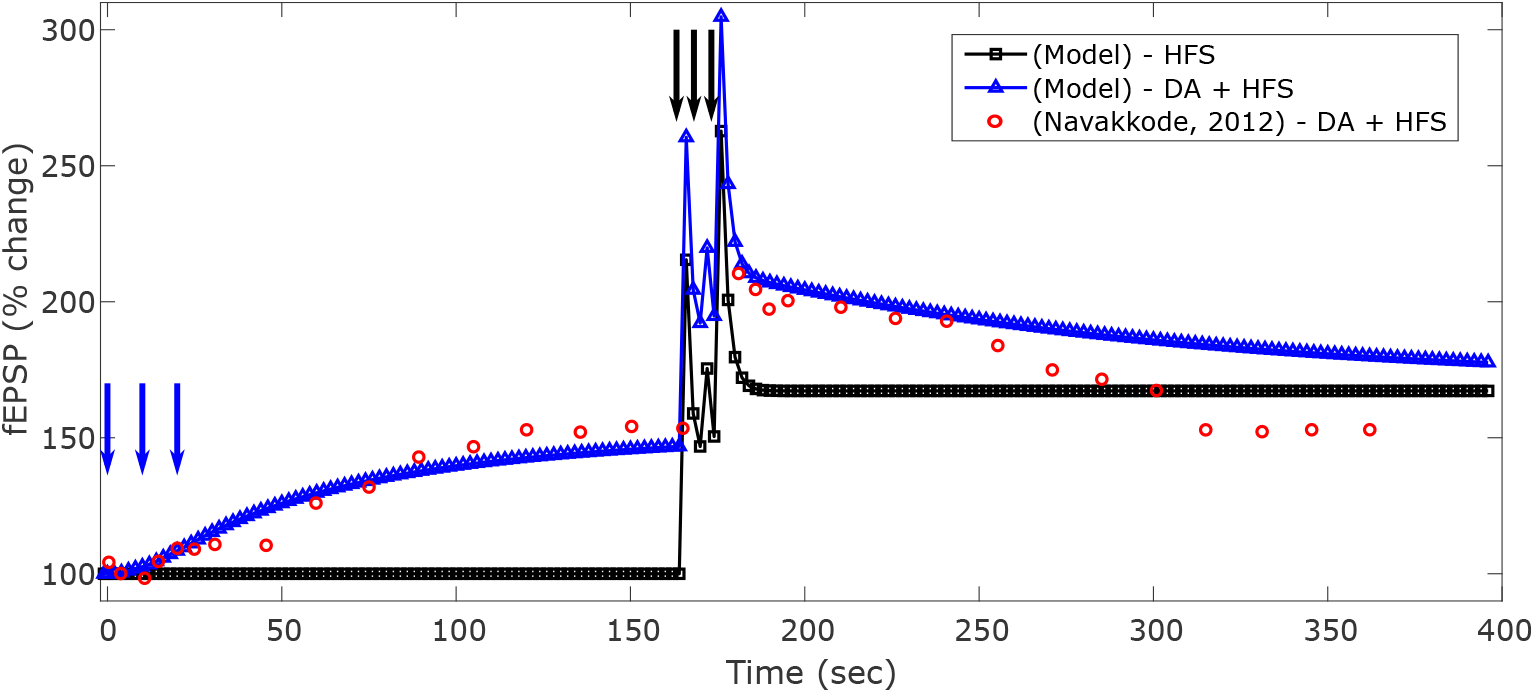
Quantitative comparison between the model predicted and experimentally observed [34] modulation of HFS-induced LTP in hippocampal SC-CA1 synapse by dopamine (DA). The induced LTP of the SC-CA1 synapse is measured in terms of the percentage (%) change in evoked fEPSP slope from the control. DA is applied 165 minutes before the HFS stimulation protocol of three trains of 100 pulses at 100 *Hz* with a 5-minute inter-train interval. The HFS induced LTP when (blue-triangles) 50 *μM* DA is delivered before the HFS protocol is compared to the HFS only induced LTP (black-squares). The experimentally reported dopaminergic enhancement of LTP [34] is shown as the red-circles (Δ*t* = −165 min).

Finally, we used our model to make specific predictions by conducting two simulation experiments where we administered 50 *μM* SKF 38393 30 minutes before the HFS protocol (see Figure 3A) and 10 minutes after the HFS protocol (see Figure 3B). Based on the experimental results on the two extreme cases (i.e., SKF 38393 application 200 minutes before or 50 minutes after the HFS protocol) and the slow-onset potentiation of the SC-CA1 synapses by SKF 38393, one would expect a reduction in the SKF 38393 mediated LTP enhancement as the time difference between the SKF 38393 administration and the HFS protocol decreases. As shown in Figure 3A, the injection of 50 *μM* SKF 38393 30 minutes before HFS in our model led to approximately 20% enhancement in the HFS-induced late LTP 60 minutes after HFS compared to 30% enhancement when SKF 38393 was delivered 200 minutes before the HFS protocol. Additionally, the injection of SKF 38393 10 minutes after the end of the HFS protocol resulted in no significant enhancement (approximately 4% enhancement after 60 minutes of the HFS protocol) in HFS-induced late LTP (see Figure 3B) due to the occlusion by HFS.

**Fig 3.**
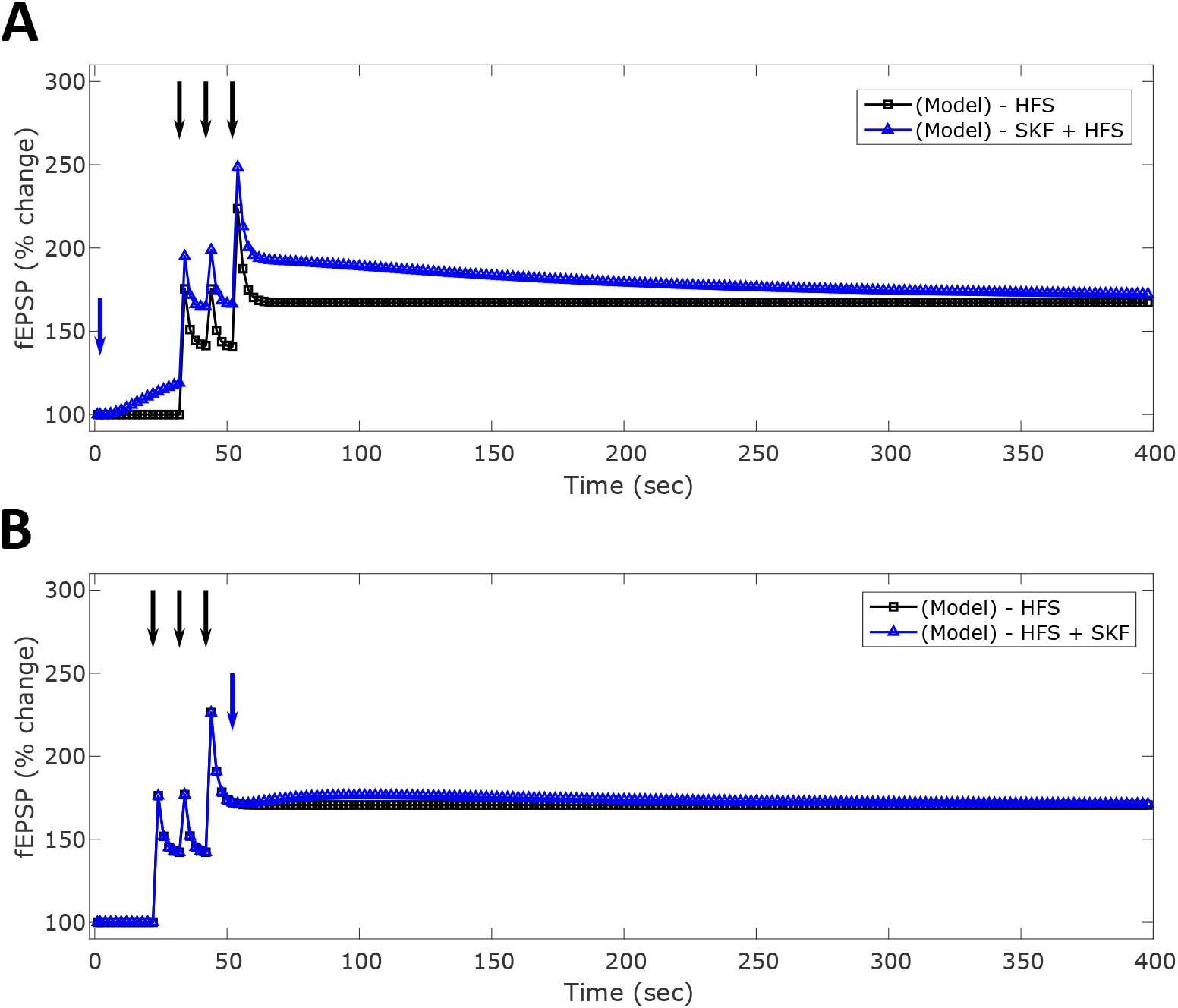
Predictions from the model on the modulation of HFS-induced LTP in the hippocampal SC-CA1 synapse by D1/D5 agonist SKF 38393 when the D1/D5 agonist SKF 38393 is delivered closer in time relative to the applied HFS protocol. In these simulation results, SKF 38393 is applied 30 minutes before **(A)** and 10 minutes after **(B)** the HFS protocol of 3 trains of 100 pulses at 100 Hz with a 10 minute inter-train intervals. The induced LTP of the SC-CA1 synapse is measured in terms of the percentage (%) change in evoked fEPSP slope from the control. **(A)** shows approximately 20% enhancement immediately after the HFS protocol in the HFS-induced LTP by SKF 38393 when delivered 30 minutes before the HFS protocol (blue-triangles) of (Δ*t* = 30 min). The LTP induced by only HFS is shown as the black-squares. **(B)** shows a small (negligible) enhancement in the HFS-induced LTP when SKF 38393 is delivered 10 minutes (blue-triangles) after the HFS protocol (Δ*t* = 10 min).

In sum, our model shows reasonable quantitative predictions of existing experimental results on the HFS-induced late LTP modulation of SC-CA1 synapses by SKF 38393 and DA. It generates new predictions based on the relative timing between HFS and SKF 38393 administrations by capturing the dynamical mechanism of the HFS-induced late LTP modulation by SKF 38393 and DA. Moreover, our simulation results indicate that the enhancement in the HFS-induced LTP of the hippocampal SC-CA1 synapses by D1/D5 agonists strongly depends on the timing and order of the applied agonists relative to the HFS protocol in a nonlinear fashion.

### 6-bromo-APB enhances weak HFS-induced early LTP in hippocampal SC-CA1 synapses

In hippocampal slices from rats [40], Otmakhova and Lisman showed that the 5 minutes application of a D1/D5 receptor agonist 6-bromo-APB 5 minutes before a weak HFS protocol of 10 bursts of 4 pulses at 100 Hz with a 30 ms interval enhanced the HFS-induced LTP in the hippocampal SC-CA1 synapses by approximately 11% immediately after the weak HFS protocol and approximately 8% after 40 minutes. Furthermore, the application of 6-bromo-APB 35 minutes after the weak HFS protocol produced no significant changes in the weak HFS-induced LTP. These results highlight the time dependent modulation of a weak HFS-induced LTP in SC-CA1 synapses by the D1/D5 agonist 6-bromo-APB. Moreover, 6-bromo-APB interacts with HFS in a nonlinear fashion to modulate the HFS-induced LTP.

Before we used our model to predict the above experimental results, we first validated our model’s capability in predicting slow-onset potentiation induced by 6-bromo-APB in the absence of a HFS protocol for inducing LTP in the SC-CA1 synapse. *In vitro* experimental data from [4] shows that the application of 5*μM* 6-bromo-APB for 5 minutes three times (5 minutes inter-spacing) leads to a slow-onset potentiation of the SC-CA1 synapse. Under the same protocol as described in [4], we used our model (see Eqs. (15a)–(15h) in the Materials and methods section) to predict this slow-onset potentiation by 6-bromo-APB. Figure 4A compares the prediction from our model with the experimental data in [4]. As shown in this figure, our model captures the essential dynamics to predict the slow-onset potentiation induced by 6-bromo-APB.

**Fig 4.**
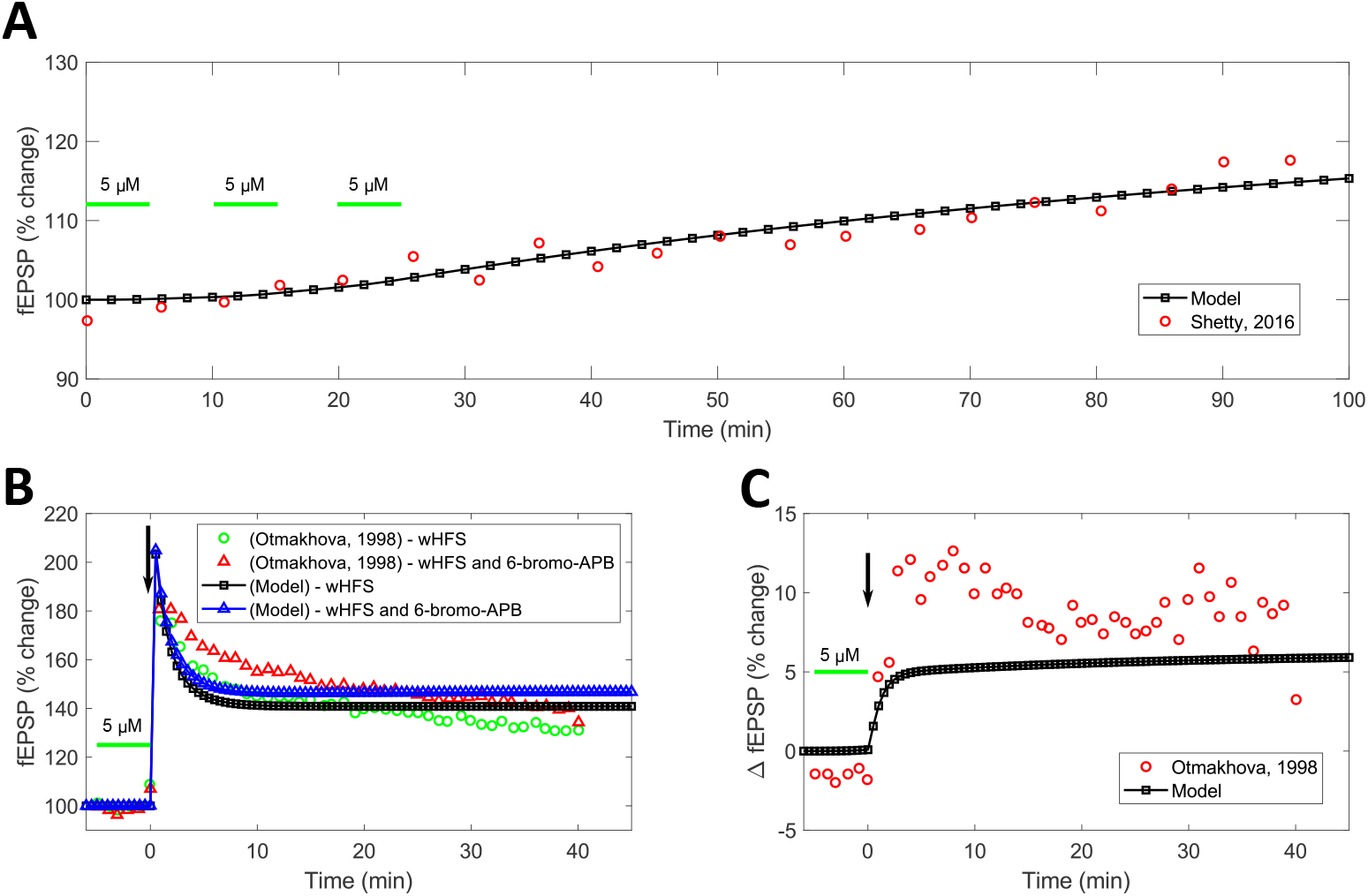
Comparison between the model predicted and experimentally observed modulation of weak HFS-induced LTP in hippocampal SC-CA1 synapse by D1/D5 agonist 6-bromo APB. The induced LTP of the SC-CA1 synapse is measured in terms of the percentage (%) change in evoked fEPSP slope from the control. **(A)** shows the slow-onset potentiation due to the application of 5 *μM* of 6-bromo-APB for 5 minutes with 5 minute intervals (green-bars) observed in the experiment [4] (red-circles) and predicted by our model (black-squares). **(B)** shows the dopaminergic enhancement by 5 *μM* of 6-bromo-APB for 5 minutes (green-bar) of LTP induced by a weak HFS protocol of 10 bursts of 4 pulses at 100 Hz with a 30 *ms* interval (black-arrow). The green-circles show the LTP induced by the weak HFS protocol alone and the red-triangles show the LTP induced by the weak HFS with 6-bromo-APB in the experiment [40]. In the result predicted by our model, the induced LTP from a weak HFS protocol is shown as black-squares and the LTP induced by a weak HFS protocol with the 5 *μM* of 6-bromo-APB is shown as the blue-triangles. **(C)** shows the absolute dopaminergic enhancement of LTP (Δ fEPSP) by 6-bromo-APB in the experiment [40] (black-squares) and our model (red-squares) computed by subtracting the measured potentiation of the weak HFS plus 6-bromo-APB from potentiation by weak HFS alone.

We then used our model to predict the experimental results from [40] on the modulation of a weak HFS-induced LTP by 6-bromo-APB. Figure 4B compares the prediction from our model with the experimental data from [40] when 6-bromo-APB was applied 5 minutes before the HFS protocol for 5 minutes. As shown in this figure, our model is able to capture the key dynamical features presented in the data qualitatively, such as a sharp enhancement in the HFS-induced LTP just after the HFS protocol (see Figure 4C) and the temporal changes in the LTP over time. Figure 4C highlights the faster dynamics of dopaminergic potentiation when the dopaminergic agonist 6-bromo-APB is applied close to a weak HFS protocol compared to the much slower potentiation dynamics observed by the application of only 6-bromo-APB in Figure 4A.

While our model accurately captures the dopaminergic potentiation dynamics, the model predictions of the overall change in SC-CA1 potentiation differs significantly from the experimental data quantitatively. We wondered whether this is because of the differences in the HFS-induced LTP predicted from our model and the experimental data. Would our model predict the data quantitatively better if we had the exact HFS-induced LTP changes predicted by our model as in the experimental data? To investigate this question, we tuned the HFS model parameters in Eq. (6c) (*p*_*p*_ and *M*_*p*_) through hand-fitting to match the HFS-induced LTP data from the experiment in the absence of 6-bromo-APB. We then again used our model to predict the result on the modulation of the weak HFS-induced LTP by 6-bromo-APB. Figures 5A and 5B compare the prediction from our model and the experimental data. As shown in Figure 5A, our model’s performance improved significantly in predicting the experimental data quantitatively. This illustrates that our combined model (HFS+6-bromo-APB) captures the essential biophysiological mechanisms underlying the modulation of HFS-induced LTP by 6-bromo-APB in the hippocampal SC-CA1 synapses.

**Fig 5.**
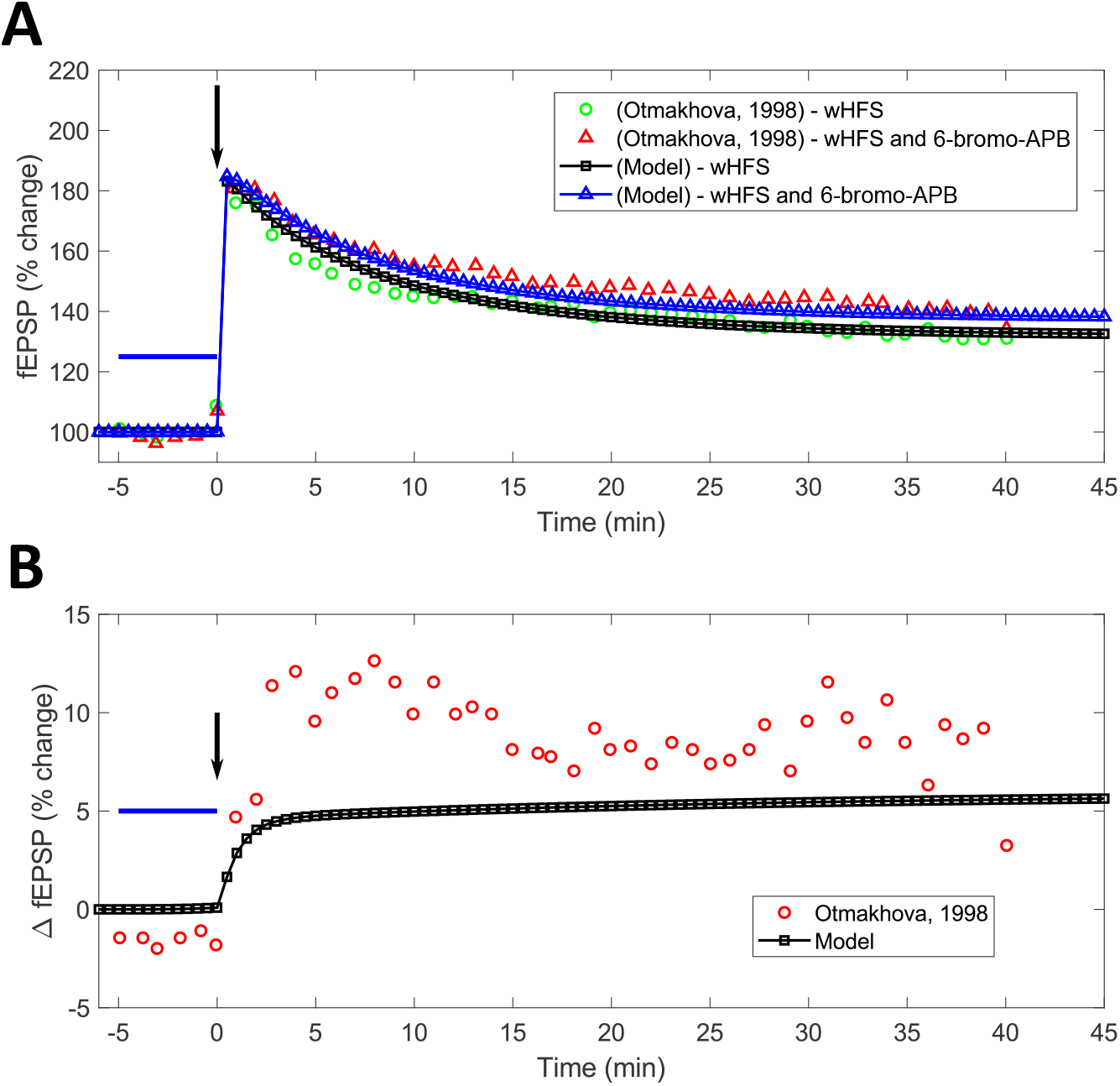
Enhancement in the model predictions shown in Figure 4 with improved HFS-induced LTP prediction. We tuned the model parameters of the HFS model to match the experimental data on weak HFS-induced LTP from [40]. Then we used our model to again predict the changes in the weak HFS-induced LTP after the application 5 *μM* of 6-bromo-APB for 5 minutes (green-bar). **(A)** shows the enhancement in the weak HFS-induced LTP by 6-bromo-APB with the new parameters. The experimental and model predicted data on the simultaneous application of weak HFS protocol and 6-bromo-APB are shown by the red-triangles and blue-triangles, respectively. The experimental and model predicted data on the weak HFS application alone are shown by the green-circles and black-squares, respectively. **(B)** shows the comparison between the absolute dopaminergic enhancement of LTP (Δ fEPSP) by 6-bromo-APB observed in the experiment [40] (black-squares) and predicted our model (red-squares) with the modified HFS model parameters. Δ fEPSP is computed by subtracting the measured potentiation of the weak HFS plus 6-bromo-APB from potentiation by the weak HFS alone. The modified HFS model parameters in Eq. (6c) are *p*_*p*_ = 1.5099 × 10^−6^ *ms*^−1^, *M*_*p*_ = 7.4938 × 10^−9^ *ms*^−1^, and *f* = 298.

Finally, we used our model to predict the experimental result on the modulation of the weak HFS-induced LTP by 6-bromo-APB when 6-bromo-APB was applied 35 minutes after the HFS protocol. Figure 6 compares our model’s prediction with the hand-fit parameters to the experimental data [40]. As shown in Figure 6, our model accurately predicts the experimental data, i.e., no significant enhancement in the LTP by 6-bromo-APB. This highlights our model’s capability at capturing the time dependency of dopaminergic modulation of SC-CA1 synaptic plasticity.

**Fig 6.**
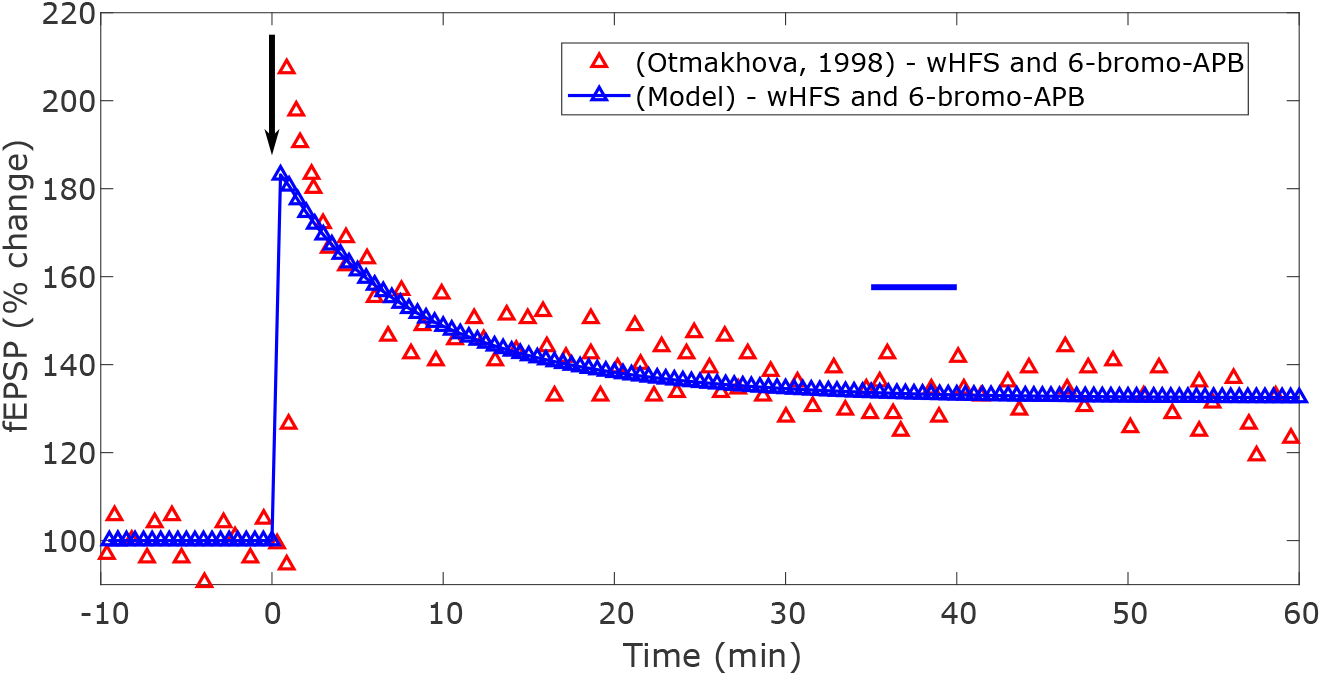
Comparison between the model predicted and the experimentally observed [40] modulation of a weak HFS-induced LTP when 6-bromo-APB was delivered after the weak HFS protocol. The dopaminergic agonist 6-bromo-APB is applied 35 minutes after a weak HFS protocol of 10 bursts of 4 pulses at 100 Hz with a 30 *ms* interval (black-arrow). The model parameters of the HFS model is the same as the one used in Figure 5. The induced LTP of the SC-CA1 synapse is measured in terms of the percentage (%) change in evoked fEPSP slope from the control. The 6-bromo-APB enhanced weak HFS-induced LTP predicted by the model is shown as the blue-triangles and observed in the experiment as the red-triangles [40].

### Antagonizing D1/D5 receptors blocks HFS induced late-LTP in hippocampal SC-CA1 synapses

One of the critical questions in understanding dopamine’s role in the hippocampal SC-CA1 long-term synaptic plasticity is whether the basal level of dopamine is essential for the induction of LTP, or it only plays a role in modulating the late LTP. Several research groups have examined this question pharmacologically by blocking the D1/D5 receptors using a D1/D5 selective antagonist SCH 23390 in hippocampal slices from rats [2, 8]. Huang and Kandel [2] showed that pharmacologically blocking the D1/D5 receptors using SCH 23390 blocks the HFS-induced late LTP in hippocampal SC-CA1 synapses with a minimal effect on early LTP. Similarly, Sajikumar et al. [41] showed that the application of SCH 23390 blocked the late-LTP induced by a weak HFS protocol of 100 pulses at 100 *Hz* in the SC-CA1 synapses.

We again used our model described in the previous sections (see Eqs. (1),(6a) -(6d), (6f) - (6i), (15a) -(15r), and (18) in the Materials and methods section) to predict these experimental findings quantitatively. We incorporated the effect of SCH 233390 on the late LTP by introducing a model parameter *k*_*basal*_, which takes a nonzero value whenever SCH 23390 is applied to block the D1/D5 receptors (see Eq. (15o) in the Materials and methods section). The parameter *k*_*basal*_ decreases the HFS-induced LTP predicted by our model to the baseline in approximately 400 minutes when D1/D5 receptors are blocked with the simultaneous application of SCH 23390 and HFS, as observed in the experimental studies.

Figure 7A compares our model predictions with the experimental results from Huang and Kandel [2] and Sajikumar et al. [41] on the blockade of weak HFS (100 pulses at 100 *Hz*) induced late LTP by SCH 23390. As shown in this figure, our model reasonably predicts the decay of the weak HFS-induced LTP to the baseline (i.e., 100%) in approximately 400 minutes, as observed in the experimental data. Figure 7B compares our model predictions with the experimental results from Huang and Kandal [2] on the blockade of a strong HFS protocol (3 trains of 100 pulses at 100 *Hz*) induced LTP in the hippocampal SC-CA1 synapses by SCH 23390. As shown in this figure, our model struggles to make quantitative prediction of the changes in the early LTP but provides a reasonable prediction of changes in the late LTP.

**Fig 7.**
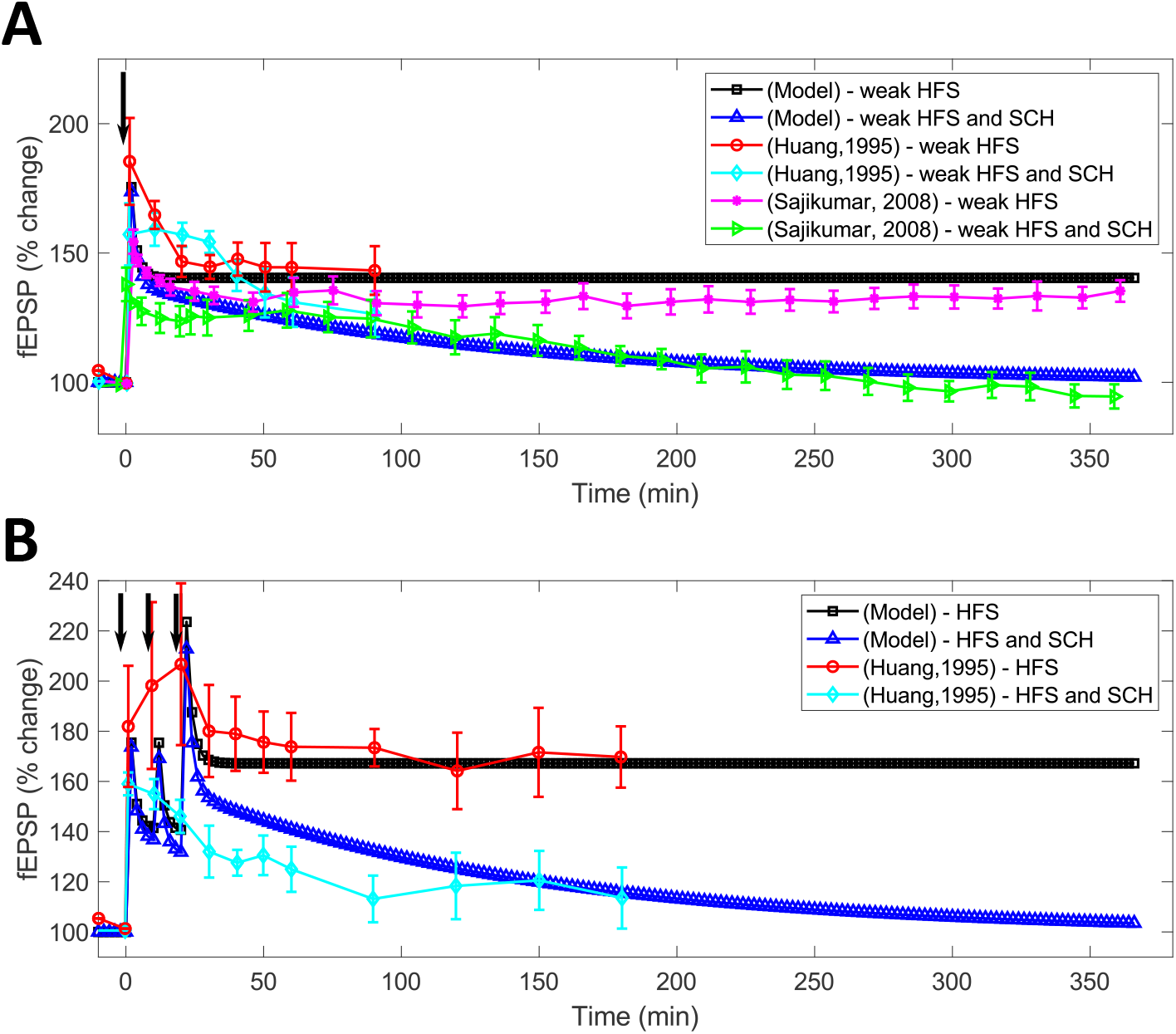
The effect of basal dopamine level on HFS-induced LTP. **(A)** compares the LTP induced by a weak HFS protocol of 100 *Hz* stimulation in the presence of a D1/D5 antagonist SCH 23390 from experiments (shown in cyan-diamonds and green-triangles) [2, 41] with the predictions from our model (blue-triangles). The only HFS-induced LTP from the experiments are shown in red-circles and magenta-stars and from the model is shown in black-squares. **(B)** compares the LTP induced by a strong HFS protocol (3 trains of 100 *Hz*) in the presence of a D1/D5 antagonist SCH 23390 from the experiment (cyan-diamond) [2] with the prediction from our model (blue-triangles). The only HFS-induced LTP from the experiment is shown in red-circles and from the model is shown in black-squares.

### Concentration dependent effect of SKF 38393 on the modulation of HFS-induced LTP

It has been shown in hippocampal slice experiments that SKF 38393 induces a slow-onset potentiation in SC-CA1 synapses in the absence of a LTP induction protocol [2, 4] and this potentiation strongly depends on the SKF 38393 concentration in a nonlinear fashion. However, no experimental results exist on how various concentrations of SKF 38393 impact the modulation of HFS-induced late LTP. After validating our model with the available experimental data on the modulation of HFS-induced late LTP by D1/D5 agonists and antagonists in the previous sections, we used our model (see Eqs. (1),(6a) -(6d), (6f) - (6i), (15a) -(15r), and (18)) with SKF 38393 parameters given in Table 5 in the Materials and methods section to investigate this question.

We systematically investigated the effect of seven different concentrations of SKF 38393, ranging between 1 − 50 *μ*M, on the modulation of strong HFS-induced LTP in the hippocampal SC-CA1 synapses while varying the relative time difference between the SKF 38393 injection and the applied HFS protocol. Figures 8A, 8B and 8C show the predictions of our model when seven different concentrations of SKF 38393 (i.e., 1 *μ*M, 2 *μ*M, 5 *μ*M, 10 *μ*M 15 *μ*M, 25 *μ*M, and 50 *μ*M) were administered 212 minutes, 30 minutes, and 15 minutes, respectively, before a strong HFS protocol (3 trains of 100 pulses at 100 Hz). We applied each concentration of SKF 38393 for 15 minutes, which induced the slow-onset potentiation in the SC-CA1 synapse before the HFS protocol. As shown in these figures, our model predicted a slow decay of the SKF 38393 induced slow-onset potentiation in the hippocampal SC-CA1 synapse after the application of a strong HFS protocol. Particularly, our model predicted a bifurcation regime where the higher concentration of SKF 38393 (5 *μ*M or higher) led to significant modulation of HFS-induced LTP while lower concentration of SKF 38393 (below 5 *μ*M) led to insignificant changes in the HFS-induced LTP. Moreover, the observed concentration dependent enhancement of HFS-induced LTP decreased as the time difference between the application the HFS protocol and the injection of SKF 38393 increased. This highlights not only the importance of timing but also the dopamine agonist concentration on the dopaminergic modulation of a strong HFS-induced potentiation of the SC-CA1 synapse.

**Fig 8.**
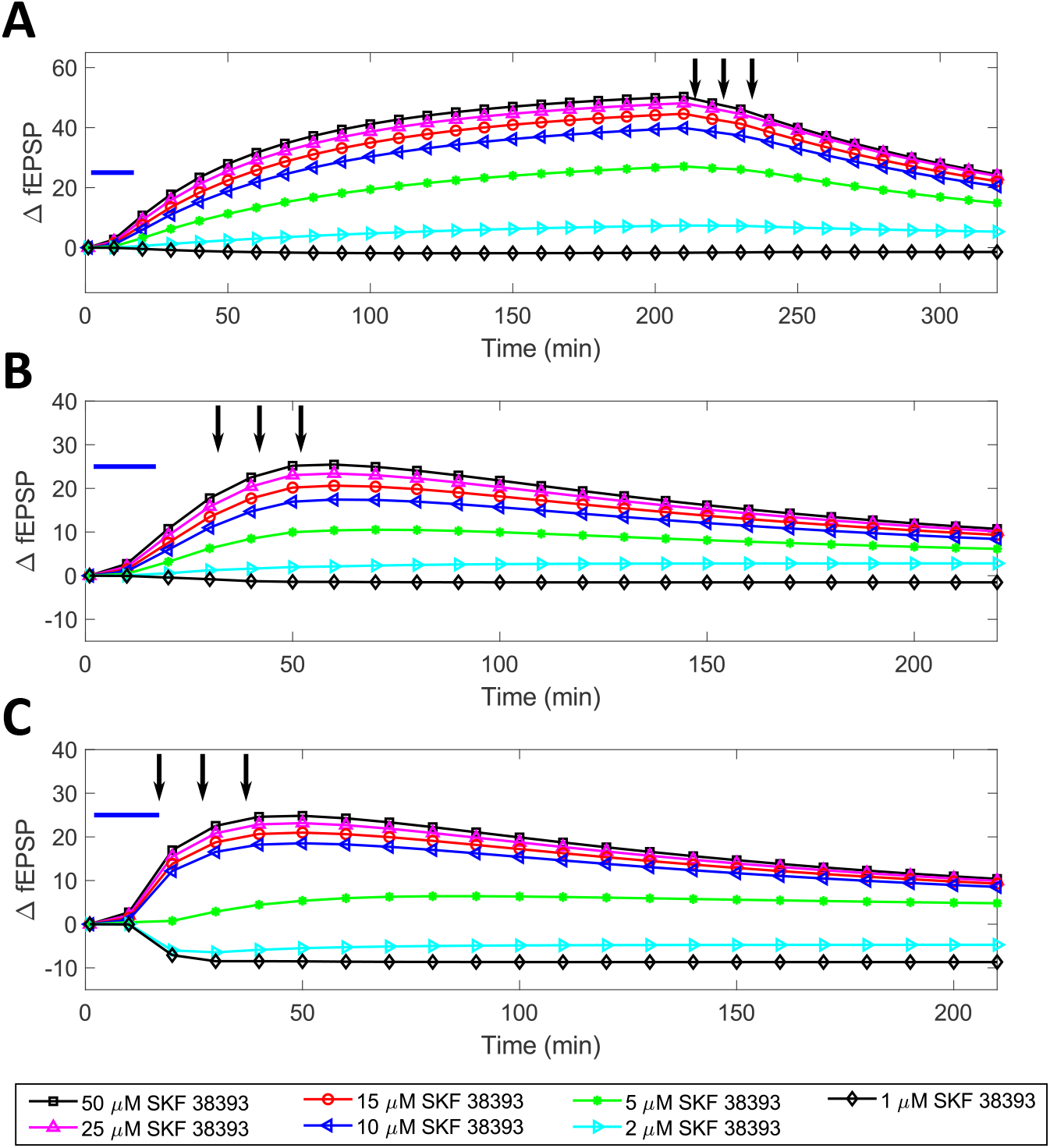
Model predictions on the effect of the concentration and timing of SKF 38393 relative to a strong HFS protocol (3 trains of 100 pulses at 100 Hz) in modulating HFS-induced LTP when SKF 38393 was delivered before the HFS protocol. SKF 38393 (blue-bar) was delivered **(A)** 212 minutes (Δ*t* = 212 min), **(B)** 30 minutes (Δ*t* = 30 min), and **(C)** 15 minutes (Δ*t* = 15 min) before the strong HFS protocol (black-arrow).

We noted in Figure 8C that when we applied SKF 38393 immediately before the HFS protocol, our model predicted a bifurcation regime in the modulation of the HFS-induced LTP depending on the concentration of SKF 38393. At a high concentration (5 – 50 *μM*), SKF 38393 further potentiated the HFS-induced LTP whereas at a low concertation (1 – 2 *μM*), SKF 38393 depressed the HFS-induced LTP. A potential mechanism underlying this observation may be the domination of the PLC pathway at a low concentration of SKF 38393, which has been shown to be critical for the induction of LTD [42, 43]. It has been hypothesized that there exists a D1-like dopamine receptor that is coupled to a Gq-protein and selectively activates the phospholipase C (PLC) pathway [7, 44]. This receptor may explain the observation that low concentrations of SKF 38393 induce slight depotentiation of the SC-CA1 synapse, as also observed in [4]. Furthermore, HFS primarily activates the CAMKII/PKA pathway [45] but also slightly activates the PLC pathway [46]. Therefore, the CAMKII/PKA pathway dominates to produce LTP. If a low concentration of SKF 38393 arrives immediately before the HFS protocol, the PLC pathway may still be active and could be boosted by the HFS protocol. The higher activity of the PLC pathway may slightly counteract the CAMKII/PKA pathway and reduce the level of LTP induced by the HFS protocol.

Additionally, our model also correctly predicted the dynamical features of the nonlinear interaction between SKF 38393 and HFS when we applied SKF 38393 just before the HFS protocol. As shown in Figure 5B, the application of the weak HFS protocol immediately after the administration of a dopamine agonist 6-bromo-APB quickened the dynamics and amplified the effect of the agonist on the SC-CA1 synapse. Therefore, high concentrations of SKF 38393, which induced slow-onset potentiation [4], quickly enhanced LTP induced potentiation, while low concentrations of SKF 38393, which have been shown in experiment to induce a slight depotentiation [4], depressed LTP induced potentiation.

We then used our model to investigate the concentration-dependent modulation of HFS-induced LTP by SKF 38393 delivered after the HFS protocol (3 trains of 100 pulses at 100 Hz). Figures 9A, 9B, and 9C show the model predicted enhancement in the HFS-induced LTP by SKF 38393 at seven different concentrations (1 *μ*M, 2 *μ*M, 5 *μ*M, 10 *μ*M 15 *μ*M, 25 *μ*M, and 50 *μ*M) when SKF 38393 was delivered 10, 30, and 60 minutes, respectively, after the HFS protocol. As noted in these figures, the overall enhanced potentiation in the HFS-induced LTP by SKF 38393 decreased with the distance between the applied SKF 38393 and the HFS protocol. This supports our limited resources hypothesis, since more resource may be consumed by the HFS-induced LTP consolidation with the increased distance between the HFS and SKF 38393. Moreover, the potentiation by SKF 38393 decreased with the decrease in the SKF 38393 concentration.

**Fig 9.**
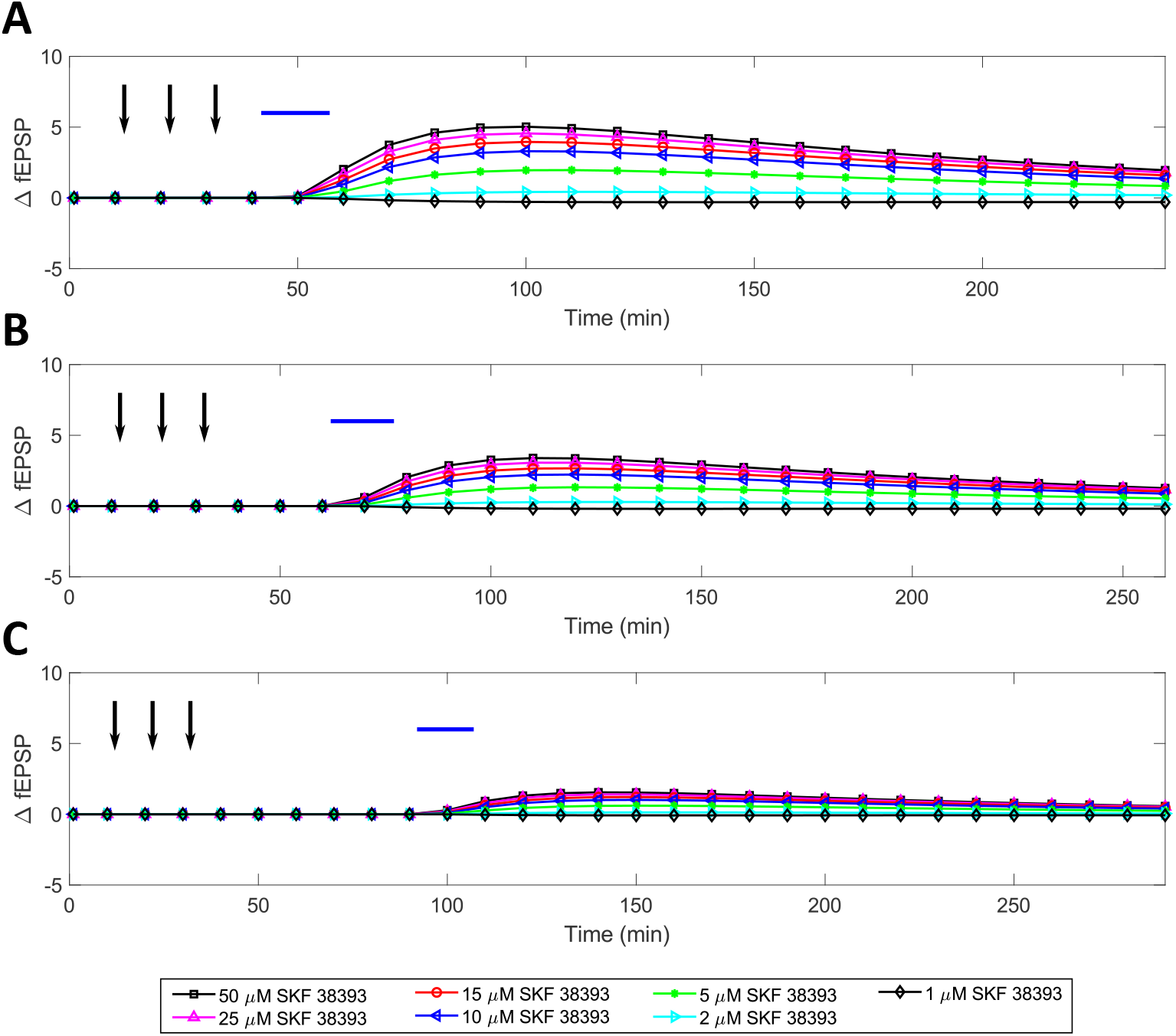
Model predictions on the effect of the concentration and timing of SKF 38393 relative to a strong HFS protocol (3 trains of 100 pulses at 100 Hz) in modulating HFS-induced LTP when SKF 38393 was delivered after the HFS protocol. SKF 38393 (blue-bar) was delivered **(A)** 10 minutes (Δ*t* = 10 min), **(B)** 30 minutes (Δ*t* = 30 min), and **(C)** 60 minutes (Δ*t* = 60 min) after the end of the strong HFS protocol (black-arrow).

Next, we investigated the effect of the same seven concentrations of SKF 38393 on the modulation of a weak HFS-induced LTP as the time difference was varied between the injection of SKF 38393 and the application of the weak HFS (100 pulses at 100 *Hz*). Based on our limited resource hypothesis, we expect that the weak HFS-induced LTP in SC-CA1 synapses will further be potentiated by the higher concentrations of SKF 38393. Particularly, SKF 38393 can convert the weaker levels of LTP to stronger levels of LTP in SC-CA1 synapse if a high concentration of SKF 38393 is administered significantly before the HFS protocol.

Figures 10A, 10B, 10C compare predictions from our model when SKF 38393 was applied 212, 30, and 15 minutes, respectively, before the weak HFS protocol. In each case, the injection of SKF 38393 alone induced the slow-onset potentiation of the SC-CA1 synapse that plateaued after the weak HFS protocol in all cases except when a concentration greater than 10 *μM* was applied 212 minutes before a weak HFS protocol. Since the weak HFS protocol alone doesn’t saturate the induced LTP, concentrations of SKF 38393 greater than 10 *μM* delivered 212 minutes before the weak HFS protocol enhanced the LTP to the saturated LTP level typically achieved by a strong HFS protocol. Thus, these high concentrations and timing of SKF 38393 injection slowly decayed back to the saturated LTP level. When we applied the weak HFS protocol immediately after the injection of SKF 38393 (see Figure 10C), the weak HFS protocol amplified the effect of the SKF 38393 modulation. Importantly, the high concentrations of SKF 38393 enhanced the weak HFS-induced LTP, while the low concentrations of SKF 38393 suppressed the weak HFS-induced potentiation.

**Fig 10.**
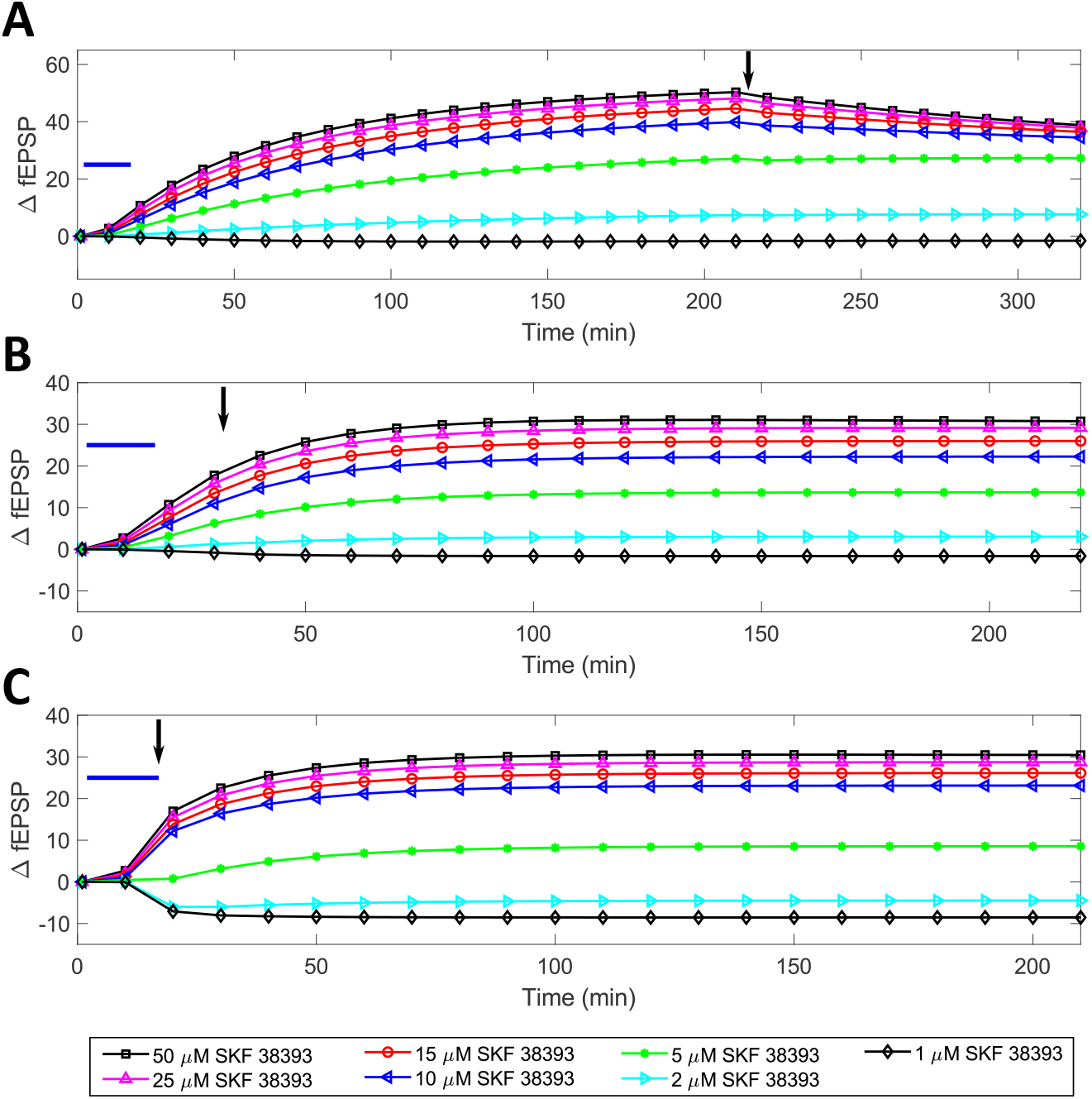
Model predictions on the effect of the concentration and timing of SKF 38393 relative to a weak HFS protocol (1 train of 100 pulses at 100 Hz) in modulating HFS-induced LTP when SKF 38393 was delivered before the HFS protocol. SKF 38393 (blue-bar) was delivered **(A)** 212 minutes (Δ*t* = 212 min), **(B)** 30 minutes (Δ*t* = 30 min), and **(C)** 15 minutes (Δ*t* = 15 min) before the weak HFS protocol (black-arrows).

Then, we applied the dopamine agonist SKF 38393 10, 30, and 60 minutes after the weak HFS protocol as shown in Figures 11A, 11B, and 11C, respectively. In contrast to the concentration dependent dopaminergic modulation of strong HFS induced LTP model predictions in Figures 8 and 9, the dopaminergic enhancement of the weak HFS-induced LTP did not decay for most of the concentrations and timings of SKF 38393. This supports our limited resource hypothesis. The weak HFS protocol consumed less resources and never saturated, which allowed the dopamine agonist to further enhance the weak HFS-induced LTP without decaying back to the pure HFS-induced LTP baseline. Additionally, the efficacy of the concentration dependent enhancement of the weak HFS-induced LTP decreased as the time interval between the administration of the weak HFS protocol and the injection of SKF 38393 increased.

**Fig 11.**
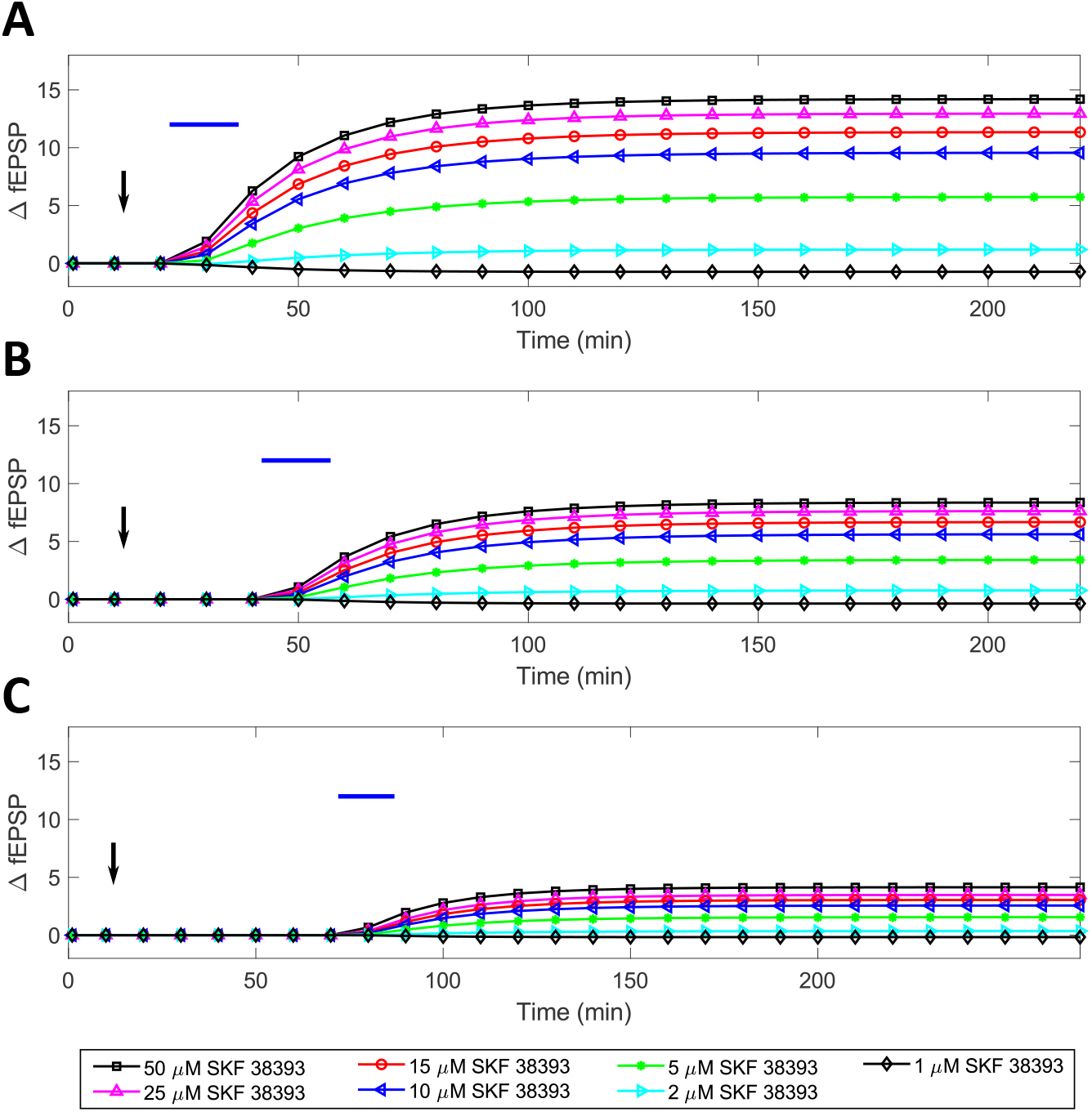
Model predictions on the effect of the concentration and timing of SKF 38393 relative to a weak HFS protocol (1 train of 100 pulses at 100 Hz) in modulating HFS-induced LTP when SKF 38393 was delivered after the HFS protocol. SKF 38393 (blue-bar) was delivered **(A)** 10 minutes (Δ*t* = 10 min), **(B)** 30 minutes (Δ*t* = 30 min), and **(C)** 60 minutes (Δ*t* = 60 min) after the weak HFS protocol (black-arrows).

### The relative time between D1/D5 receptors activation and LFS protocol significantly impacts the temporal modulation of LFS-induced LTD in hippocampal SC-CA1 synapses

We begin this section by summarizing the results from the limited *in vitro* hippocampal slice experiments on the importance of the time window of the activation of the dopamine D1/D5 receptors relative to a low-frequency stimulation (LFS) protocol used to induce the long-term depression (LTD) in the Schaffer collateral-CA1 pyramidal (SC-CA1) synapses. In *in vitro* hippocampal slice experiments [27], the authors applied 100 *μM* SKF 38393 for 20 minutes immediately after a LFS protocol of 1200 pulses at a frequency of 3 Hz. They observed a complete reversal of the LFS-induced LTD an hour after the SKF application. Next, the authors applied SKF 38393 an hour after the same LFS protocol in order to understand the temporal interaction of the two inputs. The application of 100 *μM* SKF 38393 one hour after the same LFS protocol produced no significant change in the SC-CA1 plasticity. In summary, the available experimental results suggest that the D1/D5 receptor mediated modulation of the LFS-induced LTD of the SC-CA1 synapse depends on the relative timing between the two inputs. It is not clear from these limited results whether the activation of D1/D5 receptors before an LFS protocol can reverse a LFS-induced LTD into LTP.

In order to investigate the temporal interaction of D1/D5 receptor activation by a dopamine agonist and a LFS protocol, we developed a SC-CA1 plasticity model able to predict the temporal dynamics of the dopaminergic modulation of LFS-induced LTD by various dopamine agonists (see Eqs. (1),(6a) -(6d), (6f) - (6i), (15a) -(15r), and (18) in the Materials and methods section). Our model for LFS-induced LTD modulation by D1/D5 agonists is similar to the HFS-induced LTP modulation model described in the previous sections except that we fitted the frequency-dependent plasticity parameters in Eqs. (6a) -(6d) and (6f) - (6i) using the LFS specific data. These parameters are described in Table 4.

We validated our SC-CA1 model with the experimental data from Mockett et. al. [27] where 100 *μM* SKF 38393 was delivered immediately and 60 minutes after a LFS protocol of 1200 pulses at 3 Hz. Figures 12A and 12B compare the synaptic plasticity change in the SC-CA1 synapse predicted by our model with the experimental data when SKF 38393 was applied immediately and 60 minutes, respectively, after the LFS protocol used in the experiment. As shown in Figure 12A, when we delivered SKF 38393 immediately after the LFS protocol in our model, the LFS-induced LTD reversed completely. This matched the reversal of LFS-induced LTD observed in experimental data [27]. Next, we applied SKF 38393 60 minutes after a LFS protocol of 1200 pulses at 3 Hz. The application of SKF 38393 60 minutes after the LFS protocol induced only a slight potentiation of approximately 4% (see Figure 12B). A similar inflection was also observed in the experimental data [27], although it was not large enough to be statistically significant.

**Fig 12.**
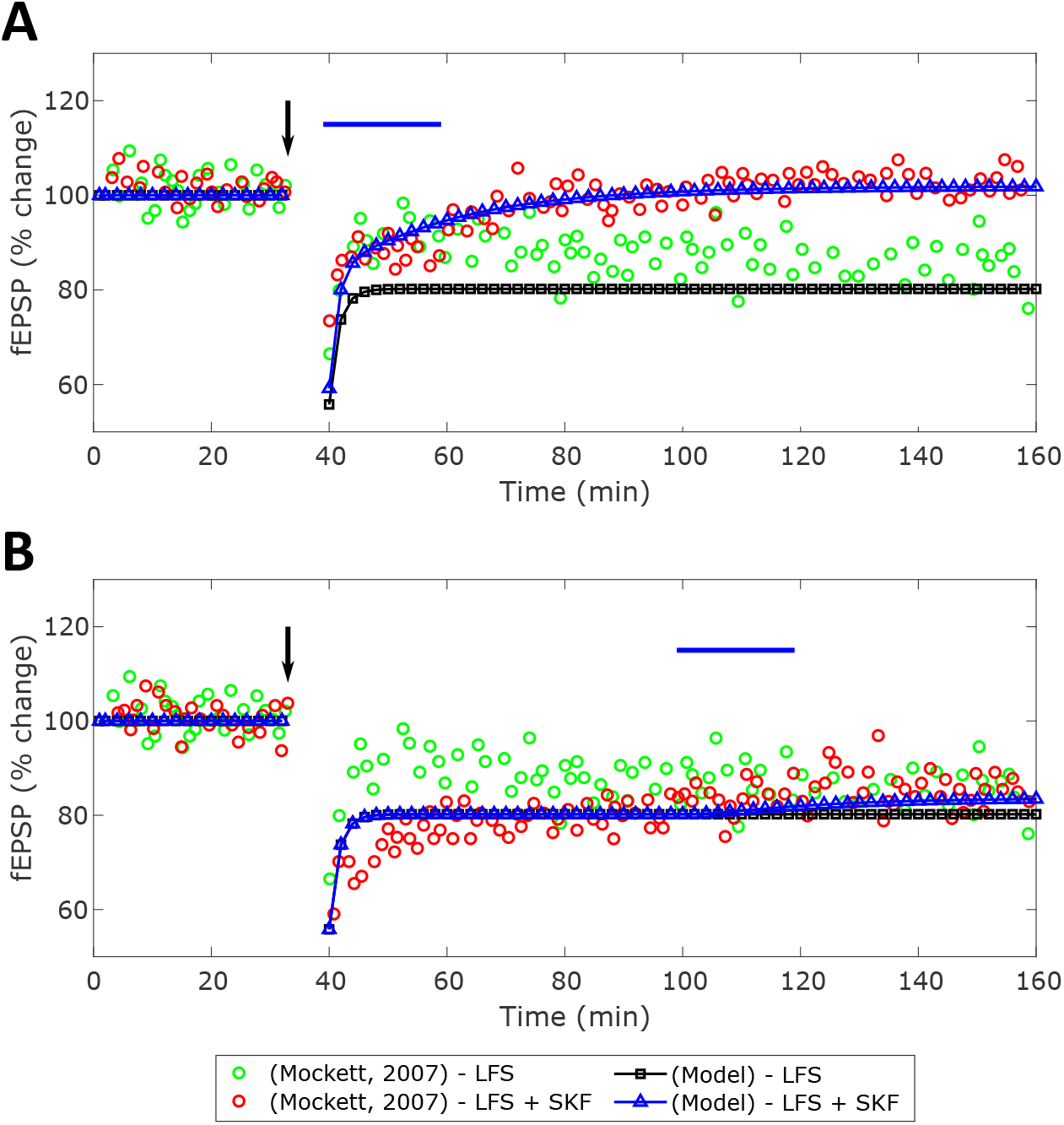
Quantitative comparison between the model predicted and experimentally observed [27] modulation of LFS-induced LTD in the hippocampal SC-CA1 synapse by a D1/D5 agonist SKF 38393. The induced LTD of the SC-CA1 synapse is measured in terms of the percentage (%) change in evoked fEPSP slope from the control. The black-squares represents the application of the LFS protocol of 1200 pulses at 3 *Hz*, while the blue-triangles represent the same LFS protocol in combination with 100 *μM* SKF 38393 for 20 minutes applied at time relative to the LFS protocol (Δ*t* = *t*_*SKF*_ − *t*_*LFS*_). **(A)** shows the LFS-induced LTD without (black-squares) and with (blue-triangles) 100 *μM* SKF 38939 delivered immediately after the LFS protocol. The experimentally reported SKF 38393 enhancement of LTD [27] is shown as the red-circles (Δ*t* = 0 min). **(B)** shows the comparison between the prediction from our model and the experimental data [27] where 100 *μM* SKF 38393 was administered 60 minutes after the same LFS protocol (Δ*t* = 60 min).

After validating our model with the experimental data from Mockett et. al. [27], we used our model to predict the dopaminergic modulation of LFS-induced LTD when SKF 38393 was delivered 100 and 30 minutes before a LFS protocol of 1200 pulses at 3 Hz. Figure 13A shows the prediction from our model when SKF 38393 was delivered 100 minutes before the LFS protocol. As shown in this figure, 100 *μM* of SKF 38393 delivered 100 minutes before the LFS protocol potentiated the SC-CA1 synapse by 48% and converted the LFS-induced LTD into LTP. Then, we applied 100 *μM* of SKF 38393 30 minutes before the same LFS protocol. Figure 13B shows a 30 minute timing between the application of the dopamine agonist and the following LFS protocol flipped the LFS-induced LTD to LTP too, although the overall potentiation was less than if it were applied 100 minutes before the LFS protocol. Our model predictions, if correct, suggest that the application of SKF 38393 before a LFS protocol could potentially convert the LFS-induced LTD into LTP.

**Fig 13.**
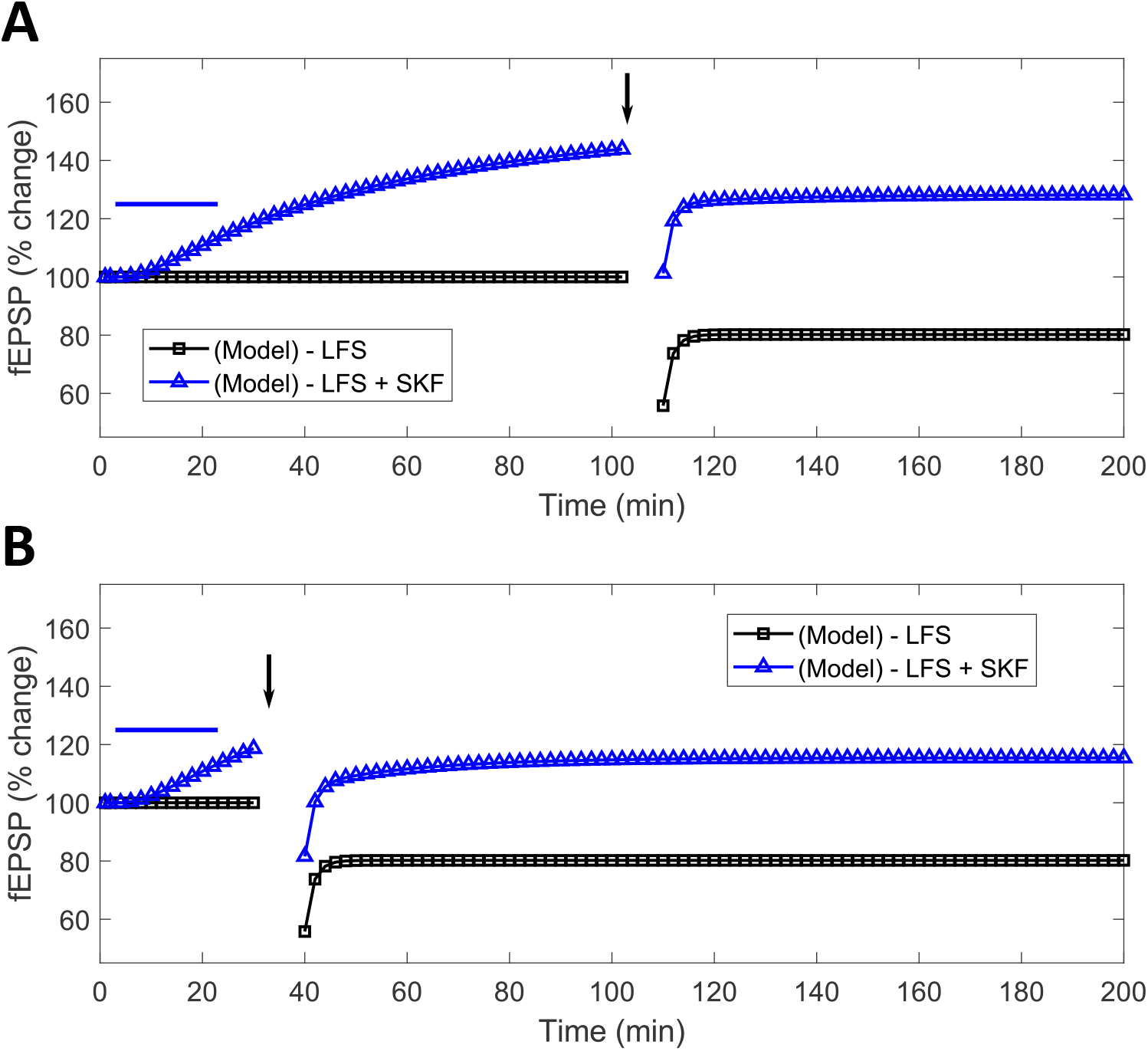
Predictions from our model on the modulation of LFS-induced LTD in the hippocampal SC-CA1 synapse by D1/D5 receptor agonist SKF 38393 when SKF 38393 is delivered prior to the LFS protocol. In these simulation results, 100 *μM* of SKF 38393 is applied 100 **(A)** and 30 minutes **(B)** before a LFS protocol of 1200 pulses at 3 Hz. **(A)** shows the modulation of the LFS-induced LTD by SKF 38393 when SKF 38393 was delivered 100 minutes before the LFS protocol marked as the blue-triangles (Δ*t* = 100). The LTD induced by LFS alone is shown as the black-squares. **(B)** shows the modulation of the LFS-induced LTD by SKF 38393 when SKF 38393 was delivered 30 minutes before the LFS protocol (Δ*t* = 30).

### Dopaminergic modulation of LFS induce LTD predictions with other LFS protocols

In [27], Mockett et al. also investigated whether SKF 38393 could also reverse a strong LFS-induced LTD in SC-CA1 synapses. Specifically, they showed in their experiments that a 20 minutes application of 100 *μM* SKF 38393 immediately after a LFS protocol of 2400 pulses delivered at a frequency of 3 *Hz* potentiated the SC-CA1 synapse by approximately 12% (although no complete reversal of the LFS-induced LTD was observed). Furthermore, the delivery of 100 *μM* SKF 38393 immediately after a LFS protocol of 2 trains of 1200 pulses at 3 *Hz* with a 5 minute intertrain interval potentiated the SC-Ca1 synapse by approximately 8%. The authors stated that the 5 minutes time difference may allow for more intracellular LTD consolidation and become more resistant to any modulation by D1/D5 receptor activation. Therefore, the LFS protocol with the 5 minutes gap between two application of 1200 pulses at 3 *Hz* was modulated less by SKF 38393. From these results, the author’s *in vitro* experimental data suggests that the SKF 38393 mediated changes in the LFS-induced LTD strongly depend on the LFS protocol.

To test the predictive capability of our model, we used our LTD model to see whether our model can predict the experimental results shown by Mockett et al. [27] on the SKF 38393 mediated changes in the LFS-induced LTD under different LFS protocols. We used the same model described in the previous section. First, we applied 100 *μM* SKF 38393 for a 20 minute duration immediately after a LFS protocol of 2400 pulses delivered at 3 *Hz* to our model, identical to the one used in [27]. Our model predicted 14% potentiation of the SC-CA1 synapse as compared to approximately 12% in the experiment (see Figure 14A) and captured the temporal changes mediated by SKF 38393 in the LFS-induced LTD, reasonably well. It should be noted here that we did not use these data to infer our model parameters. Next, we applied 20 minutes of 100 *μM* SKF 38393 immediately after the LFS protocol of 2 trains of 1200 pulses at 3 *Hz*, identical to the one used in [27]. Figure 14B shows the comparison between the prediction from our model and the experimental data from [27]. Our model predicted the observed fEPSP slope changes observed in the experimental data well. In summary, our model quantitively predicted the experimentally observed LFS-induced LTD changes altered by SKF 38393 in SC-CA1 synapses for different LFS protocols.

**Fig 14.**
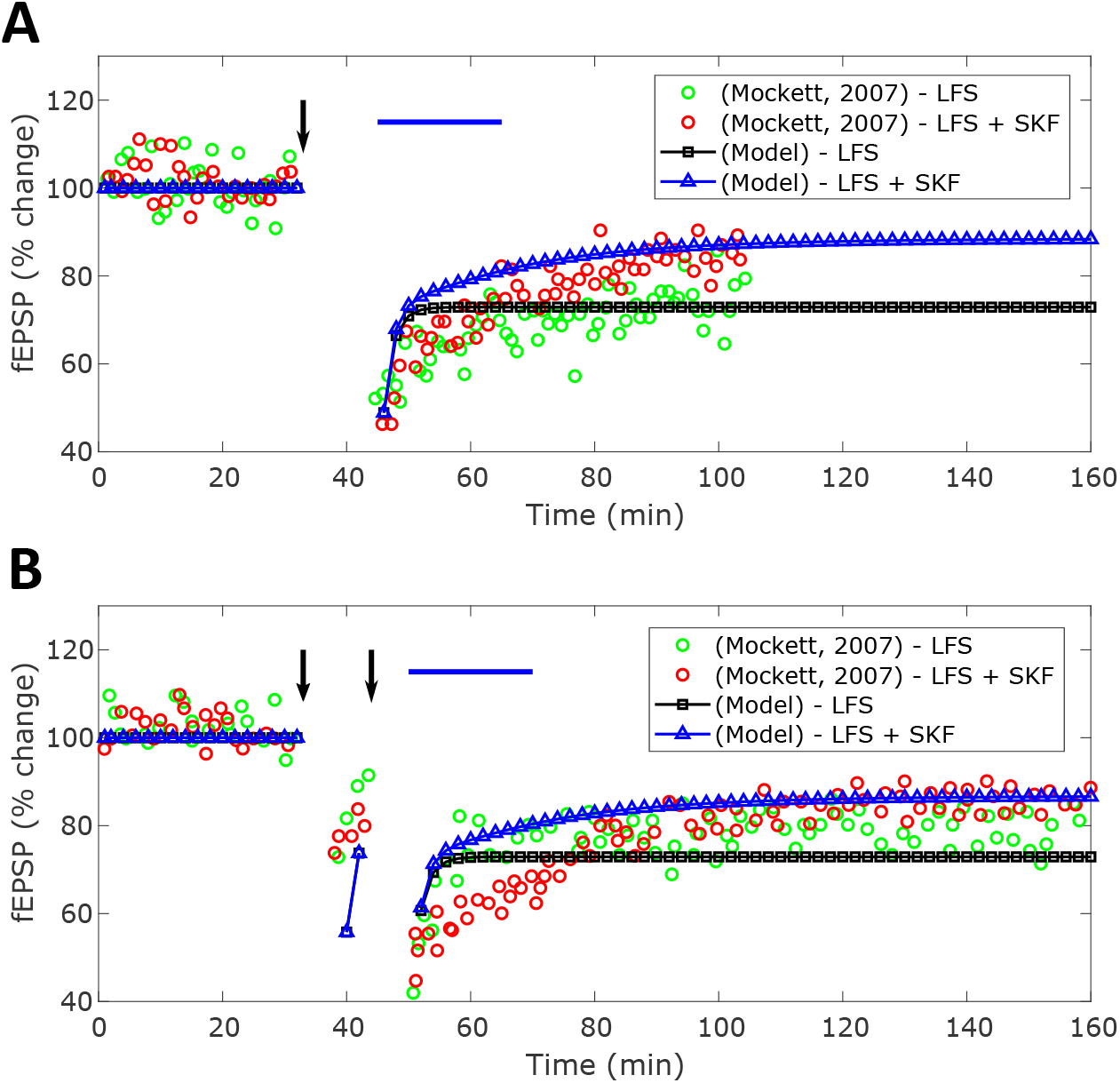
Quantitative comparison between the model predicted and experimentally observed [27] modulation of the LFS-induced LTD under various LFS protocol by SKF 38393. The induced LTD of the SC-CA1 synapse is measured in terms of the percentage (%) change in evoked fEPSP slope from the control. **(A)** shows the comparison between the potentiation of the LFS-induced LTD in the SC-CA1 synapse with the 20 minutes application of 100 *μM* SKF 38393 immediately after the LFS protocol of 2400 pulses at 3 *Hz* reported in the experiment (red-circles) and predicted by our model (blue-triangles). The model predicted LFS-induced LTD by the LFS protocol alone is shown as the black-squares. **(B)** shows the comparison between the potentiation of the LFS-induced LTD in the SC-CA1 synapse when 20 minutes of 100 *μM* SKF 38393 was administered immediately after a LFS protocol of two trains of 1200 pulses at 3 Hz with a 5 minute intertrain interval reported in the experiment (red-circles) and predicted by our model (blue-triangles).

Since our model predicted the experimentally observed modulation of LFS-induced LTD in SC-CA1 synapses by SKF 38393 quantitively under various LFS protocols, we used our model to further investigate the SKF 38393 mediated modulation of SC-CA1 synapses for two different LFS protocols. The first LFS protocol consisted of three trains of 900 pulses at 1 *Hz* with 15 minute intertrain intervals and the second protocol consisted of 900 bursts of three pulses delivered at 1 *Hz*. The rationale for considering these two LFS protocols is that they both induce a similar level of LTD but have a large difference in the duration of the applied LFS protocol. In both cases, we administered 20 minutes of 100 *μM* SKF 38393 to our model immediately after the LFS protocol similar to the protocol used in Mockett et. al. [27] in order to further examine how consolidation of LTD influences the extent SKF 38393 is able to modulate the SC-CA1 LTD. Figures 15A and 15B show the SKF 38393 mediated modulation of the LFS-induced LTD by these two different LFS protocols. The application of 100 *μM* SKF 38393 immediately after three trains of 900 pulses at 1 *Hz* potentiated the SC-CA1 synapse by approximately 8% while SKF 38393 applied immediately after the LFS protocol of 900 bursts at 1 *Hz* potentiated the SC-CA1 synapse by approximately 18%. It should be noted that none of the protocols led to the complete reversal of the LFS-induced LTD. This follows the same trend observed in the experimental data (see Figures 14A and 14B) where the longer the LFS protocol, the less potentiation induced by SKF 38393. Mockett et. al. [27] hypothesized that longer LFS protocols may allow the LFS-induced LTD more time to consolidate, which would reduce the ability of the dopamine agonist to modulate the LTD. If this hypothesis is true, then the release of SKF 38393 immediately after two different LFS protocols with dramatically different stimulation times would have more obviously different effects on the dopaminergic modulation of LFS-induced LTD for each protocol, as predicted by our model (see Figure 15).

**Fig 15.**
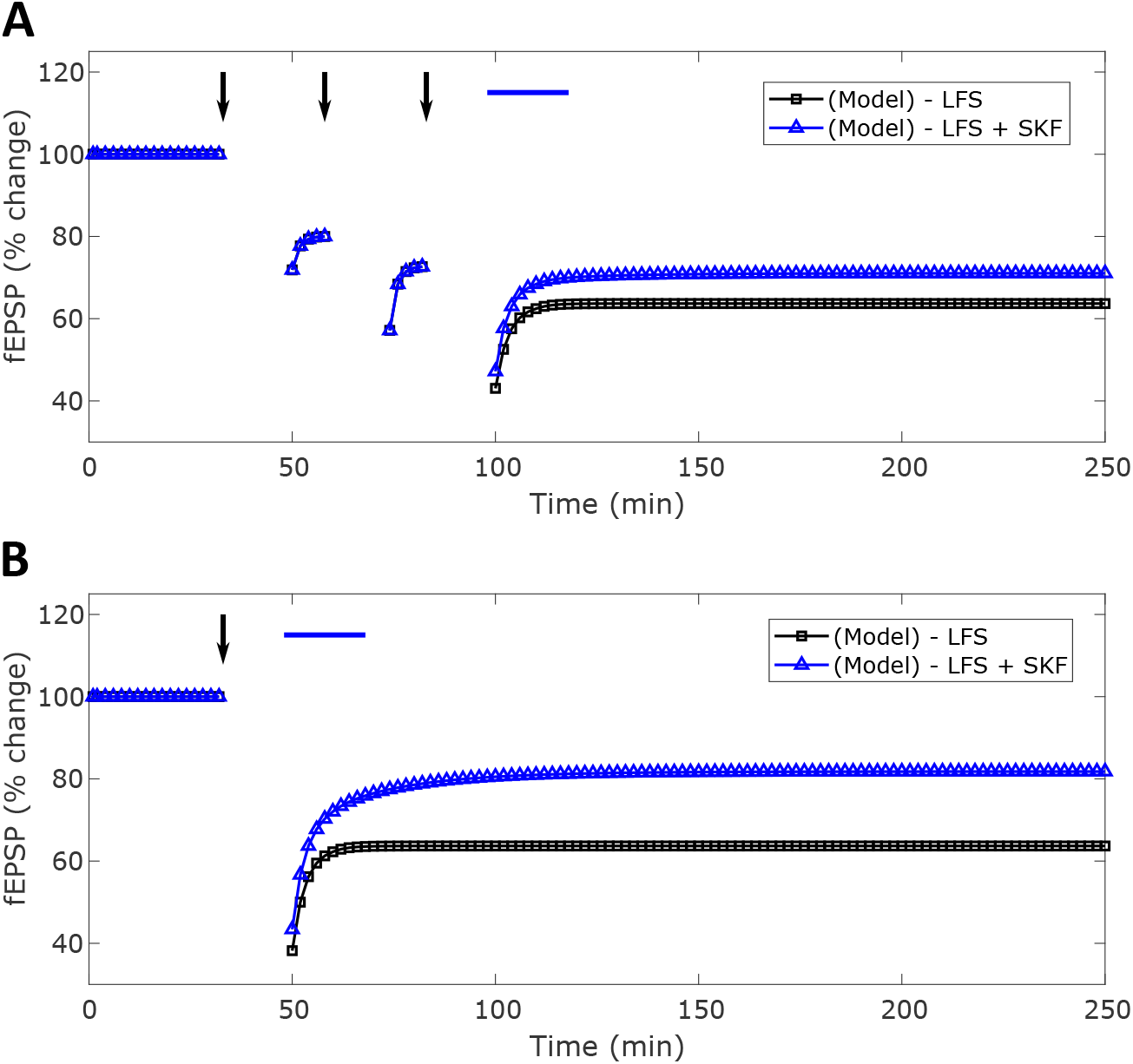
Predictions from our model on the modulation of the LFS-induced LTD under various LFS protocol by SKF 38393. 100 *μM* SKF 38393 was applied for 20 minutes immediately after two different LFS protocols. **(A)** shows the model predicted enhancement in the LFS-induced LTD by SKF 38393 for a LFS protocol of 3 trains of 900 pulses at 1 *Hz* with 10 minute intertrain intervals. **(B)** shows the model predicted enhancement in the LFS-induced LTD by SKF 38393 for a LFS protocol of 900 bursts at 1 *Hz* where each burst consists of 3 pulses delivered at 20 *Hz*. The SKF 38393 modulated LFS-induced LTD is shown as blue-triangles and the LFS-induced LTD in the absence of SKF 38393 is denoted as the black-squares.

### Antagonizing the D1/D5 receptor blocks LFS induced late-LTD

In this section, we investigated the capability of our model in predicting experimental data on the modulation of SC-CA1 long-term synaptic plasticity by the simultaneous application of the D1/D5 receptor antagonist SCH 23390 and a LFS protocol. In *in vitro* experiment, Frey et. al. [8] observed that the application of the D1/D5 receptor antagonist SCH 23390 blocked consolidation of LFS-induced LTD in SC-CA1 synapses. They applied 0.1 *μM* SCH 23390 for 60 minutes starting 30 minutes before the LFS protocol of 900 bursts of 3 pulses at 1 *Hz* and observed the reversal of LFS-induced LTD back to the baseline over a period of 450 minutes. Since the blockage of basal levels of dopamine binding to D1/D5 receptors with SCH 23390 blocked the consolidation of LTD, Frey [8] hypothesized that dopamine is required for the induction of late-LTD.

We used our model (see Eqs. (1),(6a) -(6d), (6f) - (6i), (15a) -(15r), and (18)) to predict the experimental data from [8] under the same protocol, quantitatively. We incorporated the effect of SCH 23390 in our model through a model parameter *k*_*basal*_. This parameter takes a non-zero value in the presence of SCH 23390 (see Eq.(15o)). Figure 16 compares the prediction from our model with the experimental data from Frey et. al. [8]. As shown in this figure, our model predicts the reversal of the LFS-induced LTD in the presence of SCH 23390, as observed in the experiment.

**Fig 16.**
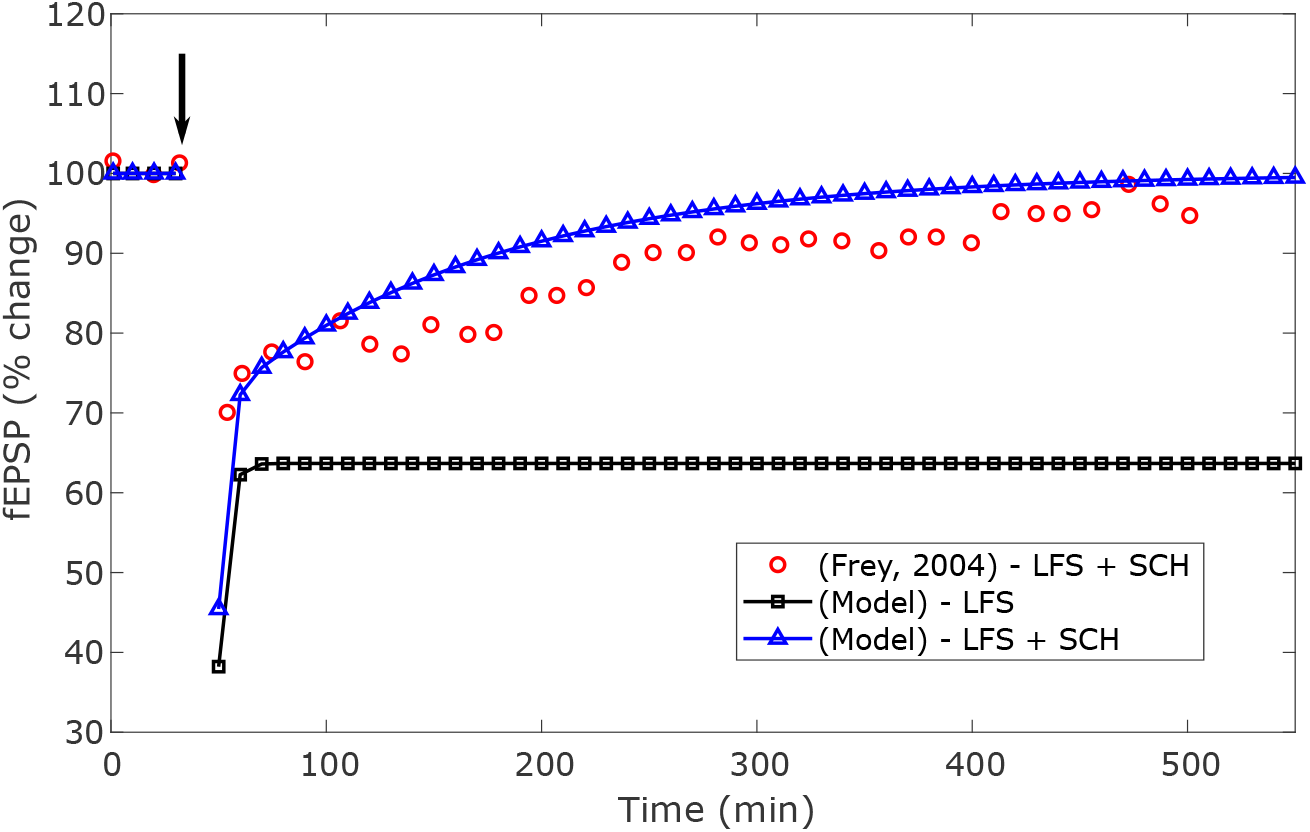
Quantitative comparison between the model predicted and experimentally observed [8] modulation of LFS-induced LTD in the SC-CA1 synapse in the presence of a D1/D5 receptors antagonist SCH 23390. The D1/D5 receptors were blocked with the application of 0.1 *μM* SCH 23390 during a LFS protocol of 900 bursts of 3 pulses at 1 *Hz* where the 3 pulses of each burst were delivered at 20 *Hz* (black-arrow). The slow reversal of the LFS-induced LTD observed in the experiment and predicted by our model are shown as the red-circles and the blue-triangles, respectively. The LTD induced by LFS alone is denoted by the black-squares.

### Concentration dependent spatiotemporal modulation of LFS-induced LTD by SKF 38393

In this section, we used our model to investigate the effect of the concentration of SKF 38393 and its application time relative to the LFS protocol on the modulation of LFS-induced LTD in SC-CA1 synapses. Experimental data from *in vitro* hippocampal slice experiments have shown that the modulation of a LFS-induced LTD in SC-CA1 synapses strongly depends on the applied concentration and timing of SKF 38393 relative to the LFS protocol. Particularly, Chen et. al. [39] showed that the application of 3 *μM* of SKF 38393 during a LFS protocol of 450 pulses at 1 *Hz* enhanced the induced LTD by approximately 18% (see Figure 17). This result highlights a different role that a dopamine agonists can play if applied at a low concentration. Furthermore, the experimental observation was consistent with the observed slow-onset depotentiation of the SC-CA1 synapse by the application of low concentrations of dopamine alone [8].

**Fig 17.**
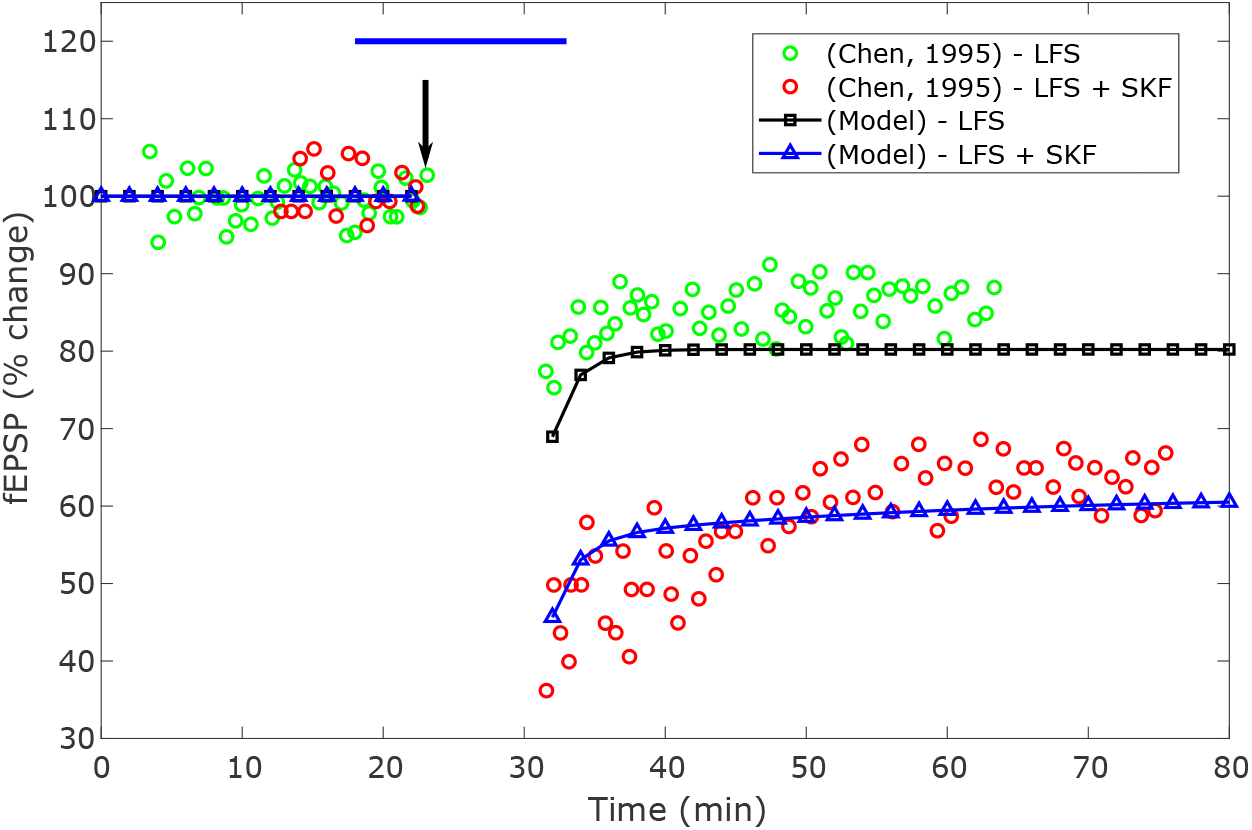
Quantitative comparison between the model predicted and experimentally observed [39] modulation of LFS-induced LTD in the SC-CA1 synapse by SKF 38393 applied at a low concentration. A low concentration of 3 *μM* SKF 38393 was applied for a 15 minute duration (blue-bar) and 5 minutes before a LFS protocol of 450 pulses at 1 *Hz* (black-arrow). The experimentally observed and model predicted changes in the LFS-induced LTD by SKF 38393 are shown as the red-circles and blue-triangles, respectively. The LTD induced by LFS alone in the experiment and predicted by our model are denoted as the green-circles and black-squares, respectively.

To investigate the capability of our model in predicting the observed depotentiation of SC-CA1 synapse at a low concentration of SKF 38393 by Chen et.al. [39], we applied the same low concentration of SKF 38393 to our model (see Eqs. (1),(6a) -(6d), (6f) - (6i), (15a) -(15r), and (18) in Materials and methods section) under the same LFS protocol. Figure 17 compares the prediction from our model with the experimental data from [39]. As shown in this figure, our model predicts the experimental data on the SKF 38393 modulated LFS-induced LTD at a low concentration quantitatively (enhancement in the LFS-induced LTD by approximately 20% an hour after the LFS protocol). Additionally, our model captures the temporal dynamics of these modulations.

After validating our model with the limited experimental data, we further used our model to investigate how various concentrations and timing of the application of SKF 38393 relative to a LFS protocol modulates the LFS-induced LTD in SC-CA1 synapses. We applied seven different concentrations (1 – 100 *μM*, 20 minutes duration) of SKF 38393 at various times before and after a LFS protocol of 1200 pulses at 3 *Hz*. Figures 18A, 18B, and 18C show the predictions from our model on the SKF 38393 mediated modulation of the LFS-induced LTD for timings of SKF 38393 applied 212, 30, and 20 minutes, respectively, before the LFS protocol. Here, SKF 38393 alone induced a slow onset potentiation of the SC-CA1 synapse that plateaued after the LFS protocol in Figure 18A. In Figure 18B, the LFS protocol applied 30 minutes after the administration of SKF 38393 reduced the concentration dependent slow-onset potentiation by SKF 38393, since there was less time for the consolidation of the SKF 38393 induced potentiation. After the LFS protocol, the SKF 38393 induced potentiation remained at the saturation level. The application of SKF 38393 20 minutes before the LFS protocol highlighted the concentration dependent bifurcation of the SKF 38393 mediated modulation of the LFS-induced LTD, similar to the LTP case (see Figure 8C). Particularly, low concentrations of SKF 38393 enhanced the depotentiation of the SC-CA1 synapse, while high concentrations of SKF 38393 reversed LFS induced-LTD to varying degrees.

**Fig 18.**
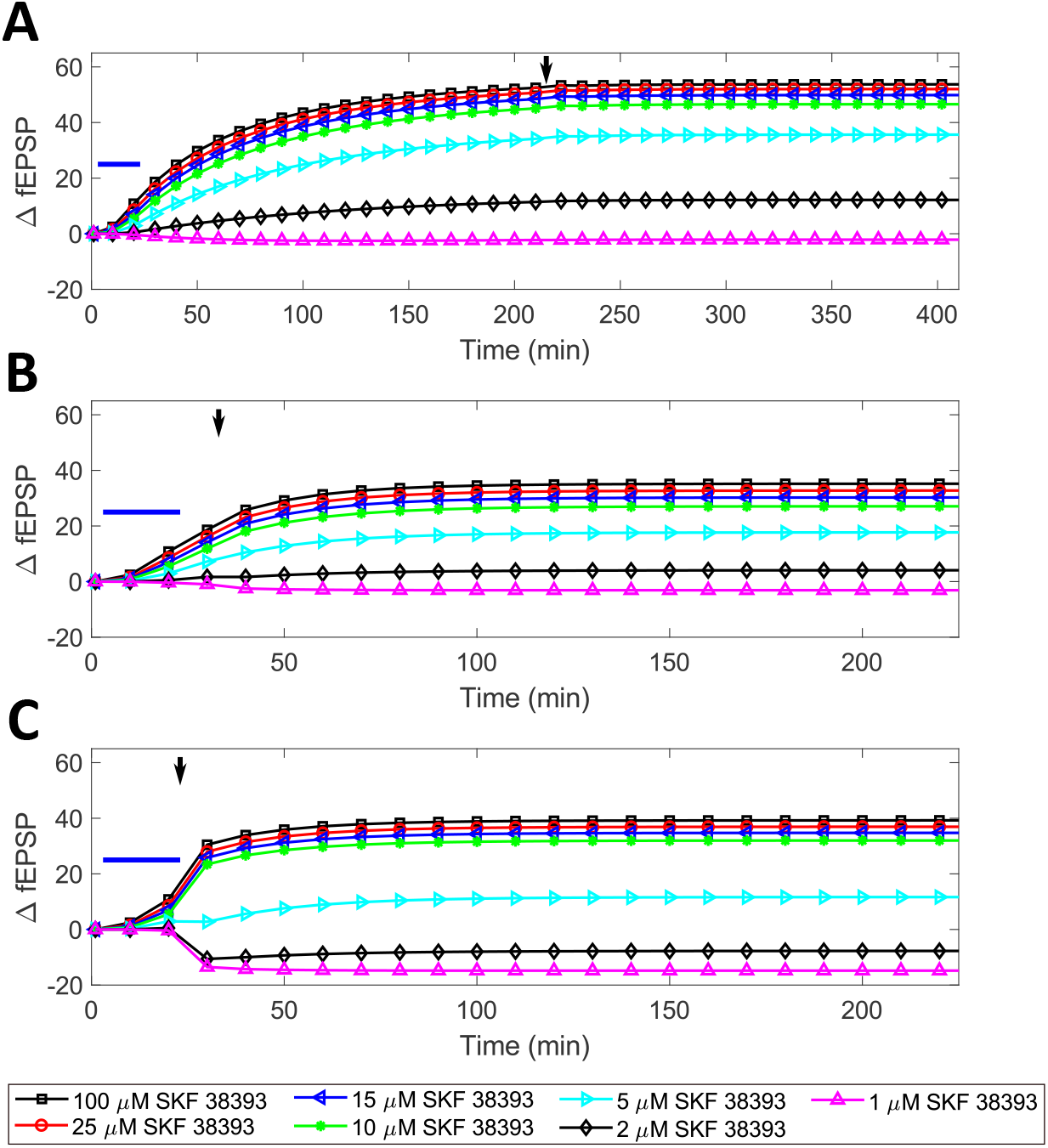
Model predictions on the effect of the concentration and timing of SKF 38393 relative to the LFS protocol (1200 pulses delivered at 3 *Hz*) in modulating the LFS-induced LTD when SKF 38393 was applied before the LFS protocol for 20 minutes. SKF 38393 (blue-bar) was delivered **(A)** 212 minutes (Δ*t* = 212 min), **(B)** 30 minutes (Δ*t* = 30 min), and **(C)** 20 minutes (Δ*t* = 20 min) before the LFS protocol (black-arrow).

Next, we used our model to investigate how various concentrations of SKF 38393 modulate the LFS-induced LTD in SC-CA1 synapses when SKF 38393 is applied at different times relative to LFS but after the LFS protocol. Figures 19A, 19B, and 19C show the prediction from our model when we applied SKF 38393 10, 30, and 60 minutes after the same LFS protocol (1200 pulses at 3 *Hz*). As shown in these figures, the concentration dependent potentiation by SKF 38393 decreased for the same given concentration as the relative timing between the application of SKF 38393 and the LFS increased. Moreover, the potentiation by SKF 38393 was much smaller compared to when the SKF 38393 was applied before the LFS protocol to induce LTD, which is consistent with results in the previous sections.

**Fig 19.**
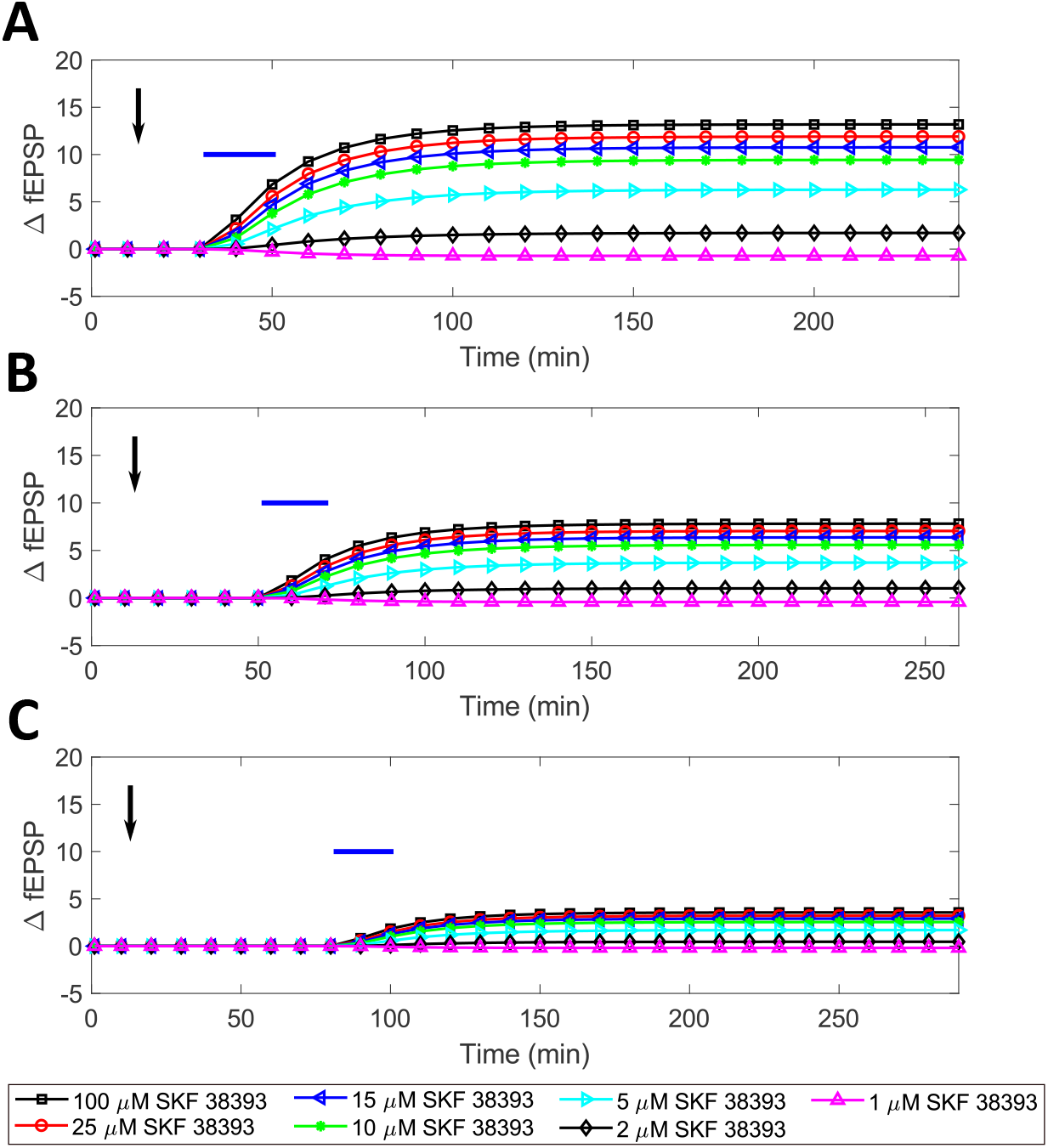
Model predictions on the effect of the concentration and timing of SKF 38393 relative to the LFS protocol (1200 pulses delivered at 3 *Hz*) in modulating the LFS-induced LTD when SKF 38393 was applied after the LFS protocol for 20 minutes. SKF 38393 (blue-bar) was delivered **(A)** 10 minutes, **(B)** 30 minutes, and **(C)** 60 minutes after the LFS protocol (black-arrow).

## Discussion

In this paper, we modeled the dose-dependent modulation of high/low-frequency stimulation (HFS/LFS) induced long-term potentiation/depression (LTP/LTD) of a Schaffer collateral – CA1 pyramidal neuron (SC-CA1) synapse by three dopamine D1/D5 receptor agonists, SKF 38393, dopamine and 6-bromo-APB. Our model predicted the experimentally observed temporal effects of these D1/D5 agonists’ concentrations, and the relative time between the applied agonist and the HFS/LFS induction on both the early and the late LTP/LTD, quantitatively. Specifically, we derived the following conclusions from our modeling results:

1. For a given concentration and duration of a dopamine D1/D5 receptors agonist, the maximum change in the late LTP/LTD occurred when the agonist was delivered a long-time (approximately 200 minutes) before the HFS/LFS induction, which is consistent with the experimental results from Huang et.al. [2].
2. For a given concentration and duration of a dopamine D1/D5 receptors agonist, the maximum change in the late LTP/LTD decreased slowly with the decrease in the time difference between the application of the agonist and the HFS/LFS protocol when the agonist was applied before the HFS/LFS protocol but not immediately before or during the HFS/LFS protocol. The conclusion is consistent the experimental data available in the literature [2, 34].
3. For a given concentration and duration of a dopamine D1/D5 receptors agonist, the maximum change in the late LTP/LTD quickly decreased with the increase in the time gap between the application of the dopamine agonist and the HFS/LFS protocol when the agonist was applied after the HFS/LFS protocol. The conclusion is consistent with the experimental data available from [27].
4. For a given concentration and duration of a dopamine D1/D5 receptors agonist, application of the agonist immediately before or during an HFS/LFS protocol enhanced the dopaminergic modulation of the SC-CA1 synapse in a highly nonlinear fashion (sharp changes), which is consistent with the experimental data from [39, 40].
5. For a given concentration and duration of a dopamine D1/D5 receptors agonist, high concentrations of the agonist injected immediately before or during a HFS/LFS protocol modulated SC-CA1 synaptic plasticity by further potentiating the HFS/LFS induced potentiation/depotentiation. This result is consistent with the experimentally observed data from [40].
6. For a given concentration and duration of a dopamine D1/D5 receptors agonist, low concentrations of the agonist injected immediately before or during a HFS/LFS protocol modulated SC-CA1 synaptic plasticity by further depotentiating the HFS/LFS induced potentiation/depotentiation. This is consistent with the limited experimental data from [39].
7. For a fixed duration of the applied dopamine D1/D5 receptors agonist and a large time gap between the dopamine agonist and the HFS/LFS protocol, the dopaminergic mediated modulation of LTP/LTD increased with an increase in the concentration level of the applied agonist.

We estimated our LTP model parameters using the available experimental data (i.e., the delivery of 50 *μM* concentration of SKF 38393 for 15 minutes more than two hours before the HFS-induced LTP induction [34] and the delivery of 5 *μM* concentration of 6-bromo-APB immediately before a weak HFS protocol for 5 minutes [40]). We then used our model to make predictions on the D1/D5 mediated enhancements of the late LTP when SKF 38393 was administered at various concentrations closer in time relative to the HFS protocol. We found that the injection of a high concentration of SKF 38393 immediately before the HFS protocol non-linearly modulated the HFS-induced LTP in the SC-CA1 synapse by further potentiating the HFS-induced potentiation, while the injection of a low concentration of SKF 38393 depotentiated the HFS-induced LTP. Based on the current experimental results in the literature, it is not clear if such a concentration dependent bifurcation should happen. We believe that additional experiments, such as the application of a high (50 *μM*) and a low (1 *μM*) concentration of SKF 38393 immediately before a HFS protocol of 3 trains of 100 pulses at 100 *Hz*, will clarify this observation.

Similarly, we estimated our LTD model parameters using the limited available experimental data, i.e., the delivery of 100 *μM* concentration of SKF 38393 for 20 minutes immediately and 60 minutes after a LFS protocol [27]. Based on this, our LTD model predicted similar trends in the D1/D5 mediated enhancements in the recorded fEPSP slope, as observed in our LTP model. Based on the results from our model, one particular instance we believe would be useful to further validate our model is to perform an *in vitro* hippocampal slice experiment by administering 100 *μM* SKF 38393 100 minutes before a LFS protocol. If the predicted outcome from our model is correct, we will expect that the administration of SKF 38393 100 minutes before the LFS protocol will convert the LFS-induced LTD into LTP. Additionally, we believe that another experiment, such as the application of a high (100 *μM*) and low (2 *μM*) concentration of SKF 38393 injected immediately before a LFS protocol, will clarify the experimental observation by Chen [39] (see Figure 17) and validate our model predictions.

Our modeling approach to integrate the LTP/LTD effects mediated by HFS/LFS and D1/D5 agonists is based on the hypothesis that there exists limited resources for the consolidation of the HFS/LFS-induced LTP/LTD into the late-LTP/LTD. We formulated this hypothesis based on the limited experimental data from [2, 27]. Particularly, it was shown in an *in vitro* hippocampal slice experiment that the application of 100 *μM* of SKF 38393 immediately after the LFS protocol reversed the consolidation of the late-LTD, whereas the application of 100 *μM* of SKF 38393 an hour after the LFS protocol produced no significant reversal of the late-LTD consolidation [27]. Similarly, the application of 50 *μM* SKF 38393 200 minutes before HFS induction resulted in a substantial magnitude change of HFS induced LTP, while the delivery of 50 *μM* SKF 38393 50 minutes after resulted in no enhancement of HFS induced LTP [2]. These experimental observations highlight when the HFS/LFS protocol induces a change in the synaptic conductance, regardless if the conductance is increased or decreased, it impedes the D1/D5 receptors mediated modulation of the SC-CA1 synaptic LTP/LTD more as the administration of a dopamine D1/D5 agonist is delayed further relative to the HFS/LFS protocol. We believe that it would be worth performing additional experiments that would support the limited resource hypothesis by applying a dopamine agonist immediately before two different LFS protocols that induce similar levels of LTD but have drastically different stimulation durations, such as the application of 100 *μM* SKF 38393 immediately after three trains of 900 pulses at 1 *Hz* (see Figure 15A) and immediately after the a LFS protocol of 900 bursts of 3 pulses at 1 *Hz* (see Figure 15B). This will not only validate our modeling hypothesis of the limited resources but support many predictions from our model.

We have developed our modeling framework in a way so that any part of the model can be swapped out if an improved model becomes available. For example, if a better HFS/LFS model becomes available, the new HFS/LFS model could replace the frequency dependent plasticity dynamics in Eqs. (6a)- (6i) without making substantive changes in the entire model. In order to fully incorporate the new model’s dynamics into the rest of the model, the output of the frequency dependent model to the other segments of our SC-CA1 model must be updated, as well. Therefore, the derivative of the output of the new HFS/LFS model, which determines the SC-CA1 synaptic plasticity change, must be updated in Eqs. (15i), (15j), (15n), and (15o). Additionally, our model could possibly be used to make prediction about the dopaminergic modulation of STDP by swapping out the frequency dependent plasticity model for a STDP model. The rest of the parameters would then need to be re-fit to the STDP specific experimental data. Finally, the dopaminergic model could be expanded to other dopamine agonists if the concentration dependent slow-onset potentiation experimental data becomes available. The inherent segmented nature of our model makes it more flexible and may allow incorporation of different types of plasticity and neuromodulators to be considered in the future.

Our model assumes that SKF 38393 can induce a slow-onset-potentiation in the SC-CA1 synapses in the absence of an LTP/LTD stimulation protocol (HFS/LFS). Although we found some controversy on this in the literature [1–4, 27], recent experiments on applying various concentration levels of SKF 38393 in the absence of any HFS/LFS protocol [4, 47] support our modeling assumption. While the lack of sufficient data may limit the quality of our model, it still provides important insights into the gaps in our current understanding. Moreover, it suggests specific experiments that we believe will help to fill in the gaps in our understanding of the spatiotemporal modulation of SC-CA1 synapses by dopamine.

## Materials and methods

### Hippocampal CA1 pyramidal neuron model

We used an experimentally validated single compartment Hodgkin-Huxley model [29] to represent the CA1 pyramidal neuron dynamics. The details of the model can be found in [29]. Here, we briefly described the model. The time evolution of the membrane potential, *V* , is governed by the sum of various ionic currents, as given by Eq. (1).

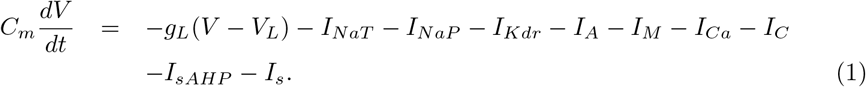

In Eq. (1), *I*_*NaT*_ and *I*_*NaP*_ are the transient and persistent sodium (*Na*^*+*^) currents, respectively. *I*_*Kdr*_ is the delayed rectifier potassium (*K*^*+*^) current, and *I*_*M*_ is the muscarinic-sensitive *K*^*+*^ current. *I*_*A*_ is the A-type *K*^*+*^ current. *I*_*Ca*_ is the high threshold calcium (*Ca*^*2+*^) current, and *I*_*C*_ is the *Ca*^*2+*^-activated *K*^*+*^ current for the rapid spike repolarization. *I*_*sAHP*_ is the slow *Ca*^*2+*^-activated *K*^*+*^ current responsible for the slow after hyperolarization and the spike frequency adaptation. *I*_*s*_ is the synaptic current to the CA1 pyramidal neuron from the Schaffer collateral-CA1 synapse, which we described in the next section. *C*_*m*_, *g*_*L*_, and *V*_*L*_ are the membrane capacitance, the leaky current conductance, and the leaky current reversal potential, respectively.

The ionic currents in Eq. (1) are voltage-dependent, and we provided their functional forms in Eqs. (2a)-(2h).

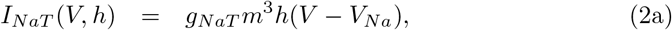

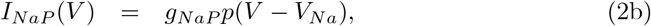

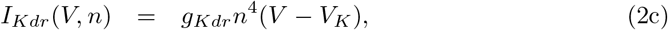

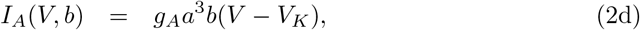

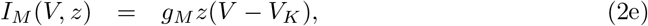

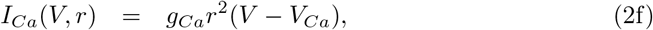

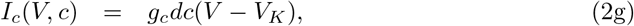

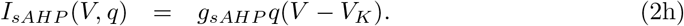

For a given ionic current *I*_*i*_ where *i* ∈ {*NaT, NaP, Kdr, A, M, Ca, c, sAHP*} , *g*_*i*_ is the maximum conductance of the *i*^*th*^ ionic channel type. *V*_*Na*_, *V*_*K*_, and *V*_*Ca*_ are the reversal potential of the sodium, potassium, and calcium channels, respectively. The dynamics of the gating variables (*m, h, p, n, a, b, z, r, d, c, q*) are described by Eqs. (3a)-(3q). Their activation and deactivation dynamics are described by either a differential equation, a generic activation/inactivation function *x*_∞_(*V*) = {1 + *exp*[−(*V − θ*_*x*_)]/*σ*_*x*_}^−1^, or a combination of both. The dynamics of the *Ca*^*2+*^ activated ion channel gating variables *d* and *q* depend on the intracellular *Ca*^*2+*^ concentrations. We provided the governing equation of the *Ca*^*2+*^ dynamics in Eq. (3r), where *v* is the rate of *Ca*^*2+*^ into the neuron and *τ*_*Ca*_ is the *Ca*^*2+*^ decay constant.

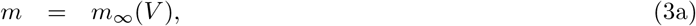

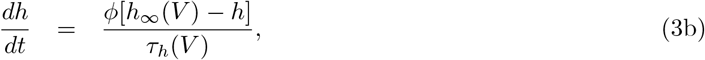

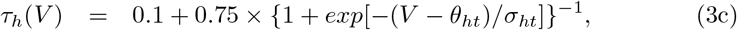

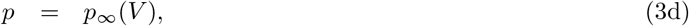

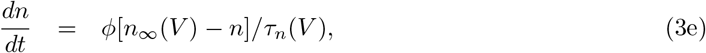

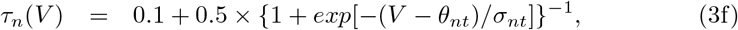

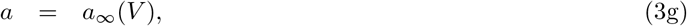

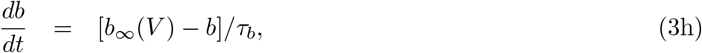

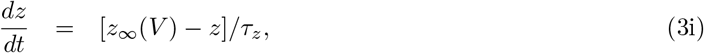

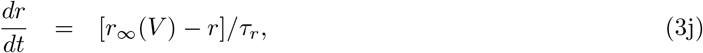

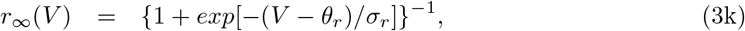

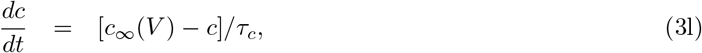

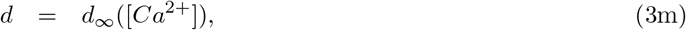

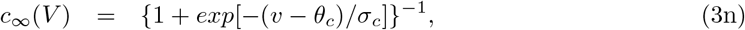

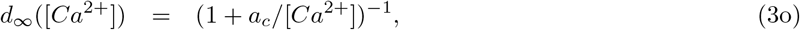

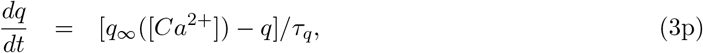

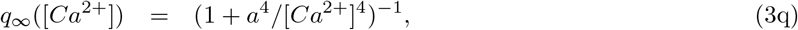

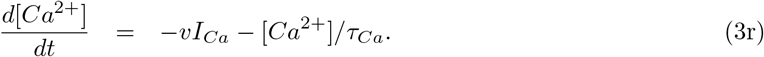

We provided the CA1 pyramidal neuron model parameters used in this paper in Table 1.

**Table 1.** CA1 Pyramidal Neuron Model Parameters [29].

| Parameter | Value | Parameter | Value |
| --- | --- | --- | --- |
| $C$ | 1 $\mu\text{F}/\text{cm}^2$ | $\theta_m$ | -30 mV |
| $g_L$ | 0.05 $\text{mS}/\text{cm}^2$ | $\sigma_m$ | 9.5 mV |
| $g_{NaT}$ | 35 $\text{mS}/\text{cm}^2$ | $\theta_h$ | -45 mV |
| $g_{NaP}$ | 0.3 $\text{mS}/\text{cm}^2$ | $\sigma_h$ | -7 mV |
| $g_{Kdr}$ | 6 $\text{mS}/\text{cm}^2$ | $\theta_{ht}$ | -40.5 mV |
| $g_A$ | 1.4 $\text{mS}/\text{cm}^2$ | $\sigma_{ht}$ | -6 mV |
| $g_M$ | 1 $\text{mS}/\text{cm}^2$ | $\phi$ | 1 |
| $g_{Ca}$ | 0.08 $\text{mS}/\text{cm}^2$ | $\theta_p$ | -46 mV |
| $g_c$ | 10 $\text{mS}/\text{cm}^2$ | $\sigma_p$ | 3 mV |
| $g_{sAHP}$ | 5 $\text{mS}/\text{cm}^2$ | $\theta_n$ | -35 mV |
| $V_L$ | -70 mV | $\sigma_n$ | 10 mV |
| $V_{Na}$ | 55 mV | $\theta_{nt}$ | -27 mV |
| $V_K$ | -90 mV | $\sigma_{nt}$ | -15 mV |
| $V_{Ca}$ | 120 mV | $\theta_a$ | -50 mV |
| $\nu$ | 0.13 $\text{cm}^2/(\text{ms} \times \mu\text{A})$ | $\sigma_a$ | 20 mV |
| $\tau_{Ca}$ | 13 ms | $\theta_b$ | -80 mV |
| $\sigma_b$ | -6 mV | $\theta_z$ | -39 mV |
| $\sigma_z$ | 5 mV | $\tau_b$ | 15 |
| $\tau_z$ | 75 | $\tau_r$ | 1 |
| $\theta_r$ | -20 mV | $\sigma_r$ | 10 mV |
| $\tau_c$ | 2 | $\theta_c$ | -30 mV |
| $\sigma_c$ | 7 mV | $a_c$ | 6 |
| $\tau_q$ | 450 | $a_q$ | 2 |

### Schaffer collateral - CA1 pyramidal neuron synaptic dynamics

To represent the dynamics of the synaptic current, *I*_*s*_, from the Schaffer collateral fiber to the CA1 pyramidal neuron (see Eq. (1)), we used a phenomenological synaptic model [30], described by Eqs. (4a) -(4b). The model captures the synaptic contributions from the AMPA and NMDA neurotransmitters.

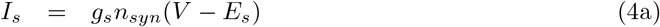

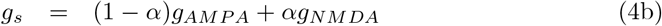

In Eq. (4a), *g*_*s*_, *V* , and *E*_*s*_ are the synaptic conductance, the postsynaptic CA1 pyramidal neuron membrane potential, and the synaptic reversal membrane potential, respectively. The synaptic conductance *g*_*s*_ (see Eq. (4b)) is the weighted sum of the AMPA (*g*_*AMPA*_) and NMDA (*g*_*NMDA*_) conductances, where *α* determines the relative contribution of the AMPA and NMDA conductances. We presented a dynamical model of the AMPA and NMDA conductances in Eqs. (4c)-(4h).

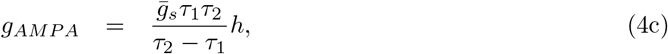

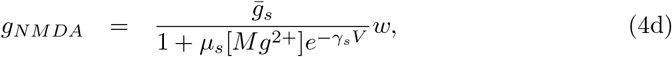

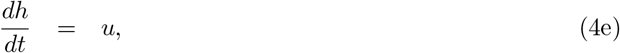

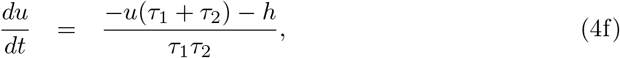

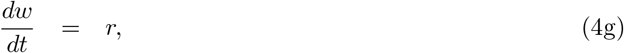

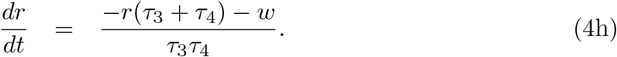

In Eqs. (4c)-(4h), 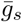 is the maximum synaptic conductance of the Schaffer collateral - CA1 pyramidal neuron (SC-CA1) synapse, measured in the presence of AMPA only receptors. The gating variables *h* and *w* gate the AMPA and NMDA conductances, respectively. *τ*_1_ and *τ*_2_ are the AMPA ionotropic receptors’ rise and decay time constants, respectively. Similarly, *τ*_3_ and *τ*_4_ are the NMDA ionotropic receptors’ rise and decay time constants, respectively. The NMDA conductance *g*_*NMDA*_ is dependent on both the postsynaptic membrane potential *V* and the magnesium concentration [*Mg*^2+^]. *μ*_*s*_ and *γ*_*s*_ are scaling parameters.

The complete synaptic dynamics of the SC-CA1 synapse described by Eqs. (4a)-(4h) is governed by a set of model parameters 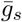, *τ*_1_, *τ*_2_, *τ*_3_, *τ*_4_, *μ*_*s*_, *γ*_*s*_, [*Mg*^2+^], *α, E*_*s*_. We fixed the model parameters [*Mg*^2+^], *α*, and *E*_*s*_ to 1 *μM* , 0.1, and 0 *mV* , respectively [30]. Then we inferred the remaining model parameters using an approximate Bayesian inference approach based on the sequential Monte Carlo (ABC-SMC) [33] from experimental data of AMPA and NMDA excitatory postsynaptic current induced by stimulating the SC-CA1 synapse with a brief electrical pulse [48]. Briefly, we first fitted the parameters 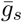, *τ*_1_, and *τ*_2_ with the ABC-SMC approach using a sum of the squared errors distance function shown in Eq. (5c) between the AMPA current (Eq. (5a)) in our model and the experimental data available from [48]. Then, we fitted the parameters *τ*_3_, *τ*_4_, *μ*_*s*_, and *γ*_*s*_ using a distance function that measured the distance between the NMDA current (Eq. (5b)) in our model and the data from [48]. We provided the inferred model parameters in Table 2 and Figure 20 shows the histograms representing the approximate posterior distributions for each of the parameters.

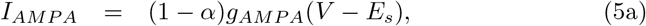

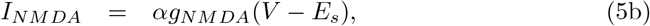

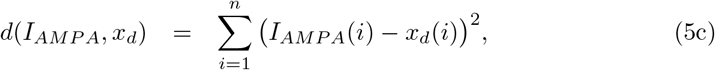

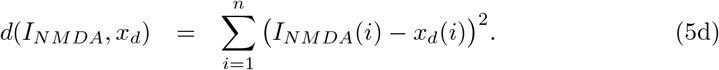

**Table 2.** SC-CA1 synaptic dynamics parameters.

| Parameter | Value | Reference |
| --- | --- | --- |
| $[Mg^{2+}]$ | $1 \mu M$ | [30] |
| $\alpha$ | 0.1 | [30] |
| $E_s$ | $0 mV$ | [30] |
| $\eta_{syn}$ | 17.9 | [30, 48] |

**Table 3.** Inferred SC-CA1 synaptic dynamics parameters.

| Parameter | Value |
| --- | --- |
| $\bar{g}_s$ | $4.981 \times 10^{-3} \pm 0.7 \times 10^{-3} mS$ |
| $\tau_1$ | $0.8263 \pm 0.17 ms$ |
| $\tau_2$ | $4.548 \pm 0.74 ms$ |
| $\tau_3$ | $0.8189 \pm 0.49 ms$ |
| $\tau_4$ | $74.788 \pm 12 ms$ |
| $\mu_s$ | $0.2866 \pm 0.096 \mu M^{-1}$ |
| $\gamma_s$ | $1.3 \times 10^{-2} \pm 0.52 \times 10^{-2} mV^{-1}$ |

**Fig 20.**
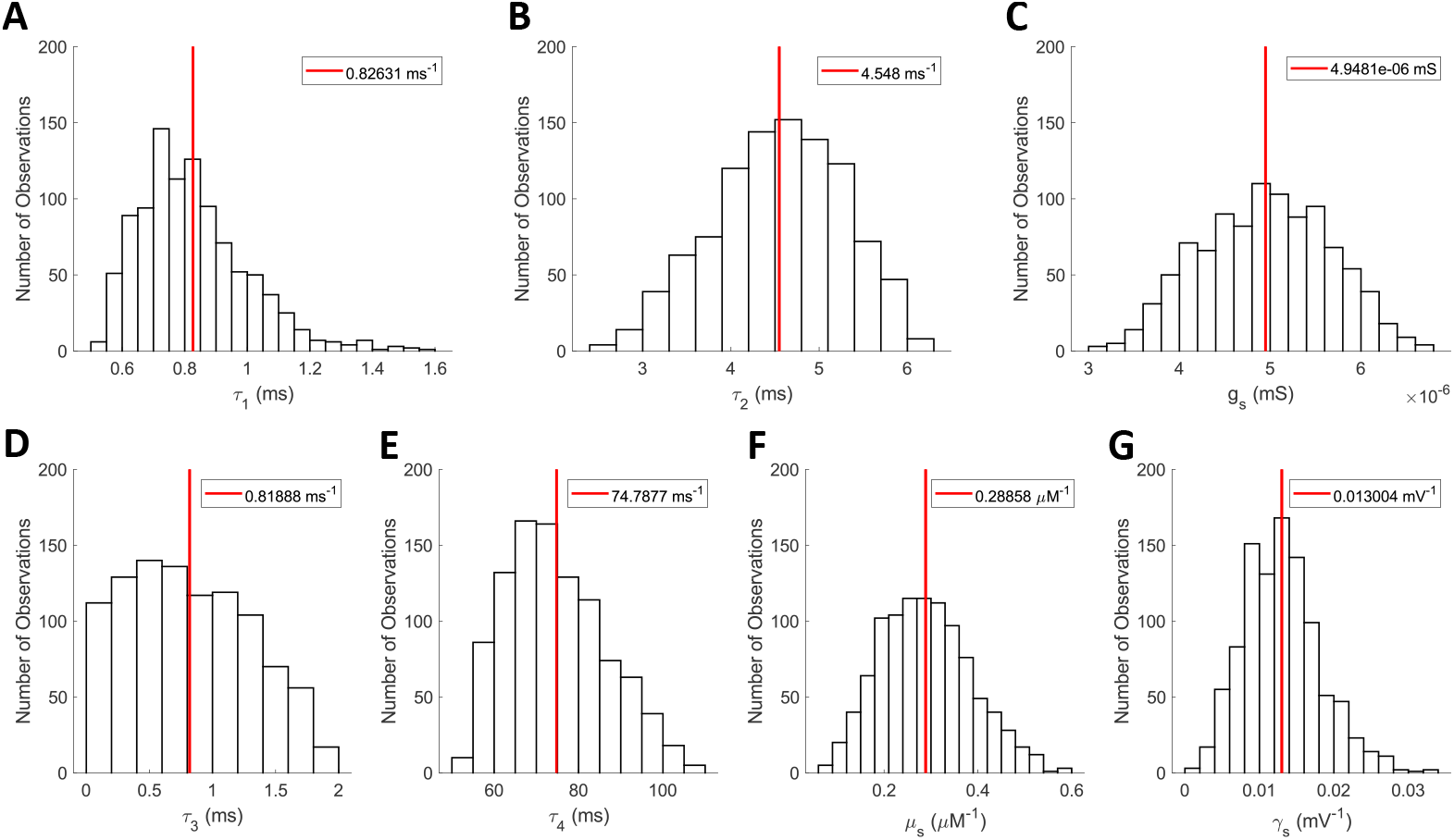
Inferred posterior distribution of the SC-CA1 synaptic dynamics parameters. Each histogram represents the approximate posterior distributions of the parameters **(A)** *τ*_1_, **(B)** *τ*_2_, **(C)** 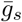, **(D)** *τ*_3_, **(E)** *τ*_4_, **(F)** *μ*_*s*_, and **(G)** *γ*_*s*_. The red-line represents the mean value.

### Frequency dependent SC-CA1 LTP/LTD dynamics

In order to model the dynamics of the high-frequency stimulation (HFS) induced long-term potentiation (LTP) and the low-frequency stimulation (LFS) induced long-term depression (LTD), we modified a published phenomenological model [31, 32] and inferred the model parameters using the available hippocampal slice experimental data in the literature on the HFS/LFS induced LTP/LTD. This simplified model assumes that the HFS/LFS induced LTP/LTD is mediated by the change in the maximum synaptic conductance 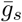 of the SC-CA1 synapse due to the changes in the calcium (Ca^2+^) current that enters through the NMDA receptors. Although the model does not include the intermediate biochemical dynamics, it exhibits multiple fixed points, which allow it to capture the frequency-dependent effects of the HFS/LFS protocol on the induced LTP/LTD. We provided the mathematical representation of the modified model in Eqs. (6a)-(6i).

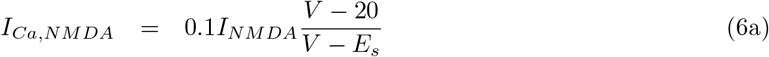

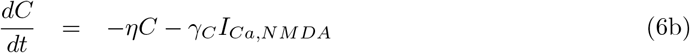

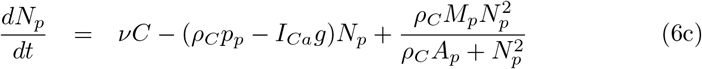

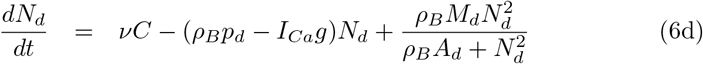

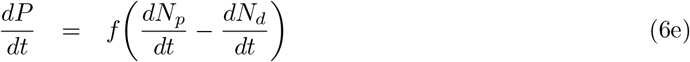

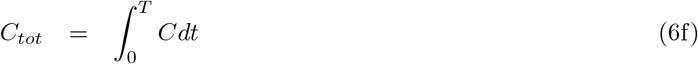

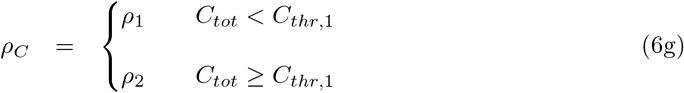

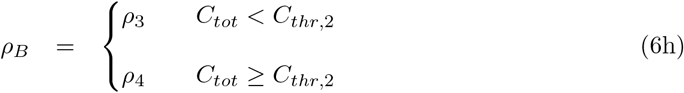

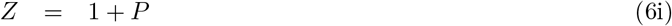

Eq. (6a), together with Eq. (1), Eqs. (3a)-(3r), and Eqs. (4a)-(4h), describe the frequency-dependent HFS/LFS induced modulation by the NMDA Ca^2+^ current, *I*_*Ca,NMDA*_. *γ*_*C*_ is the rate constant and 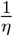 is the time constant of the dynamics. Eqs. (6c)-(6i) model the HFS/LFS induced net changes in the maximum synaptic conductance 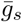 (see Eqs. (4a)-(4h)) of the SC-CA1 synapse. The model parameters *ρ*_*C*_, *ρ*_*B*_, *p*_*p*_, *p*_*d*_, *g*, *M*_*p*_, *M*_*d*_, *A*_*p*_, and *A*_*d*_ in Eqs. (6c)-(6d) control the number of fixed points exhibited by the model. The model parameter *f* in Eq. (6e) is a constant scaling factor, which determines the relative contribution of the HFS/LFS induced changes in the maximum synaptic conductance 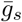. *ρ*_*C*_ toggles the equilibrium points in order to shift the weak HFS (*ρ*_1_) to the strong HFS (*ρ*_2_) protocol. Furthermore, *ρ*_*C*_ and *ρ*_*B*_ toggle LFS induced plasticity between the different LTD equilibrium points. The equilibrium points shift when the total amount of calcium, *C*_*tot*_, crosses one of the calcium thresholds *C*_*thr*,1_ or *C*_*thr*,2_. *P* is the change in the strength of the SC-CA1 synapse from the control condition. This change has been incorporated in our model by modifying the maximum synaptic conductance 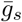 by the factor 1 + *P*.

Using Eqs. (6a)-(6i), we modified our synaptic current model shown in Eq. (4a) as

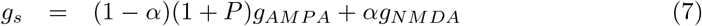

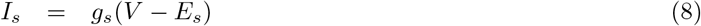

to incorporate the effect of the HFS/LFS induced changes in the synaptic current to the CA1 pyramidal neuron. With this, the complete model of the HFS/LFS induced LTP/LTD is described by Eq (1), Eqs (2a)-(2h), Eqs (3a)-(3r), Eqs (4b)-(4h), Eqs (6a)-(6i), and Eq (8).

To complete our modeling such that our model predicts the experimentally observed HFS/LFS induced LTP/LTD in hippocampal slices, we fixed the model parameters in Eq. (1), Eqs. (2a)-(2h), Eqs. (3a)-(3r), and Eq. (4b)-(4h) to the estimated model parameters values tabulated in Tables 1 and 2. We inferred the remaining model parameters in Eqs. (6a)-(6i) using the available experimental data in the literature from various hippocampal slice experiments on the % change in field excitatory postsynaptic potential (fEPSP) slope in response to various HFS/LFS protocols [2, 7, 27, 35, 37, 38, 49–57].

Our model provides the intracellular excitatory post-synaptic potential (iEPSP). Therefore, we required a transformation from iEPSP to fEPSP to compare our model output to the experimental measurements of fEPSPs from the stratum radiatum of the CA1 region. To find a transformation function, we used recent data on the simultaneous recording of fEPSP slope and iEPSP slope from the hippocampal slices [28]. Since the experimental data showed an almost linear relation between fEPSP and iEPSP, we performed a least-squares regression to obtain a linear transformation function (see also Figure 21A)

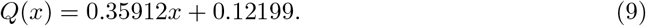

**Fig 21.**
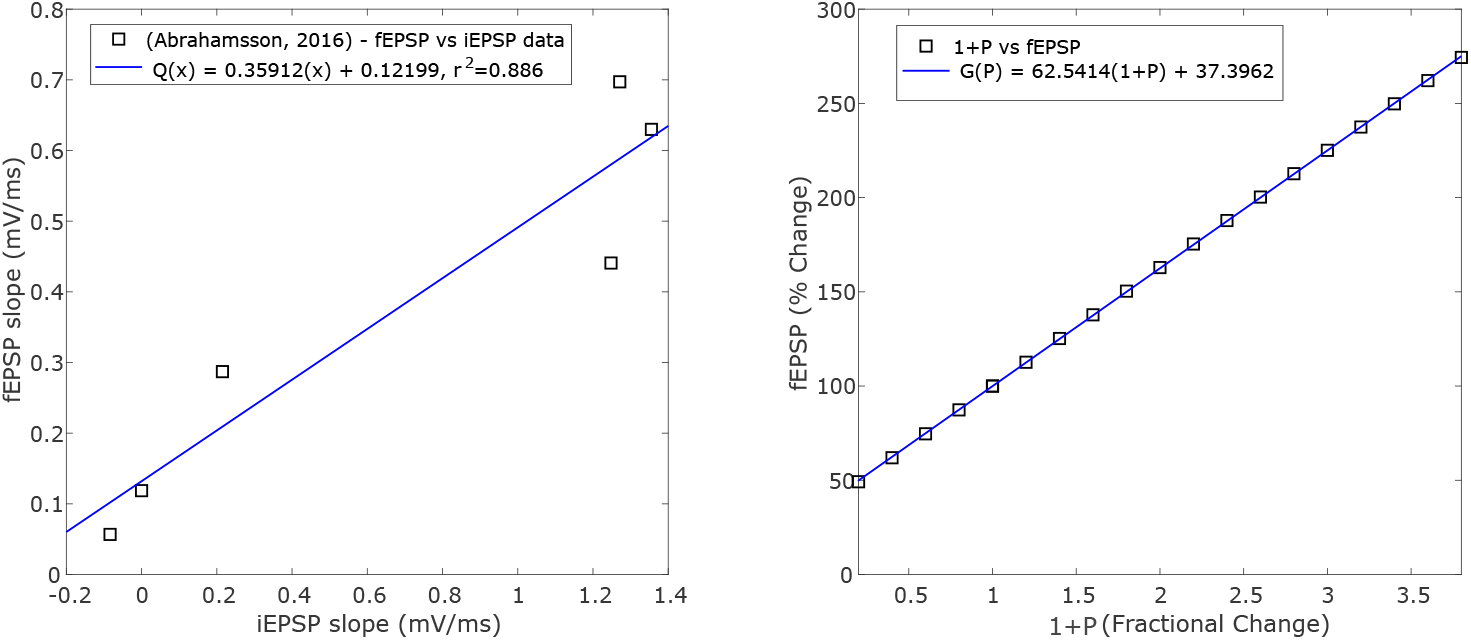
Mapping Functions. **(A)** shows a linear relationship between fEPSP and iEPSP (black-squares). The fitted linear least-squares regression *Q*(*x*) mapping fEPSP to iEPSP is shown as a blue-line. **(B)** shows a linear relationship between the fractional change in the maximum synaptic conductance (1 + *P*) and % change in the slope of the evoked fEPSPs (black-squares). The linear relationship was fit with a least-squares regression *G*(*P*) shown as a blue-line.

Here, *Q*(*x*) and *x* are the fEPSP and iEPSP slopes, respectively, at a given time.

Furthermore, to infer the model parameters more efficiently, we derived a linear mapping between the fractional change in the maximum SC-CA1 synaptic conductance 1 + *P* (see Eq. (6i)) and the % change in the evoked fEPSP measurements in our model by performing a least-squares regression (see Eq. 10 as well as Figure 21B).

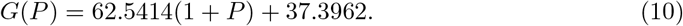

Here, *G*(*P*) is the % change in the evoked fEPSP slope at a given time with *G*(*P*) = 100% as the normalized base line value corresponding to *P* = 0.

To find a single set of parameters values for the model parameters *η*, *γ*_*C*_, *ν*, *ρ*_1_, *ρ*_2_, *ρ*_3_, *ρ*_4_, *p*_*p*_, *p*_*d*_, *g*, *M*_*p*_, *M*_*d*_, *A*_*p*_, *A*_*d*_, and *f* involved in Eqs. (6a)-(6i) that captures the experimentally observed HFS and LFS induced LTP and LTD, respectively, in this simplified model is a challenging problem. We first hand-fitted the parameters *ρ*_1_, *ρ*_2_, *ρ*_3_, and *ρ*_4_ to establish the multiple equilibrium points present in LTP and LTD, as observed in the experimental data. Then, we inferred two sets of parameters values, one for the HFS-induced LTP and another for the LFS-induced LTD, from the available experimental data in the literature on the HFS-induced LTP [2, 35, 37, 38, 49–55] and on the LFS-induced LTD [7, 27, 56–59]. The model parameters for the HFS-induced LTP and the LFS-induced LTD were inferred separately using an approximate Bayesian inference approached based on sequential Monte Carlo [33]. Below we describe the details of the parameter estimation approach from the experimental data for both LTP and LTD.

We first categorized the unknown model parameters into two sets. The first set consisted of model parameters *p*_*p*_, *p*_*d*_, *M*_*p*_, *M*_*d*_, *A*_*p*_, and *A*_*d*_, which are associated with the late-LTP or late-LTD and govern the steady-state changes in the late-LTP or late-LTD. The second set consisted of model parameters *f* , *γ*_*C*_, *η*, and *g*, which govern the fast early-LTP or early LTD. To infer the model parameters, we noticed that the slow late-LTP or late-LTD parameters are independent of the spiking activity induced by the HFS or LFS protocol in the CA1 pyramidal neuron. This led us to set the NMDA calcium current *I*_*Ca,NMDA*_ = 0 in Eq. (6a) which resulted in the decay of *C* to 0 on the order of few minutes based the fast rate of decay parameter *η*. We then applied the steady-state conditions on the Eqs. (6a)–(6d), which resulted in

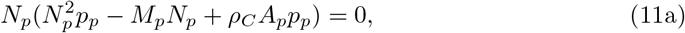

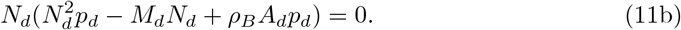

Note that Eqs. (11a)-(11b) are cubic in terms of the model parameters *N*_*p*_ and *N*_*d*_, and thus the solution of these equations can exhibit at most 6 equilibrium points (3 from Eq (11a) and 3 from Eq. (11b)). Upon examination of Eqs. (11a) and (11b), we noticed that two of the equilibrium points are the trivial stable fixed points (i.e., 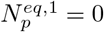 and 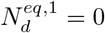). Assuming that 6 equilibrium points exist for Eqs. (11a) and (11b), we further noticed that Eq. (11a) (or Eq. (11b)) can exhibit either two stable equilibrium points and one unstable equilibrium point or one stable equilibrium point and two unstable equilibrium points. Based on these observations, we developed a cost (distance) function (see Eq. (13)) to enforce the existence of 3 equilibrium points with two stable and one unstable equilibrium points for *N*_*p*_ and *N*_*d*_ in inferring the model parameters. In our notations, 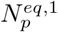 and 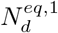 represent the stable equilibrium points of the system in the absence of a HFS/LFS protocol. 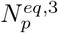 and 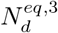 are the stable equilibrium points of the system in the presence of a HFS/LFS protocol. 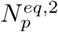 and 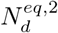 are the unstable equilibrium points.

Then, we used the equilibrium points from Eqs. (11a) and (11b) to solve for the scaling parameter 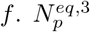 and 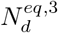 represent the late-LTP (or late-LTD) equilibrium points induced by HFS (or LFS). We used the mean LTP induced by one train of 100 *Hz* stimulation for one second in experimental data to compute the scaling parameter *f* of the LTP model [52, 54, 55]. Additionally, the scaling parameter *f* of the LTD model was computed using the experimental data on the mean LTD induced by either a stimulation protocol of 900 pulses at 1 Hz or 1200 pulses at 3 Hz [7, 27, 58]. Since the LTD protocols consisting of 900 pulses at 1 Hz or 1200 pulses at 3 Hz induced the same level of LTD in experiments, we averaged the scaling parameter *f* computed from each protocol to determine *f* in the LTD case.

Using Eq. (10) and the solution of the differential equation (6e) (*f*(*N*_*p*_ − *N*_*d*_)), we computed *f* as

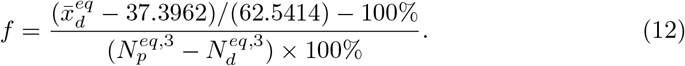

Here 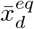 represents the average HFS/LFS induced LTP/LTD from the corresponding experimental data.

It is computationally expensive to infer the frequency dependent HFS/LFS induced LTP/LTD model parameters using the the biophysical CA1 pyramidal Hodgkin-Huxley model (Eq. (1)) and the HFS/LFS model (Eqs. (6a)-(6e)) together. Therefore, we used a reduced model to fit the remaining parameters *γ*_*C*_, *η*, *g*, *p*_*p*_, *p*_*d*_, *M*_*p*_, *M*_*d*_, *A*_*p*_, and *A*_*d*_. In the reduced model, we only considered the frequency dependent plasticity model (Eqs. (6a)-(6e)) and the characteristic NMDA calcium current for each HFS and LFS protocol when fitting the frequency dependent plasticity parameters. In order to determine the % change in the fEPSP slope, we used Eq. (10) to map the change in the conductance *P* to % change in fEPSP slope. Then, the reduced model was used to infer the frequency dependent model parameters using an approximate Bayesian inference approach based on the sequential Monte Carlo (ABC-SMC) [33] with a modified mean sum of squared errors distance function (Eq. (13)) averaged over the *m* experimental data sets. The inferred parameters are provided in Table 4. Figures 22 and 23 show the histograms representing the approximate posterior distributions for the HFS and LFS parameters, respectively.

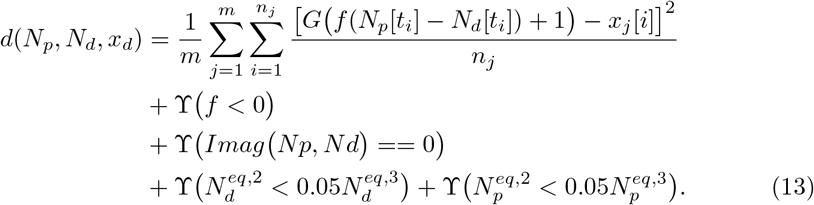

**Table 4.** Frequency Dependent Plasticity Parameters.

| Parameters | LTP Parameter Value | LTD Parameter Value |
| --- | --- | --- |
| $M_p$ | $4.83 \times 10^{-8} \pm 4.92 \times 10^{-9} \mu A ms^{-1}$ | $4.95 \times 10^{-7} \pm 5.06 \times 10^{-8} \mu A ms^{-1}$ |
| $p_p$ | $1.01 \times 10^{-5} \pm 1.03 \times 10^{-6} ms^{-1}$ | $2.16 \times 10^{-4} \pm 2.41 \times 10^{-5} ms^{-1}$ |
| $A_p$ | $2.30 \times 10^{-6} \pm 2.54 \times 10^{-7} \mu A^2$ | $9.76 \times 10^{-7} \pm 1.11 \times 10^{-7} \mu A^2$ |
| $M_d$ | $4.91 \times 10^{-7} \pm 5.16 \times 10^{-8} \mu A ms^{-1}$ | $5.81 \times 10^{-8} \pm 5.41 \times 10^{-9} \mu A ms^{-1}$ |
| $p_d$ | $2.23 \times 10^{-4} \pm 2.51 \times 10^{-5} ms^{-1}$ | $1.63 \times 10^{-5} \pm 1.81 \times 10^{-6} ms^{-1}$ |
| $A_d$ | $6.81 \times 10^{-7} \pm 7.88 \times 10^{-8} \mu A^2$ | $2.07 \times 10^{-6} \pm 2.40 \times 10^{-7} \mu A^2$ |
| $\gamma_C$ | $0.0364 \pm 0.0036 ms^{-1}$ | $0.0238 \pm 0.0027 ms^{-1}$ |
| $\eta$ | $0.0035 \pm 0.000382 ms^{-1}$ | $0.0024 \pm 0.000276 ms^{-1}$ |
| $\nu$ | $0.107 \pm 0.0122 ms^{-1}$ | $0.0348 \pm 0.0038 ms^{-1}$ |
| $g$ | $214.1 \pm 24.6 \mu A^{-1} ms^{-1}$ | $52.0 \pm 6.13 \mu A^{-1} ms^{-1}$ |
| $f$ | $460 \mu A^{-1}$ | $281.3 \mu A^{-1}$ |
| $\rho_1$ | 2.2 | 1 |
| $\rho_2$ | 1 | 1.31 |
| $\rho_3$ | 1 | 1 |
| $\rho_4$ | 1 | 0.35 |
| $C_{thr,1}$ | 0.45 | 3 |
| $C_{thr,2}$ | 0 | 4.5 |

**Fig 22.**
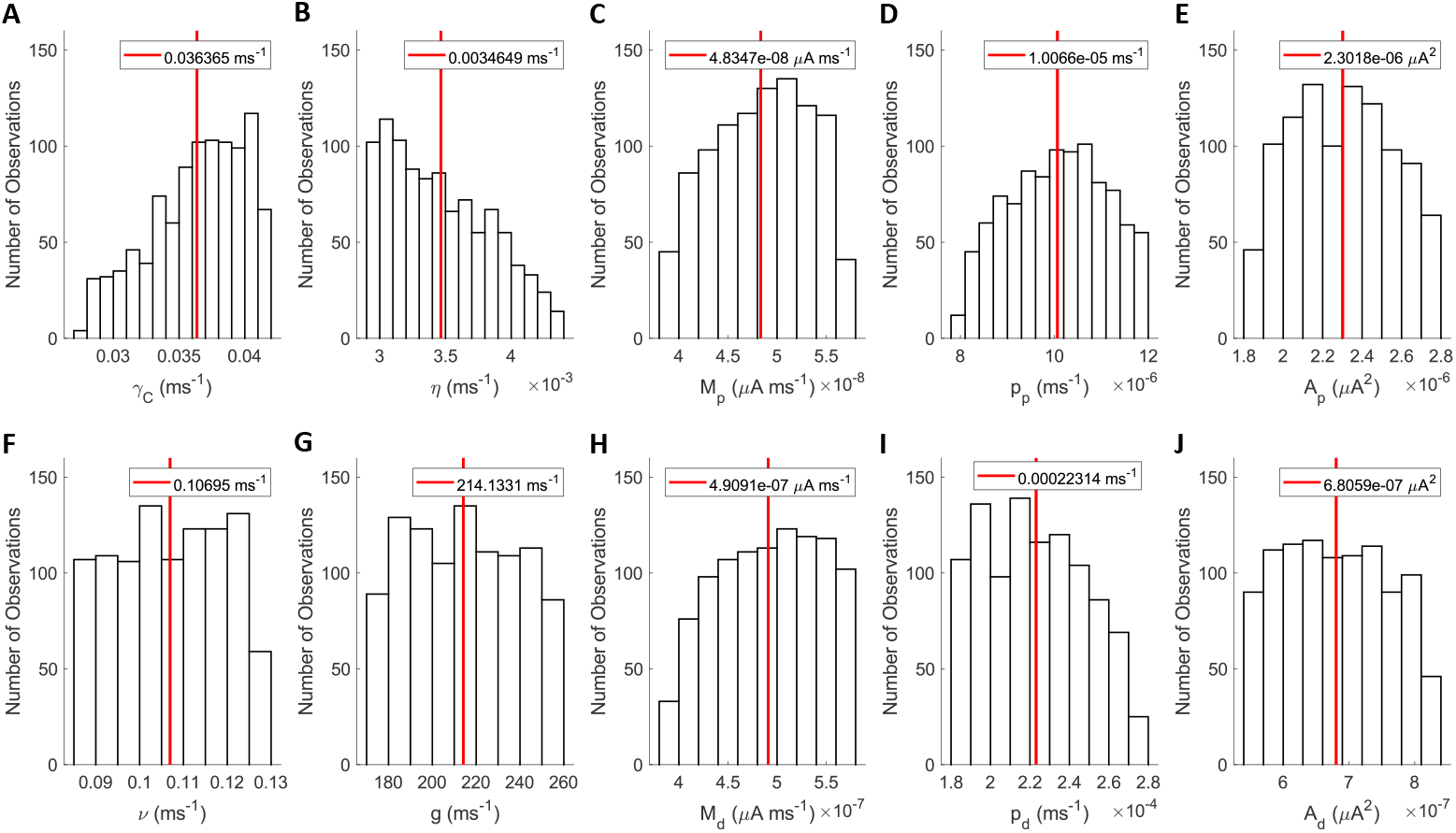
Inferred posterior distribution of the HFS-induced LTP parameters. Each histogram represents the approximate posterior distributions of the parameters **(A)** *γ*, **(B)** *η*, **(C)** *M*_*p*_, **(D)** *P*_*p*_, **(E)** *A*_*p*_, **(F)** *ν*, **(G)** *g*, **(H)** *M*_*d*_, **(I)** *P*_*d*_, and **(J)** *A*_*d*_. The red-line represents the mean value.

**Fig 23.**
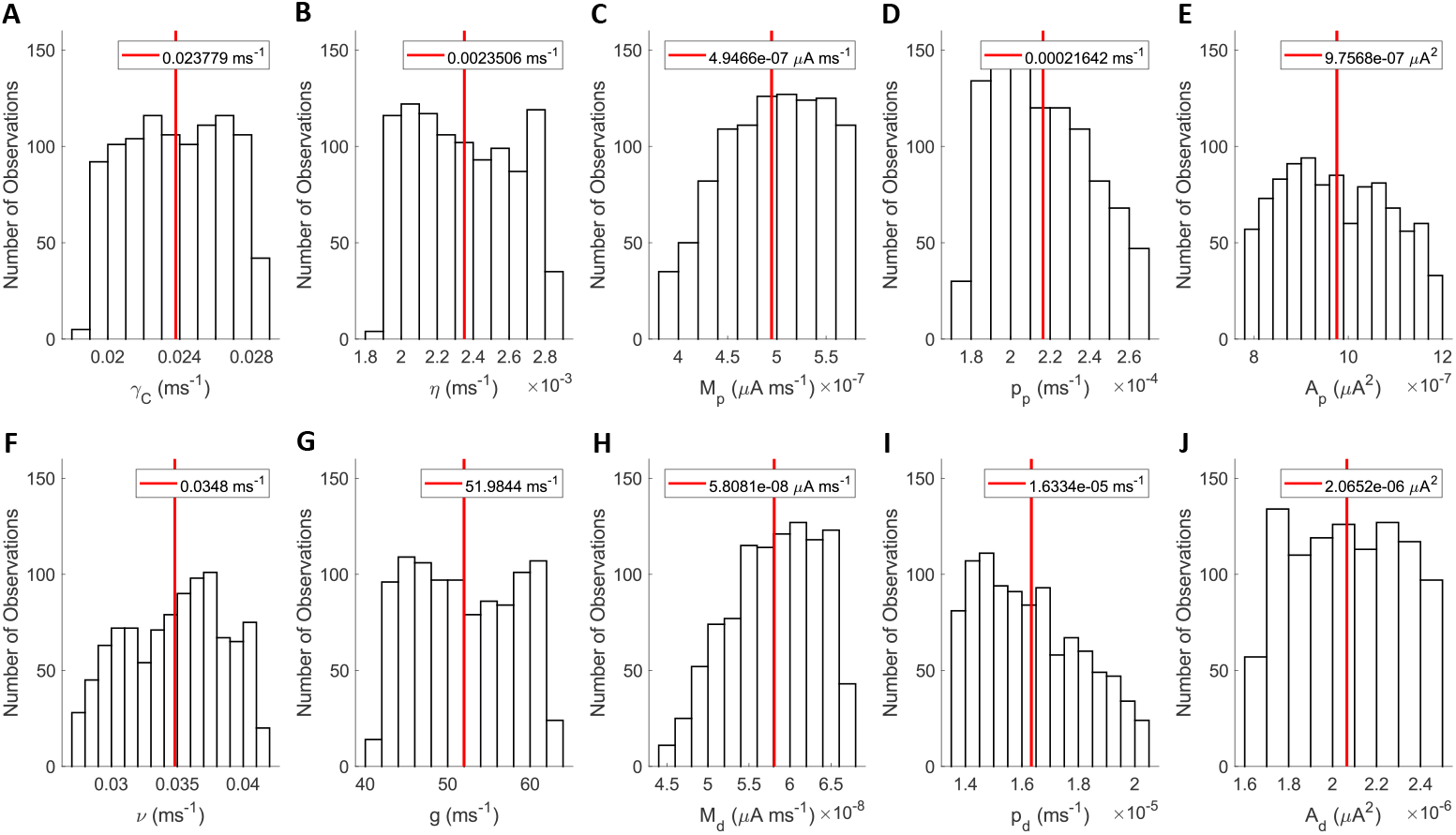
Inferred posterior distribution of the LFS-induced LTD parameters. Each histogram represents the approximate posterior distributions of the parameters *γ*, **(B)** *η*, **(C)** *M*_*p*_, **(D)** *P*_*p*_, **(E)** *A*_*p*_, **(F)** *ν*, **(G)** *g*, **(H)** *M*_*d*_, **(I)** *P*_*d*_, and **(J)** *A*_*d*_. The red-line represents the mean value.

The first term on the right hand side of the distance function *d*(*N*_*p*_, *N*_*d*_, *x*_*d*_) (see Eq. (13)) captures the error between the % change in the slope of evoked fEPSPs from our model and the corresponding % change in the fEPSP slope data from *m* different LTP/LTD experimental datasets available from the literature. The % change in the slope of evoked fEPSPs was normalized (i.e. 100%) to the slope measured prior to any HFS/LFS protocol. Each of the experimental data sets had *n*_*j*_ number of data points measuring the % change in the slope of evoked fEPSPs, *x*_*i,j*_, after the HFS/LFS protocol. We used a mapping function *G*(*P*) (see Eq. (10)) to compute the % change in the evoked fEPSP slope in our model, where 1 + *P* represents the corresponding fractional change in the maximum synaptic conductance of the SC-CA1 synapse from the baseline (i.e., *P* = 0). The fractional synaptic conductance change from the baseline in our model was determine by *P*(*t*_*i*_) = *f*(*N*_*p*_(*t*_*i*_) − *N*_*d*_(*t*_*i*_)). The baseline for the % change in fEPSP is 100%, which corresponds to *P* = 0 (see Eq. (6i)) at the time *t*_*i*_ of the experimental data point *x*_*i,j*_.

The second term on the right hand side of the distance function *d*(*N*_*p*_, *N*_*d*_, *x*_*d*_) (see Eq. (13)) ensures that only the parameters (*p*_*p*_, *p*_*d*_, *M*_*p*_, *M*_*d*_, *A*_*p*_, and *A*_*d*_) that produce a positive scaling parameter *f* are accepted. The need for this term arises out of the differential equations describing *N*_*p*_ and *N*_*d*_ in Eqs. (6c) and (6d), which are identical except for the parameters. Therefore, it is possible that the ABC-SMC algorithm could select parameters for *p*_*p*_, *M*_*p*_, and *A*_*p*_ that are typically selected for *p*_*d*_, *M*_*d*_, and *A*_*d*_ and *vice versa*. This would result in 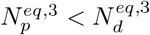 for the LTP model and 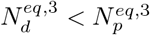 for the LTD model, which would require *f* to be negative to match the experimental data. Thus, *f* must be constrained to a strictly positive number.

The last three terms on the right hand side of the distance function *d*(*N*_*p*_, *N*_*d*_, *x*_*d*_) (see Eq. (13)) penalize the equilibrium points that are complex or have unstable equilibrium points 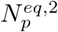 and 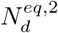 that are less than 5% of 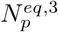 and 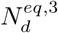, which are the stable equilibrium points of the system in the presence of a HFS/LFS protocol. The first term ϒ(*Imag*(*N*_*p*_, *Nd*) == 0) ensures that the fractional change in the maximum synaptic conductance is always a real number. The last term 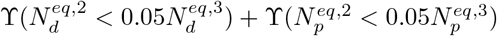 ensures that that the small evoked fEPSPs used to measure the plasticity of the SC-CA1 synapse prior to any HFS/LFS administration does not induce any unwanted LTP/LTD, such that *N*_*p*_ and *N*_*d*_ decay to 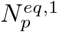 and 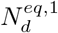 after each fEPSP measurement prior to HFS/LFS administrations. ϒ is the weighting factor.

We first used the inferred model parameters of the LTP model to validate whether our model captured the frequency-dependent effects of HFS protocol on the induced LTP, as observed in the experiments. Figures 24A, 24B, 24C, and 24D show the comparison between the prediction from our model and the experimental data for four different HFS protocols, i.e., 3 trains of pulses as 100 *Hz* for 1 second with intertrain intervals of 0.5 seconds, 10 minutes, 20 seconds, and 10 seconds, respectively. Additionally, the LTP induced by a HFS protocol of four trains of 100 pulses at 100 *Hz* is shown in Figure 25D. Here, the HFS-induced % fEPSP slope change from the control, shown as a black square-line, follows the same trend as the experimental data and achieves the same LTP magnitude change for all five protocols [35, 38, 49–52]. In order to capture the LTP induced by a weaker HFS protocol, the equilibrium points were shifted by *ρ*_*C*_ to capture the lower LTP level induced by a weaker HFS protocol. *ρ*_1_ was hand-fitted to match the LTP induced by one and two trains of pulses at 100 *Hz* [52]. Additionally, *C*_*thr*_ was chosen such that three or more trains of HFS at 100 *Hz* switched the parameter *ρ*_*C*_ to *ρ*_2_. Figure 25A shows our model is able to capture the LTP dynamics induced by 1 train of 100 *Hz* pulses reasonably well [52, 54, 55]. However, our model underpredicts the experimental HFS-induced magnitude change evoked from 2 trains of 100 *Hz* pulses [52, 53] (see Figure 25B). Even though our model underpredicts the magnitude change from 2 trains, experimental data shows that the magnitude change induced by 2 trains (see Figure 25C) is not significantly different from the magnitude change induced by 1 train [52].

**Fig 24.**
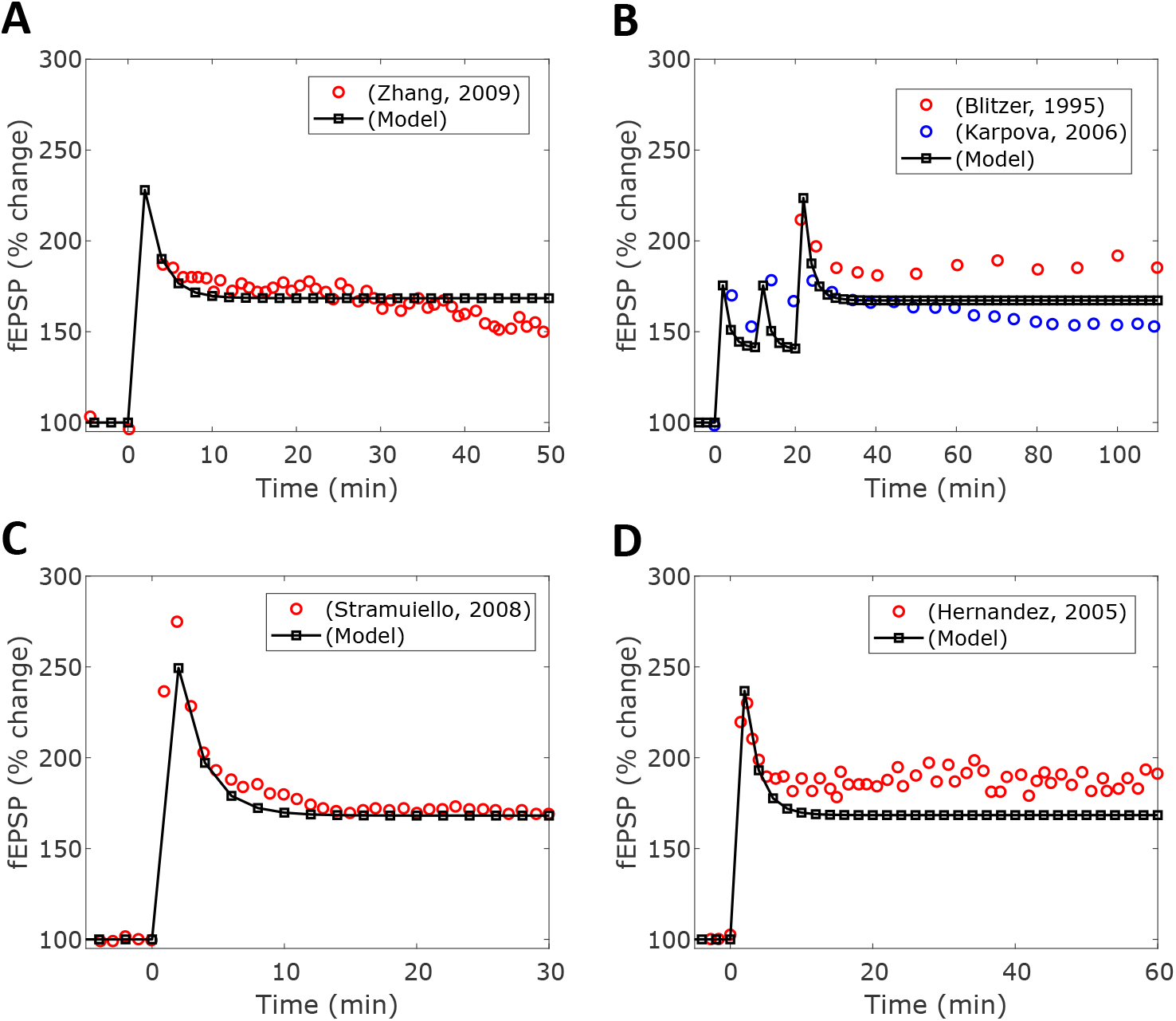
HFS-induced LTP. We provided three trains of 100 pulses delivered at 100 *Hz* to our SC-CA1 model with different inter-train intervals. The inter-train intervals were **(A)** 0.5 seconds, **(B)** 10 minutes, **(C)** 20 seconds, and **(D)** 10 seconds. The HFS-induced LTP in experiment is represented by the colored-circles and the LTP predicted by our model is shown as the black-squares.

**Fig 25.**
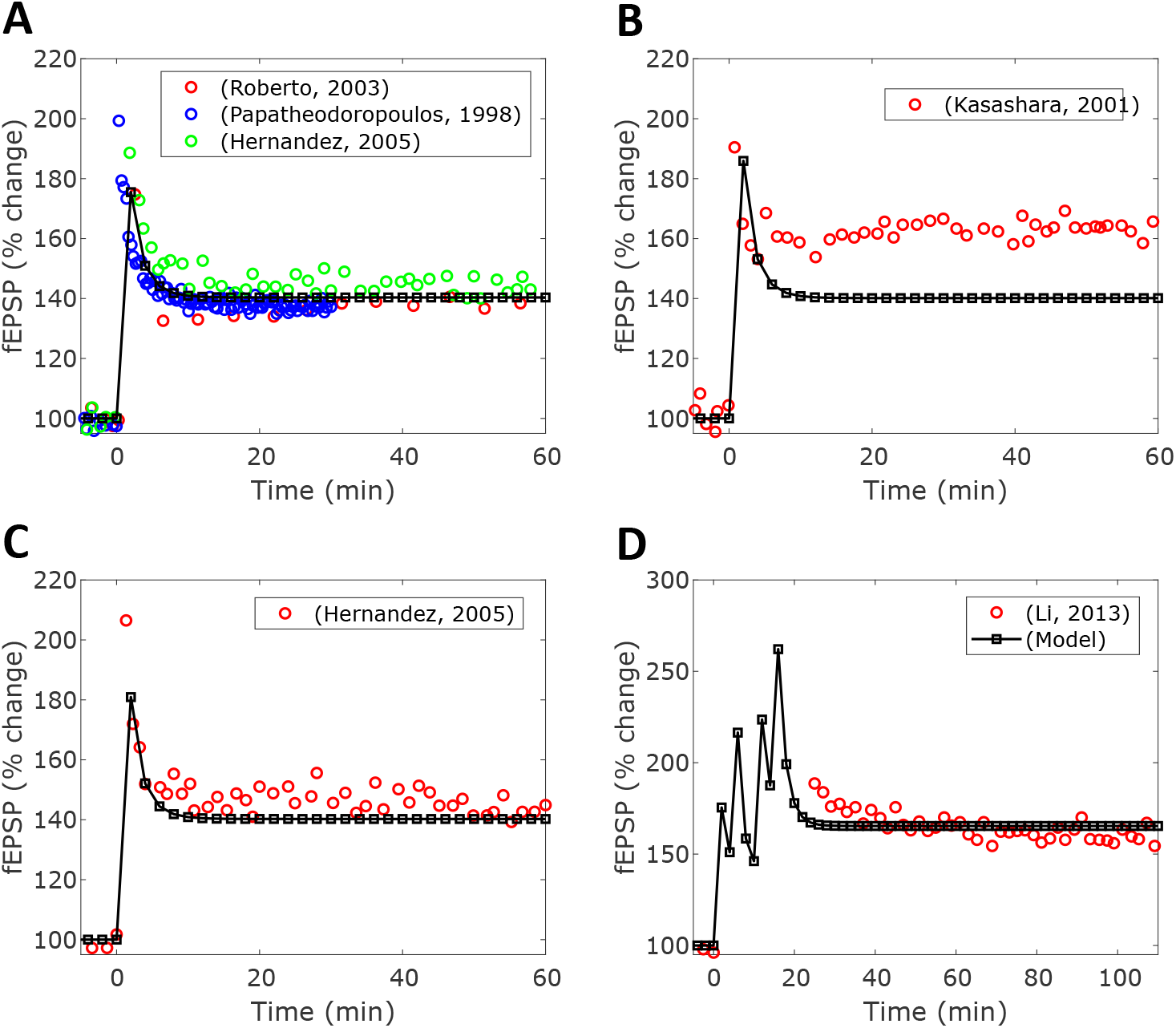
HFS-induced LTP with different HFS protocols. We provided one, two, and four trains of 100 pulses delivered at 100 *Hz* to our SC-CA1 model with different inter-train intervals. In **(A)**, we applied one trains of HFS. **(B)** and **(C)** shows the induced LTP from two trains of HFS with an inter-train interval of 10 seconds and 20 seconds, respectively. **(D)** shows the HFS-induced LTP from four trains of HFS with an inter-train of 5 minutes. In each case, the HFS-induced LTP observed in experiment is represented by the colored-circles and the LTP induced in our model is shown as the black-squares.

We then used the inferred model parameters of the LTD model to validate whether our model captured the frequency-dependent effects of LFS protocol on the induced LTD, as observed in the experiments. To validate our model, we considered seven different LFS protocols. The first LFS protocol consisted of 900 pulses delivered to the SC-CA1 synapse at the frequency of 1 Hz. Figure 26A shows the comparison between the prediction from our model and the experimental data from different experiments. As shown in this figure, our model predicted an approximately 19% decrease in the evoked fEPSP slope compared to an average of 18% decrease in experimental data (experimental data varied between 10% and 25%) [7, 56, 58, 59]. The second LFS protocol we used was 1200 pulses at 3 *Hz*, which produced a decrease in the normalized slope of the evoked fEPSPs of approximately 19% in our model compared to 21% in the experimental data [27] (see Figure 26B). Then, we considered the administration of 900 pulses at 3 *Hz*. Experimental data showed a decrement of 15% in the evoked fEPSP slope [57] while our model predicted 19% of decrement (see Figure 26C).

**Fig 26.**
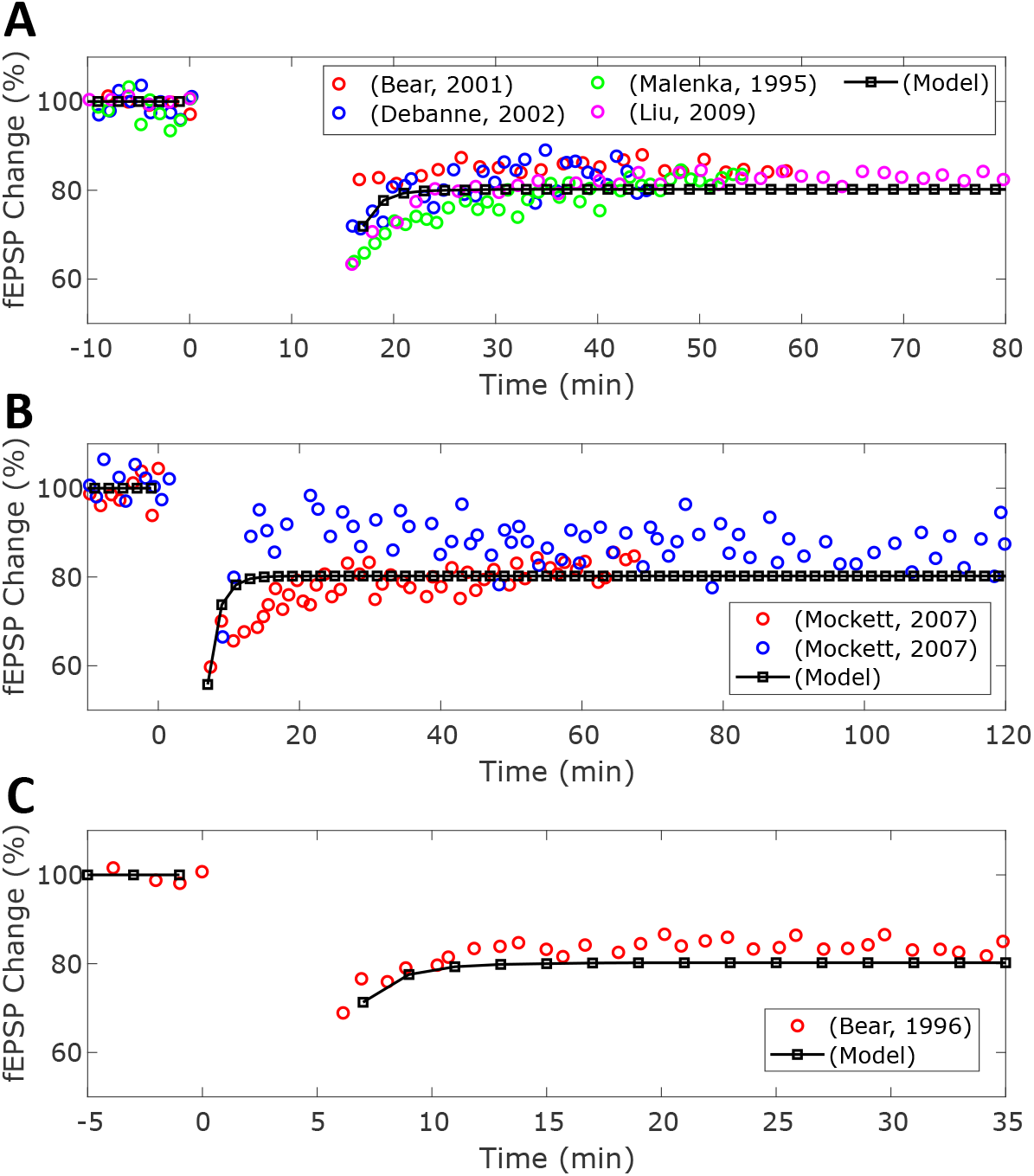
LFS-induced LTD. We provided three different LFS trains to our SC-CA1 model and compared the induced LTP in our model (black-squares) to the induced LTP in experiment (colored-circles). We considered LFS protocols consisting of **(A)** 900 pulses at 1 *Hz*, **(B)** 1200 pulses at 3 *Hz* and **(C)** 900 pulses at 3 *Hz*.

Next, we considered multiple LFS trains. Again, we captured the multiple equilibrium points of LTD induced by different LFS protocols with the parameters *ρ*_2_ and *ρ*_4_, which we fitted by hand to match the experimental data [7, 58]. Here, we used the same LTD parameters in Table 4 that were used to model one train of LFS. Then we hand-fitted the toggle parameters *ρ*_2_ and *ρ*_4_, in order to capture the multiple equilibrium points of the LFS-induced LTD. The LTD toggle parameters are found in Table 4, as well. Figure 27A shows a comparison between our model’s prediction and the experimental data from [7, 58] on the LFS-induced LTD for a LFS protocol consisting of 3 trains of 900 pulses at 1 *Hz*. As shown here, the model accurately predicts the experimentally observed decrease in the synaptic strength of the SC-CA1 synapse after each train of LFS. Figure 27B shows a comparison between the model predicted LTD in the SC-CA1 synapse with the experimental data from [8] for a LFS protocol consisting of 900 bursts at 1 *Hz* of 3 pulses at 20 *Hz*. Figure 27C compares the model predicted LTD in the SC-CA1 synapse with the experimental data from [27] for a LFS protocol consisting of 2400 pulses at 3 *Hz*. Finally, Figure 27D compares the model predicted LTD in the SC-CA1 synapse with the experimental data from [27] for a LFS protocol consisting of 2 trains of 1200 pulses at 3 *Hz*.

**Fig 27.**
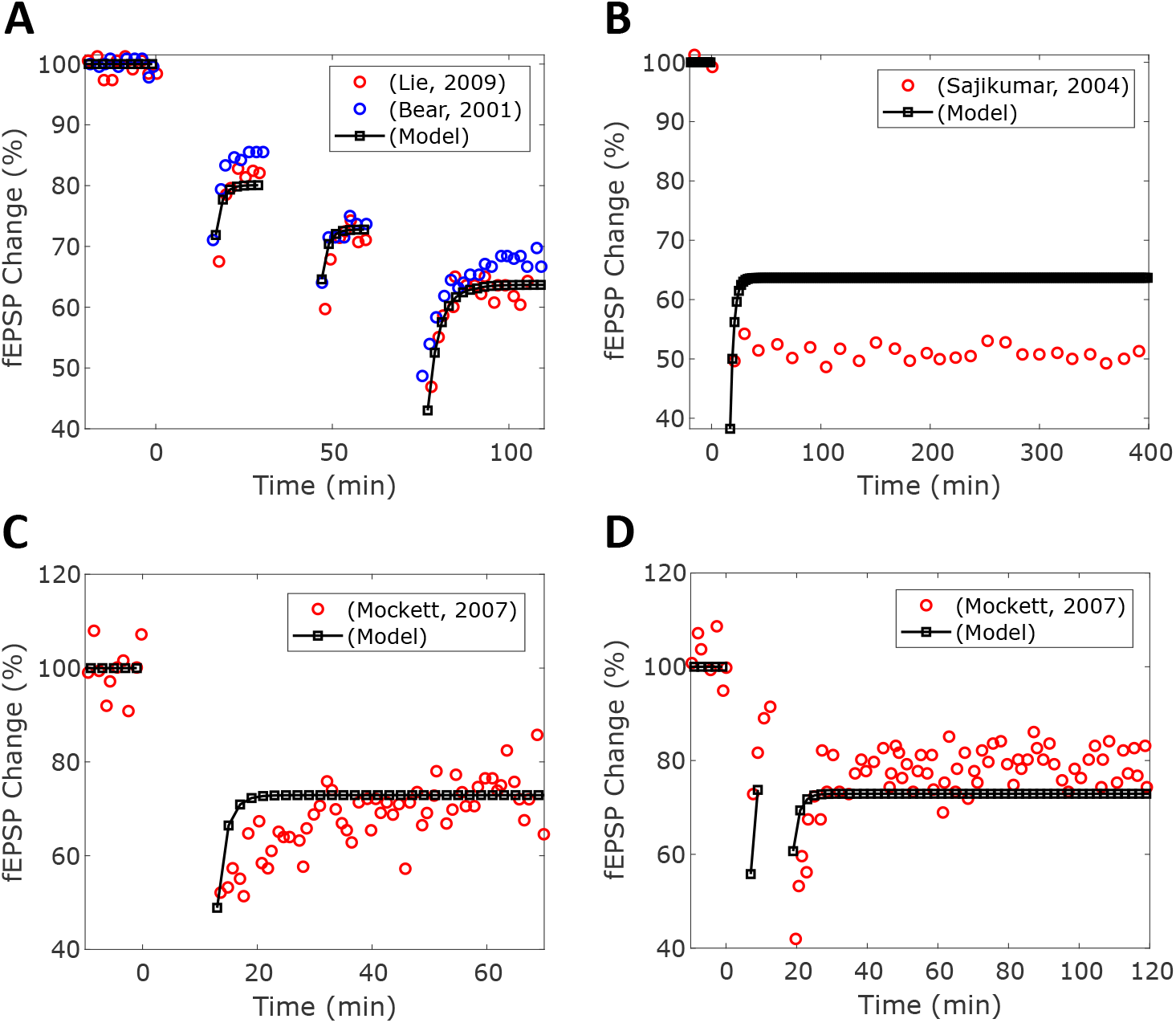
LFS-induced LTD with other LFS protocols. We provided four other LFS protocols to our SC-CA1 model and compared the induced LTP in our model (black-squares) to the induced LTP in experiment (colored-circles). We considered LFS protocols consisting of **(A)** three trains of 900 pulses at 1 *Hz* with a inter-train interval of 15 minutes, **(B)** 900 burst of 3 pulses at 1 *Hz* where the pulses in the burst were delivered at 20 *Hz*, **(C)** 2400 pulses at 3 *Hz*, and **(D)** two trains of 1200 pulses at 3 *Hz*.

### Dopaminergic modulation of HFS/LFS induced LTP/LTD in hippocampal SC-CA1 synapses

In this section, we describe a phenomenological model that we developed to integrate the dose-dependent effects of D1/D5 agonists relative to the HFS/LFS on the maximum synaptic conductance (see Eq. (8)) of the SC-CA1 synapse. Our model together with the models described in the previous sections (see Eqs (1)-(8)) is capable of predicting the experimentally observed dose-dependent modulation of HFS/LFS-induced LTP/LTD by D1/D5 agonists. Eqs. (14a)-(14d) show our model in the form of a set of kinetic reactions.

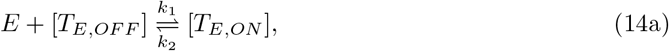

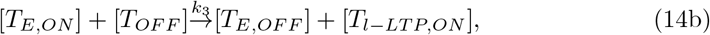

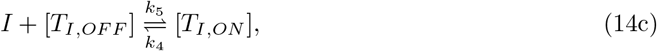

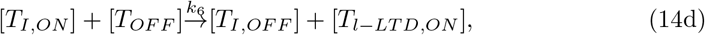

These reactions describe the modulation of the maximum synaptic conductance 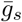 of the SC-CA1 synapse by a D1/D5 agonist, such as SKF 38393, 6-Br-APB, or dopamine, in the absence of a LTP/LTD protocol (i.e., HFS/LFS). Investigations on the biochemical mechanisms underlying the modulation of SC-CA1 long-term synaptic plasticity by D1/D5 agonists have revealed two competing pathways, the excitatory adenyl cyclase (AC) pathway and the inhibitory phosopholipase C (PLC) pathway [44, 60, 61]. In our model, we captured the dynamics exhibited by the excitatory AC pathway, that induces slow-onset-potentiation, using Eqs. (14a) and (14b). The response of the inhibitory PLC pathway, that induces a slow depotentiation in the SC-CA1 synapses, is captured by Eqs. (14c) and (14d). The change in the maximum synaptic conductance is dependent on the activation of either the AC or PLC pathway through the tags *T*_*E,ON*_ and *T*_*I,ON*_ , respectively. These pathways compete for *T*_*OFF*_ to produce l-LTP (*T*_*l*−*LTP,ON*_) or l-LTD (*T*_*l*−*LTP,ON*_) changes to the SC-CA1 synaptic strength through the maximum synaptic conductance.

Mathematically, we modeled these reactions in a deterministic kinetic rate modeling framework using Eqs. (15c)-(15h) and Eq (15m) with *θ*_*late*_ = 0 and *T*_*stim*_ = 0 in Eq. (15m). Eqs. (15a)-(15b) are the Hill equations that model the dose response of a D1/D5 agonist at a given concentration [*Drug*]. The activation of the AC and PLC pathways is bound by a maximal response (*E*_*max*_ and *I*_*max*_). The fraction of maximal response is determined by the Hill parameters *EC*50, *E*_*max*_, *IC*50, *I*_*max*_, and *nH* for each D1/D5 agonist. We estimated the Hill coefficients using the available experimental data on the % change in the CA1 fEPSP slope in response to the applied concentration of the D1/D5 agonist [4, 8, 34].

The specific D1/D5 agonist and its temporal dosing profile determine the relative activation of the AC and PLC pathways. The activation dynamics of the AC (*T*_*E,ON*_) and PLC (*T*_*I,ON*_) pathway are described by Eqs. (15c) - (15f). Eqs. (15g) - (15h) show the consumption dynamics of *T*_*OFF*_ to produce 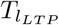 and 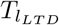.

Eqs. (15m) - (15r) model the integrative effect of HFS/LFS mediated LTP/LTD and D1/D5 agonist on the SC-CA1 LTP/LTD. The combined effect of HFS/LFS and D1/D5 agonist on the maximum synaptic conductance of the SC-CA1 synapse is described in Eq. (15o). Here, the term 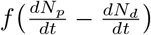 captures the contribution from the HFS/LFS protocol and the term 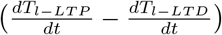 captures the contribution from a D1/D5 agonist. Experimental data from hippocampal slices show that the LTP-induced by a combination of strong HFS protocol and a D1/D5 agonist in SC-CA1 synapses slowly decays back to the HFS only induced late LTP in a period of 2-4 hours [2, 34]. We hypothesized that this may be due to the maximum available biochemical resources for inducing late LTP and captured this effect in our model by introducing a decay term *k*_*sat*_(*P − P*_*sat*_). Particularly, whenever the induced LTP exceeds the saturation level, defined by *P*_*sat*_ in the presence of a D1/D5 receptors agonist at time *t*_*DA*_, *k*_*sat*_ takes a nonzero value. The term *k*_*basal*_*P* models the effect of D1/D5 antagonist, such as SCH 23390. The parameter *k*_*basal*_ takes a nonzero value *k*_*sat*_ whenever SCH 23390 is applied to block the D1/D5 receptors. Finally, the term *sign* (*E* + *k*_*I*_*I*)*k*_*E*_*T*_*stim*_*T*_*DR*_ models the nonlinear interaction of the D1/D5 receptors activation and HFS/LFS on the induced LTP/LTD.

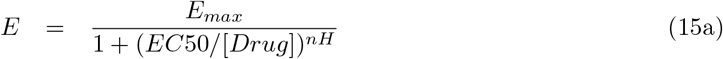

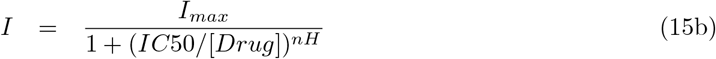

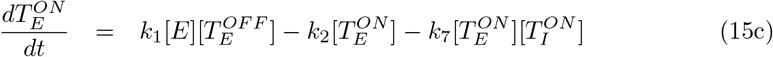

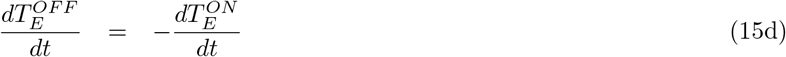

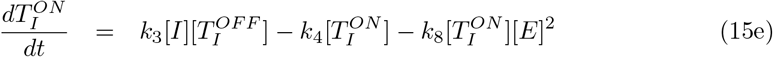

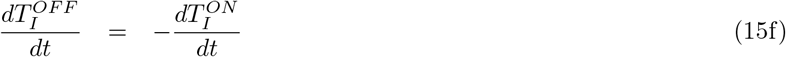

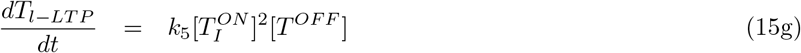

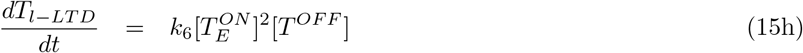

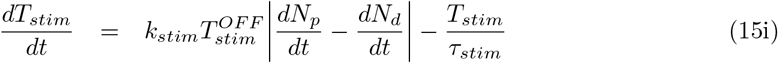

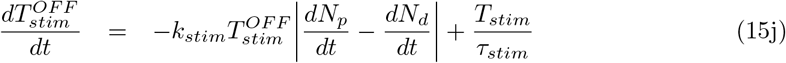

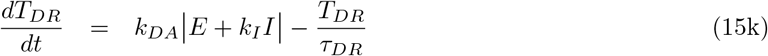

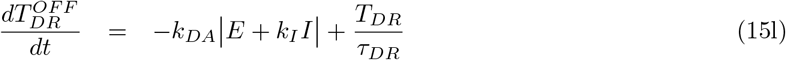

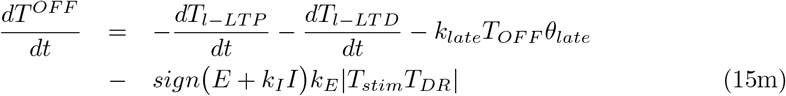

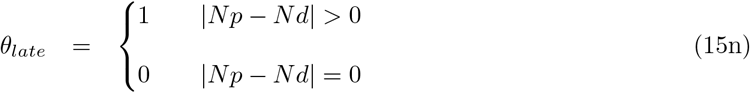

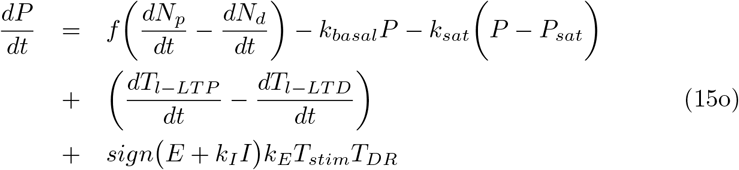

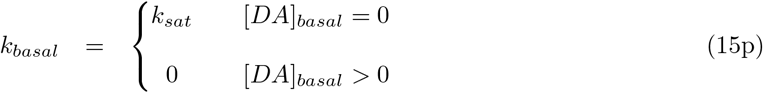

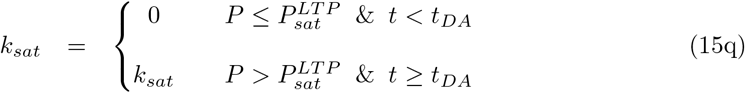

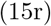

To infer the unknown model parameters *k*_1_, *k*_2_, *k*_3_, *k*_4_, *k*_5_, *k*_6_, *k*_7_, *k*_8_, *k*_*sat*_, *k*_*late*_, *k*_*E*_, *k*_*I*_ , *k*_*stim*_, *k*_*DA*_, *τ*_*stim*_, and *τ*_*DA*_ from the available experimental data, we observed that the set of Eqs. (15a)- (15h) are independent of the set of Eqs. (15i)- (15r). This allowed us to infer the unknown model parameters efficiently using the ABC-SMC approach.

Specifically, we first inferred the model parameters of the first set of equations (Eqs. (15a)- (15h)), and which was then followed by the inference of model parameters of the second set of equations (Eqs (15i)- (15r)).

We first inferred the 8 unknown model parameters *k*_1_, *k*_2_, *k*_3_, *k*_4_, *k*_5_, *k*_6_, *k*_7_, and *k*_8_ in Eqs. (15a)-(15h) using the ABC-SMC method [33]. For the parameter inference, we used the input-output data (% change in fEPSP slope from control in response to various concentrations of the D1/D5 agonists SKF 38393, 6-bromo-APB, and dopamine) available from recent rat hippocampal slice experiments [4, 8, 34]. It has been shown in [4] that the application of a D1/D5 agonist alone, in the absence of a HFS-based LTP protocol, can potentiate the SC-CA1 synapses in a dose-dependent manner. Moreover, these dopaminergic mediated potentiations occur on a slow timescale (typically, several minutes to hours). Since HFS or LFS was not applied when measuring the slow-onset dopaminergic potentiation, we only considered the dopaminergic model (Eqs. (15a)-(15h)) when inferring the parameters *k*_1_, *k*_2_, *k*_3_, *k*_4_, *k*_5_, *k*_6_, *k*_7_, and *k*_8_. We considered a modified mean sum of squared errors distance function (Eq. (16b)) averaged over the *m* experimental data sets that compares the distance between the *i*^*th*^ experimental % change in fEPSP slope (*x*_*j*_[*i*]) of the *j*^*th*^ data set to the corresponding % change in fEPSP slope in the model *x*_*m*_[*t*_*i*_] at time *t*_*i*_. In addition to the mean sum of squared error, the distance function contains an additional non-steady state penalty in the second term on the right hand side of Eq. (13) . Table 6 shows our inferred parameters value represented in terms of the mean ± standard deviation and Figure 28 shows the histograms representing the approximate posterior distribution for each parameter.

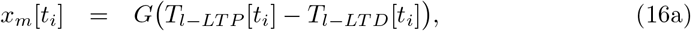

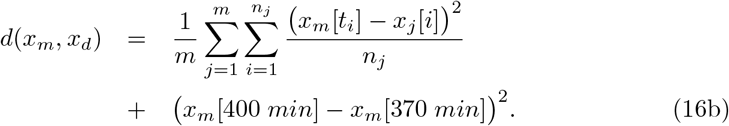

**Table 5.**
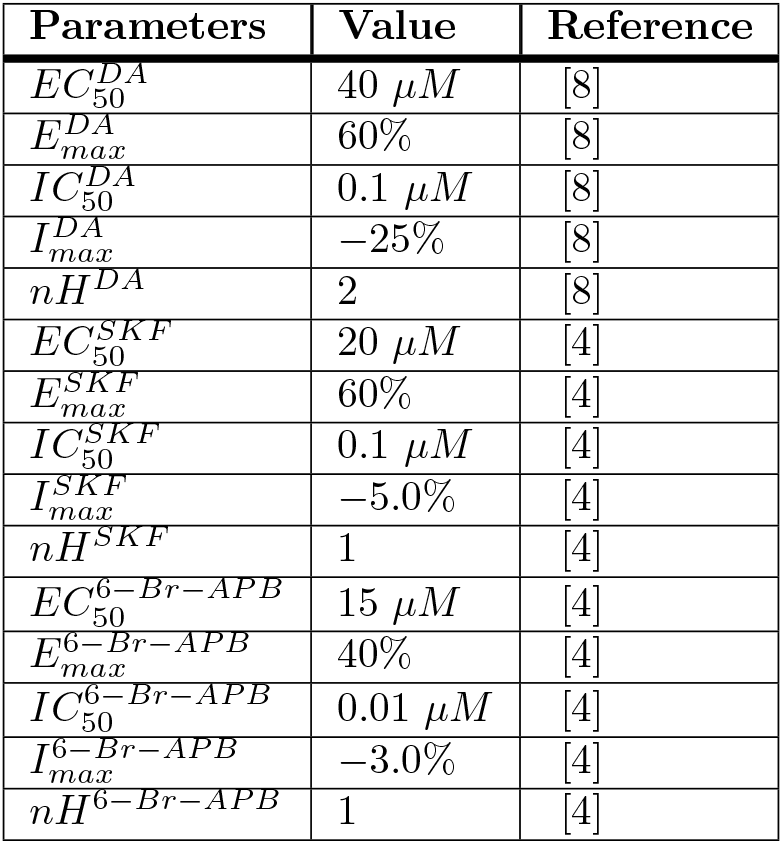
Hill Function Parameters.

**Table 6.** Dopaminergic Potentiation Parameters.

| Parameters | Value |
| --- | --- |
| $k_1$ | $8.26 \times 10^{-6} \pm 2.17 \times 10^{-6} \text{ ms}^{-1}$ |
| $k_2$ | $7.10 \times 10^{-8} \pm 2.99 \times 10^{-8} \text{ ms}^{-1}$ |
| $k_3$ | $3.24 \times 10^{-6} \pm 4.25 \times 10^{-6} \text{ ms}^{-1}$ |
| $k_4$ | $1.44 \times 10^{-7} \pm 5.27 \times 10^{-7} \text{ ms}^{-1}$ |
| $k_5$ | $2.89 \times 10^{-7} \pm 7.45 \times 10^{-8} \text{ ms}^{-1}$ |
| $k_6$ | $2.35 \times 10^{-6} \pm 4.31 \times 10^{-6} \text{ ms}^{-1}$ |
| $k_7$ | $1.63 \times 10^{-7} \pm 8.69 \times 10^{-7} \text{ ms}^{-1}$ |
| $k_8$ | $5.32 \times 10^{-4} \pm 6.90 \times 10^{-4} \text{ ms}^{-1}$ |
| $k_{sat}$ | $1.34 \times 10^{-7} \pm 1.07 \times 10^{-8} \text{ ms}^{-1}$ |
| $k_{stim}$ | $557 \pm 128 \text{ ms}^{-1}$ |
| $k_E$ | $1.51 \times 10^{-6} \pm 2.20 \times 10^{-7} \text{ ms}^{-1}$ |
| $k_I$ | $2.49 \pm 0.306$ |
| $k_{late}$ | $4.36 \times 10^{-7} \pm 4.49 \times 10^{-8} \text{ ms}^{-1}$ |
| $k_{DA}$ | $1.0 \times 10^{-3} \pm 2.36 \times 10^{-4} \text{ ms}^{-1}$ |
| $\tau_{stim}$ | $7.73 \times 10^4 \pm 1.69 \times 10^4 \text{ ms}$ |
| $\tau_{DR}$ | $2.46 \times 10^5 \pm 4.26 \times 10^4 \text{ ms}$ |

**Fig 28.**
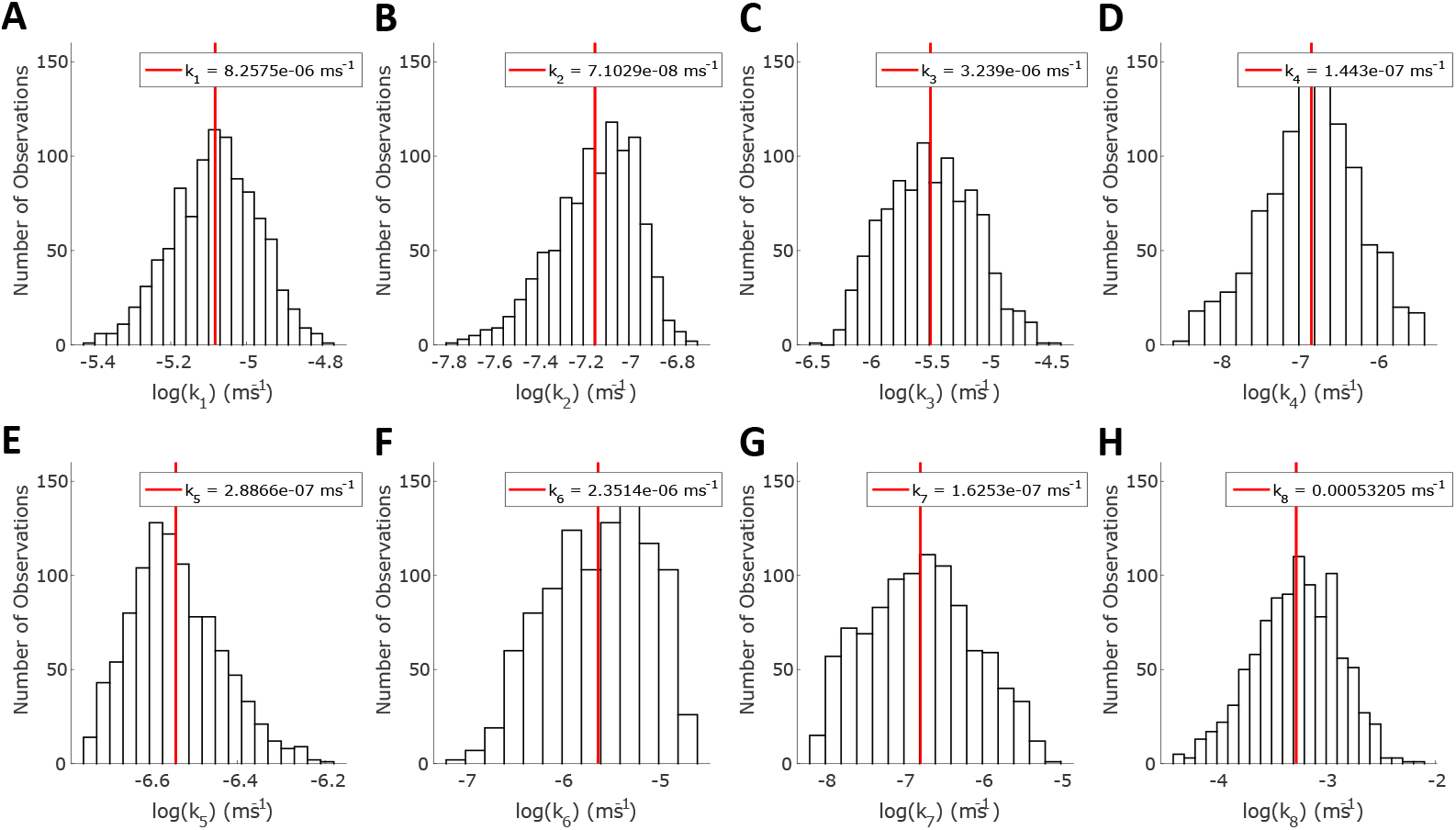
Inference of dopaminergic model parameters. Each histogram represents the approximate posterior distributions of the parameters **(A)** *k*_1_, **(B)** *k*_2_, **(C)** *k*_3_, **(D)** *k*_4_, **(E)** *k*_5_, **(F)** *k*_6_, **(G)** *k*_7_, and **(H)** *k*_8_. The red-line represents the mean log value.

We then inferred the 8 unknown model parameters *k*_*sat*_, *k*_*late*_, *k*_*E*_, *k*_*I*_, *k*_*stim*_, *k*_*DA*_, *τ*_*stim*_, and *τ*_*DR*_ in Eqs. (15i)-(15r) using the ABC-SMC method [33]. For parameter inference, we used the input-output data (% change in fEPSP slope from control by various D1/D5 agonists and HFS/LFS simultaneously) available from rat hippocampal slice experiments [2, 8, 27, 34, 39, 40]. Using our full model to infer the parameters *k*_*sat*_, *k*_*late*_, *k*_*E*_, *k*_*I*_, *k*_*stim*_, *k*_DA_, *τ*_*stim*_, and *τ*_*DA*_ is computationally expensive. Therefore, we considered a reduced a model that consisted of the frequency dependent plasticity model (Eqs. (6a)–(6e)), the dopaminergic model (Eqs. (15a)–(15h)), the temporal dopaminergic-frequency dependent stimulation interaction model (Eqs. (15i)–(15r)), and the characteristic NMDA calcium current for each HFS and LFS protocol. We considered a mean sum of squared errors distance function (Eq. (17)) averaged over the *m* experimental data sets. The distance function measures the squared distance between *i*^*th*^ % change in fEPSP slope (*x*_*j*_[*i*]) of the *j*^*th*^ data set to the corresponding % change in fEPSP slope in the model (*G*(*P* [*t*_*i*_])) at time *t*_*i*_. The model % change in fEPSP slope was determined by mapping the change in the fractional conductance (*P*) to % change in fEPSP slope with Eq. (10). Table 6 shows our inferred parameters value represented in terms of the mean ± standard deviation and Figure 29 shows the histograms representing the approximate posterior distribution for each parameter.

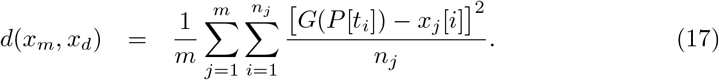

**Fig 29.**
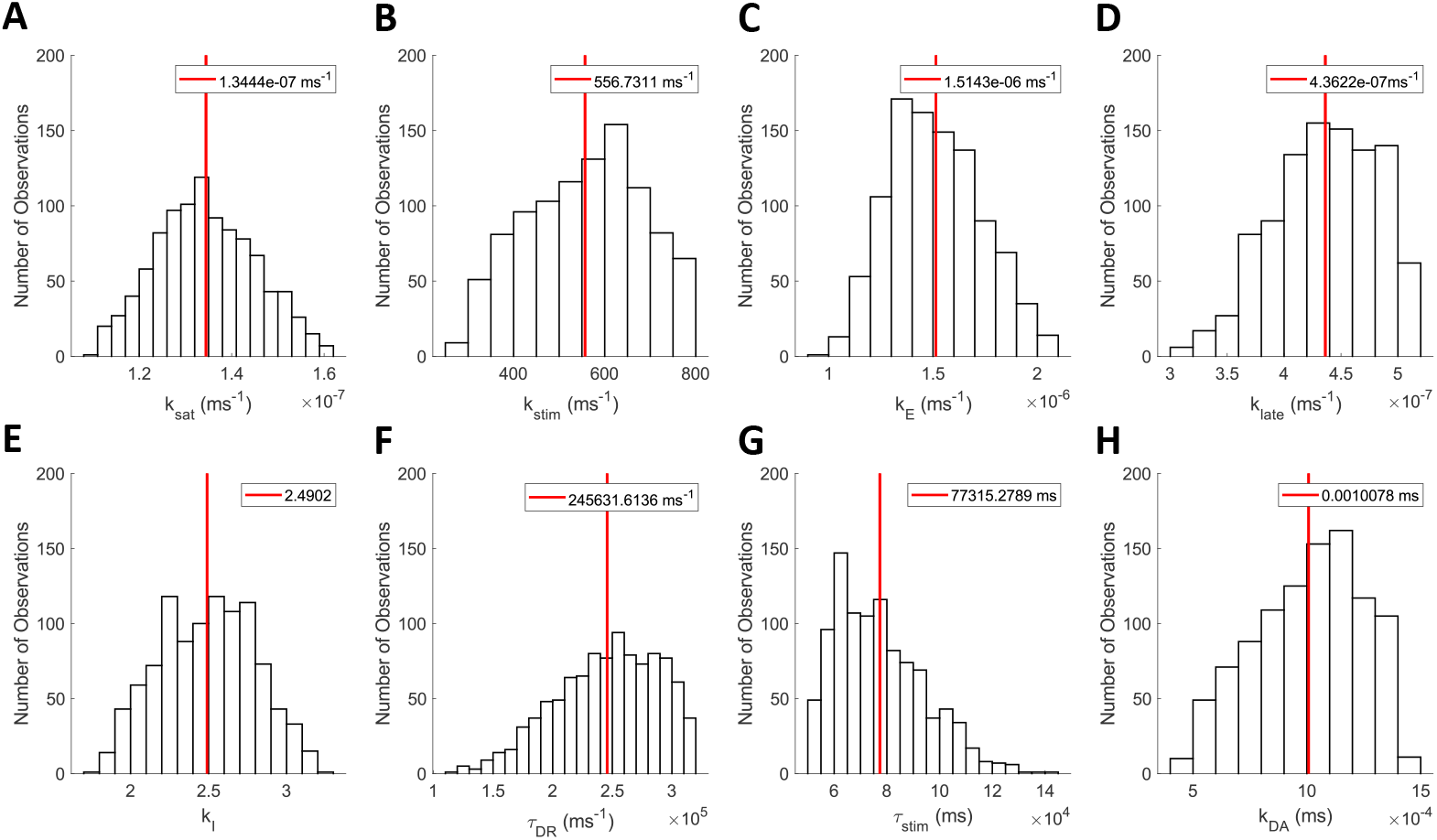
Inference of dopaminergic modulation of HFS and LFS model parameters. Each histogram represents the approximate posterior distributions of the parameters **(A)** *k*_*sat*_, **(B)** *k*_*stim*_, **(C)** *k*_*E*_, **(D)** *k*_*I*_ , **(E)** *k*_*late*_, **(F)** *k*_*DA*_, **(G)** *τ*_*stim*_, and **(H)** *τ*_*DA*_. The red-line represents the mean value.

### The Complete Model

Using Eq. (15o), we modified our synaptic current model shown in Eq. (8) as

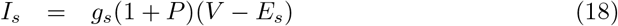

to incorporate the effect of both the HFS/LFS induced changes and dopaminergic induced changes in the synaptic current to the CA1 pyramidal neuron. With this, our complete model describing the modulation of the HFS/LFS induced LTP/LTD by a dopamine agonist in a dose-dependent manner is given by Eq. (1), Eqs. (2a)–(2h), Eqs. (3a)–(3r), Eqs. (4b)–(4h), Eqs. (6a)–(6i), Eqs. (15a)–(15r), and Eq. (18).

### Bayesian Parameter Estimation

We inferred our model parameters using an approximate Bayesian computation sequential Monte Carlo framework [33]. This approach approximates the posterior distribution *π*(*θ|x*) of desired parameters *θ* based on available experimental data *x*_*d*_. The approximate Bayesian computation sequential Monte Carlo algorithm is as follows:

0. Set population indicator to *t* = 0.
1. Set particle indicator to *i* = 1.
2. Draw *θ*\*\* from *π*(*θ*)
3. Simulate *x*\* ~ *f* (*x*|*θ*\*\*) and if *d*(*x*\*, *x*_*d*_) ≥ *ϵ*_*t*_ return to Step 1.
4. Set 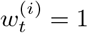.
5. If *i* < *N* return to Step 2 and set *i* = *i* + 1.
6. Normalize weights.
7. Set population indicator to *t* = 1.
8. Set particle indicator to *i* = 1.
9. Sample *θ*\* from previous population 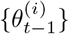 with weights *w*_*t*−1_ and perturb the particle to obtain *θ*\*\* ~ *K*_*t*_(*θ*|*θ*\*). If the *π*(*θ*\*\*) = 0 repeat step.
10. Simulate *x*\* ~ *f*(*x*|*θ*\*\*) and if *d*(*x*\*, *x*_*d*_) ≥ *ϵ*_*t*_ return to Step 9.
11. Set *θ_t_*^(*i*)^ = *θ*\*\* and calculate the weight

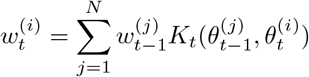
12. If *i* < *N* set *i* = *i* + 1 and return to Step 9.
13. Normalize weights if *t* < *T* and set *t* = *t* + 1, then return to Step 8.

## Acknowledgments

Thank you to Andrew Branen for reviewing and proofreading the article.

## Notes

### Competing Interest Statement

The authors have declared no competing interest.

